# Osmotic conditions shape fitness gains and resistance mechanisms during *E. coli* and T4 phage co-evolution

**DOI:** 10.1101/2025.03.14.643354

**Authors:** Michael Hunter, Eric Lyall, Kriti Verma, Carolina Tropini

## Abstract

Environmental conditions strongly influence interactions between bacteria and bacteriophages (phages). Here, we examined how osmolality (solute concentration) shapes the in vitro co-evolution of T4 phage and its host *Escherichia coli* during serial passage. When evolved independently, we observed substantial fitness gains in both bacteria and phages, particularly in high osmotic conditions. During co-evolution, however, fitness gains were limited, bacterial populations consistently evolved phage resistance, and several phage populations went extinct. Furthermore, the resistance mechanisms varied by osmolality. In lower osmolalities, mutations disrupted phage binding sites, conferring strong resistance. In higher osmolalities, mutations led to increased colonic acid production, producing a mucoid phenotype with weaker resistance. Because mucoidy has been associated with increased bacterial virulence, these findings suggest that gut-relevant osmotic conditions may constrain evolutionary trajectories, favoring resistance strategies that are less effective against phage but potentially more virulent, with important implications for phage therapy design.

**Significance:** Phages offer a promising alternative to antibiotics, but their safety and efficacy strongly depends on the environmental conditions where the bacteria and phages interact. In the human gut, for instance, solute concentrations can vary widely due to factors like food intolerances or laxative use. In this study, we show that such variations significantly impact how bacteria and phages co-evolve. In particular, we find that in higher osmolalities, bacteria evolve phage resistance through mucoidy – a phenotype linked with increased bacterial virulence – rather than receptor loss. This highlights the need to consider environmental factors when developing phage therapies.

## Introduction

Bacteriophages (phages) are viruses that infect bacteria and hold significant promise as an alternative to antibiotics [1–4]. Phages are often highly specific to their host, making them a particularly attractive prospect for applications in the gut, where traditional antibiotics can cause broad collateral damage to the microbiota [5,6]. Given that phages have been found to persist in the mammalian gut for prolonged periods (>1 month), they could potentially be used to suppress pathobionts – organisms that cause disease under certain conditions – which have been implicated in conditions such as inflammatory bowel disease (IBD) [7–11].

However, the successful application of phage therapies in the gut requires an understanding of how intestinal conditions impact phage infection and co-evolution. Unlike in sepsis, where bacteria grow exponentially and phage therapy can result in complete eradication, phage therapies in the gut are more likely to focus on suppression, as bacterial growth rates are typically lower and more variable [12–14]. This variability arises from the dynamic nature of the gut environment, where there are spatial and temporal variations in various physical and biochemical factors [15]. Previous studies have shown that such factors can influence phage stability and infectivity; however, it remains unclear how they impact the co-existence and long-term co-evolution of phages and bacteria [16–20].

Among these factors, osmolality – the concentration of solute particles – plays a critical role in shaping both the gut environment and the bacterial-viral dynamics [21]. Osmolality is determined by the movement of luminal contents and water across the intestinal epithelium, and can fluctuate substantially due to factors such as food intolerances, laxative use, and malabsorption [15, 21]. Bacteria are able to respond to such fluctuations through the accumulation of compatible solutes or changes in membrane potential, yet these changes impact metabolic activity, growth rates, and community composition [21–24]. In turn, this can affect phage adsorption and replication [25–27], as well as phage structure and integrity [28–30]. Therefore, understanding how osmolality impacts bacteria and phage (co-)evolution is critical to the design and application of phage therapies for conditions such as IBD, in which malabsorption is a common co-morbidity.

In this study, we sought to characterize the genetic and phenotypic changes that occur in bacteria and phage populations (co-)evolved under different osmotic conditions. We used *Escherichia coli* and its bacteriophage T4 as a model system for experimental evolution. Both are well studied systems, and the abundance of facultative anaerobes, including *E. coli*, is often increased in the microbiota of people with IBD [31], with several *E. coli* pathobionts being shown to exacerbate gut inflammation [32, 33], and phage-based strategies are being developed to target them [34, 35]. T4 phage follows a strictly lytic lifecycle: it binds to lipopolysaccharide (LPS) and outer membrane protein C (*ompC*) on the bacterial surface, injects its DNA, hijacks the host metabolism to produce more phages, ultimately lysing the cell to release its viral progeny [36, 37].

Crucially, *ompC* expression is upregulated under high osmotic conditions [38], leading us to hypothesize that elevated osmolality would exert distinct selective pressures and shape the adaptation of bacteria and phages. To test this, we evolved *E. coli* and T4 phage independently and in co-culture under three different osmotic conditions, tracking fitness changes and the evolution of phage resistance. During independent evolution, both bacteria and phage became more fit compared to ancestral populations, with the largest fitness increases occurring in populations grown in high-osmotic conditions. In contrast, co-evolution produced very different outcomes: fitness improvements were limited, bacterial populations consistently evolved phage resistance, and several phage populations became extinct. However, the level and mechanism of phage resistance depended on the osmotic conditions. In lower osmolalities, bacteria typically acquired mutations impacting *ompC* and genes associated with LPS (*waaR, waaG*, and *galU*) that conferred strong or complete resistance, whereas high osmolality populations developed a mucoid phenotype that conferred weaker resistance. Together, these findings suggest that osmotic conditions constrain the evolutionary trajectories available during bacteria-phage co-evolution, with important implications for the design of gut phage therapies.

## Results

### Independent evolution and co-evolution lead to distinct evolutionary trajectories

We experimentally evolved *E. coli* and phage T4 in three regimes: (i) *E. coli* alone, (ii) T4 alone, and (iii) *E. coli* and T4 together – see Fig. 1a and Methods. In both independent and co-evolution experiments, ancestral clones were used to seed three replicate populations in each of three osmotic conditions. Populations were evolved by daily serial passage for 36 days under “low”, “medium”, and “high” osmotic conditions (*∼*240, 450, and 920 mOsm/kg H_2_O, respectively). These osmolality levels were chosen to span relevant gut osmotic conditions, from the basal osmolality of the minimal medium (low), to conditions representative of a healthy gut (medium) and a perturbed gut undergoing malabsorption (high) [21].

**Figure 1:**
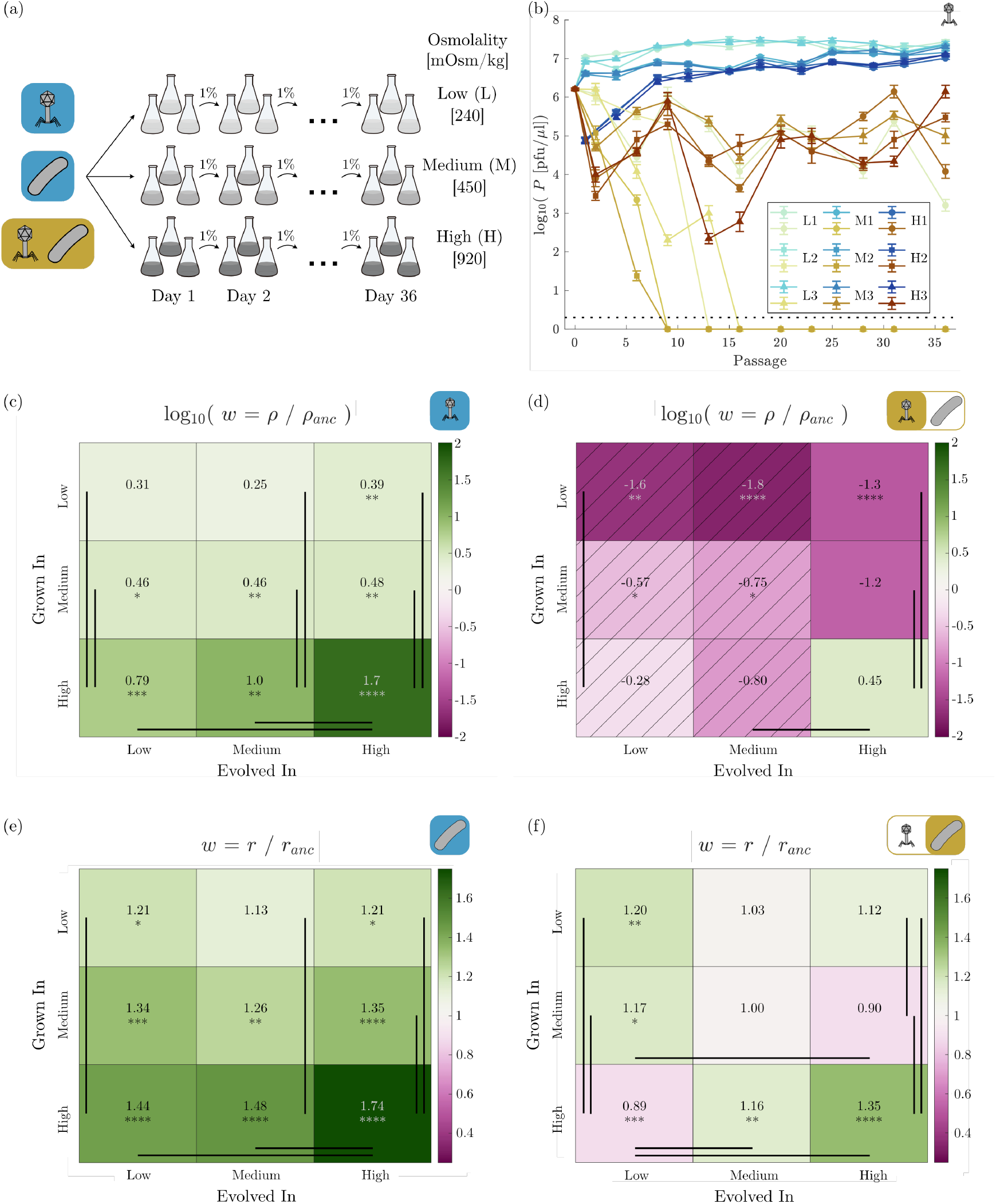
Independent evolution and co-evolution lead to distinct evolutionary trajectories. (a) A schematic of the serial passage evolution experiment. (b) Phage population size *P* throughout the experiment, measured by plaque assay on the ancestral bacteria strain (c, d) The relative fitness *w* of the final independently (c) and co-evolved (d) phage populations, defined as the ratio of their endpoint density *ρ* relative to the endpoint density of the ancestral phage *ρ*_*anc*_ grown in the same conditions. This was determined by measuring the final phage concentration reached given a defined initial culture with the ancestral strain - see Methods. Hatched lines in (d) indicate evolutionary conditions where some lines went extinct prior to the end of the experiment. (e, f) The relative fitness *w* of the final independently (e) and co-evolved (f) bacteria populations, defined as the ratio of their exponential growth rate *r* relative to the exponential growth rate of the ancestral strain *r*_*anc*_ grown in the same conditions. Statistical comparisons between evolved and ancestor populations determined using two-sample t-tests followed by the Bonferroni correction applied as *p*-value adjustment. * *p* < 0.05, ** *p* < 0.01, *** *p* < 0.001, and **** *p* < 0.0001. Statistical comparisons between evolved groupings made using two-way ANOVA followed by Tukey’s honestly significant difference test. Any comparison with *p* < 0.05 is shown using a solid bar. Note that only statistically significant comparisons *within* each “Grown In” or “Evolved in” condition are shown.

During independent phage evolution, all populations showed an overall increase in population size over the 36 days (Fig.1b). In contrast, during the co-evolution experiment, phage population sizes varied far more across both time points and replicate lines, suggesting that reciprocal adaptation between bacteria and phage generated more variable evolutionary dynamics (Fig. 1b). Unlike during independent evolution, no consistent increase in phage population size was observed across osmotic conditions during co-evolution. Indeed, four of the nine co-evolved phage populations went extinct before the end of the experiment (two under low osmolality, two under medium osmolality).

To further investigate the increasing population size during independent evolution, we assessed the relative fitness *w* of the final evolved phage populations. Relative fitness was defined as the ratio of the endpoint density of the evolved phage population, *ρ*, relative to the ancestral phage, *ρ*_*anc*_, grown under the same conditions (*w* = *ρ/ρ*_*anc*_). The endpoint density *ρ* was determined by measuring the final phage concentration when a fixed number of phages were inoculated into a dilute overnight culture of the ancestral strain with fixed density (Methods). Across all conditions, independently evolved phages exhibited higher endpoint densities (*w* > 1) than the ancestor (Fig. 1c), with the gains depending strongly on the assay environment: increases in evolved endpoint density were greatest when phages were assayed in high-osmolality conditions. Within this environment, evolutionary history also played a role, with high-osmolality populations achieving significantly higher relative fitness (log_10_(*w*) = 1.7) than those evolved under low (log_10_(*w*) = 0.79) or medium (log_10_(*w*) = 1.0) osmolality.

By contrast, co-evolved phages generally exhibited a reduced endpoint density (*w* < 1) relative to the ancestor, regardless of assay condition (Fig. 1d). The only exception was the high-osmolality populations assayed in their native environment, which showed a modest increase (log_10_(*w*) = 0.45). These results highlight the contrast between independent evolution, which reliably produced increases in endpoint density, and co-evolution, where fitness gains were limited or even negative and several lines went extinct.

We next investigated the bacterial populations and measured the maximum exponential growth rate *r* in each condition (Methods). Here, relative fitness was defined as *w* = *r/r*_*anc*_. The fitness gains for the independently evolved bacteria showed a similar pattern to those of the independently evolved phages. Across all conditions independently evolved bacteria were fitter than the ancestor (*w* > 1), and the magnitude of the gains was largely dictated by the assay condition, with the larger increases in growth rate observed in higher osmolalities (Fig. 1e). Again, however, within the high-osmolality conditions, evolutionary history appeared to play a role, with high-osmolality populations achieving significantly higher relative fitness (*w* = 1.74 for high vs *w* = 1.44 and *w* = 1.48 for low and medium, respectively). In co-evolved bacteria, changes in *w* were smaller and more variable: most populations exhibited modest increases relative to the ancestor (Fig. 1f), there was no consistent relationship between evolutionary history and assay condition, and some populations even showed decreased relative fitness (*w* < 1). All of the underlying growth curves are shown in Fig. S1. Together, these results indicate that, unlike independent evolution, co-evolution did not produce consistent fitness gains across osmotic conditions, suggesting that reciprocal adaptation constrained or destabilized adaptive trajectories.

### Distinct genetic signatures emerge during independent evolution and coevolution

The results in the previous section showed that independent evolution consistently produced fitness gains in both phages and bacteria, whereas co-evolution constrained such gains and in some cases reduced fitness. We next turned our attention to the genetic basis of the fitness gains, and sequenced all of the final independently and co-evolved bacteria and phage populations (Tables S1–S4).

For each bacterial gene, we estimated the probability of at least one mutation based on observed mutation frequencies, assuming sitewise independence (Methods). Across the independently evolved bacteria populations, the top 10 most mutated genes included the RNA polymerase genes *rpoA, rpoB*, and *rpoC*, along with various intergenic regions (Fig. 2a). We saw that some of these regions were also mutated in the co-evolved bacteria populations, although crucially, *rpoA, rpoB, rpoC*, and the intergenic region between *pyrE* and *rph* were not mutated in any of the co-evolved bacteria populations (Fig. 2b).

**Figure 2:**
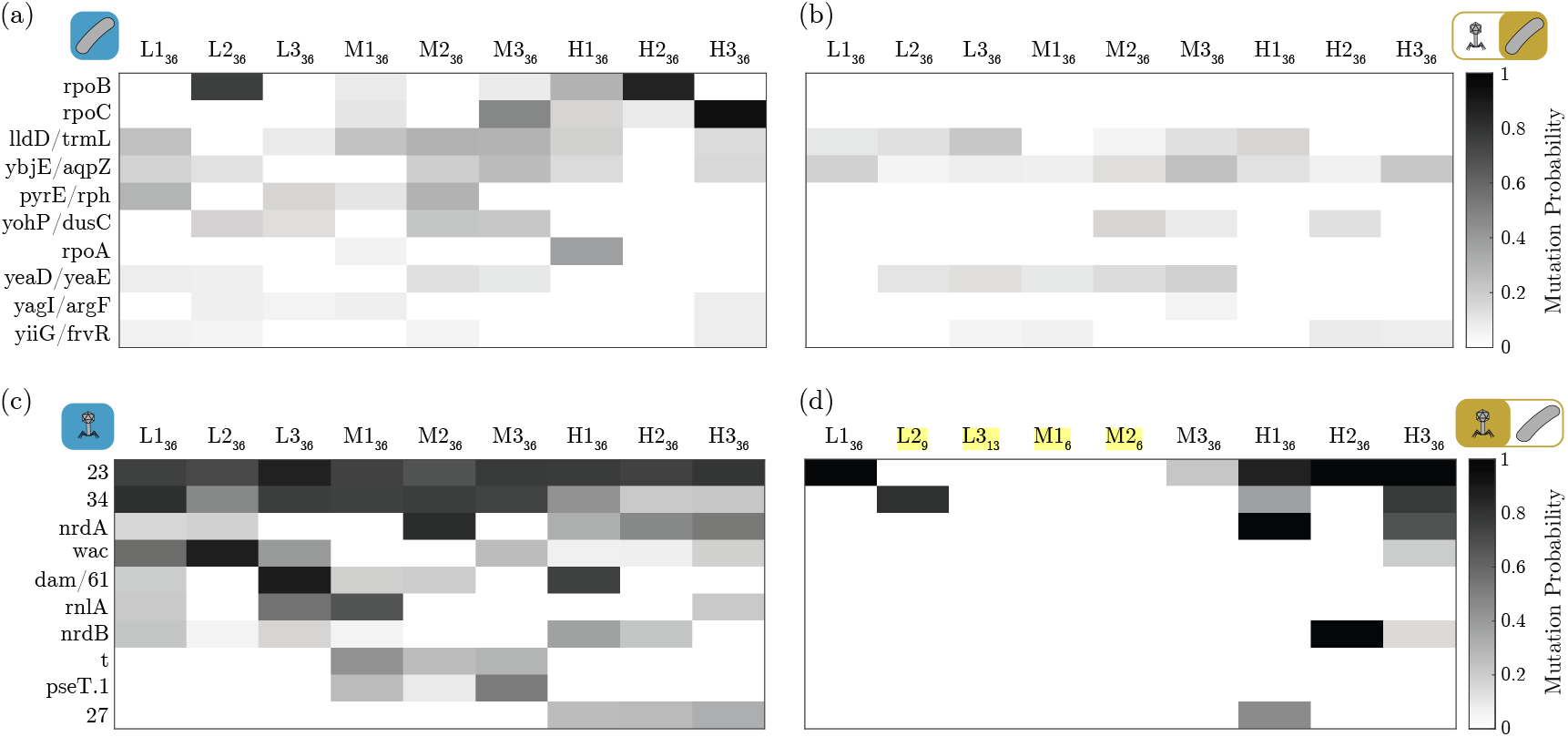
The genetic basis of fitness increases. (a, b) Per gene probability of having at least one mutation (Methods) in the independently (a) and co-evolved (b) bacteria populations. Genes are ranked based on the sum of probabilities across all independently evolved populations, with the highest 10 being shown here. Population subscripts indicate the passage of the sample. Note that here and throughout, expressions incorporating a slash such as “lldD/trmL,” refer to intergenic regions. (c, d) Per gene probability of having at least one mutation in the independently (c) and co-evolved (d) phage populations. Genes are ranked based on the sum of probabilities across all independently evolved populations, with the highest 10 being shown here. Samples taken before the end of the experiment (i.e., passage<36) due to phage extinction are highlighted in yellow.

Several previous studies investigating *E. coli* evolution in minimal media (though not in different osmotic conditions) have reported similar findings, with mutations affecting *rpoB, rpoC*, and the intergenic region between *pyrE* and *rph* having been shown to increase growth rate [39–41]. The specific mutations vary between studies and are different from those that we observe (Table S1 and Table S2); however, it seems likely that this is the primary source of the growth rate increases we observe.

We next applied the same analysis to the phage populations, estimating the per gene probability of having at least one mutation (Methods). Gene *23*, encoding the major capsid protein [37], showed a high probability of mutation in all independently evolved populations (Fig. 2c), as did gene *34*, which encodes the proximal tail fiber subunit [37]. The remainder of the top 10 mutated genes were not mutated in every population, but still had high probabilities of mutation in multiple populations.

Gene *23* was also mutated in all the co-evolved populations that did not go extinct prior to the end of the experiment (Fig. 2d). In the populations co-evolved in high osmolality, we also saw high levels of mutation in some of the other top 10 genes when tested at day 36: gene *34* in H1_36_ and H3_36_; *nrdA*, encoding the ribonucleotide reductase *α* subunit [37], in H1_36_ and H3_36_; and *nrdB*, encoding the ribonucleotide reductase *β* subunit [37], in H2_36_. However, outside gene *23*, and with the exception of gene *34* in one of the lines that went extinct (L2_9_), none of the low or medium osmolality populations had mutations in the top 10 genes identified from the independently evolved populations.

To investigate the mechanistic basis of the large increases in phage endpoint density, we measured several key life-history parameters of the evolved populations. One-step growth curves and adsorption rate assays were used to estimate burst size, lysis time, and adsorption rate, and a mathematical model was used to predict how changes in these parameters would influence endpoint density. Although some changes were observed, relative to the ancestral phage, the magnitude of the changes was small, and model predictions indicated that they were insufficient to account for the changes in endpoint density we observed. Together, these results suggest that the fitness gains observed during independent phage evolution cannot be readily explained by changes in canonical life-history parameters alone. Further details of these analyses can be found in Section S2 of the Supplementary Information.

### Environmental osmolality shapes phage-resistance phenotypes during coevolution

Given that phages went extinct in several of our co-evolved lines, we quantified phage resistance levels in the co-evolved bacteria populations by measuring the efficiency of plating (EOP) of the ancestral phage on each population (Fig. 3a, Methods). Briefly, EOP assays measure the relative ability of a phage to form plaques on different bacterial strains, calculated as the ratio of plaques formed on the test bacterial population relative to the ancestral strain. All co-evolved populations showed EOP values below 1, indicating some level of phage resistance, even in populations where phage extinction did not occur (Fig. 3b). However, resistance levels varied according to evolutionary history: populations evolved under high-osmotic conditions displayed lower resistance than those evolved under low- and medium-osmotic conditions. Indeed, in the low- and medium-osmolality populations, no susceptibility to the ancestral phage was detected (detection limit: ∼EOP 2.5× 10^*−*^_7_), indicating the evolution of effectively complete resistance in these conditions. By contrast, the persistence of measurable susceptibility in the high-osmolality populations suggested that different, less protective resistance mechanisms had evolved.

**Figure 3:**
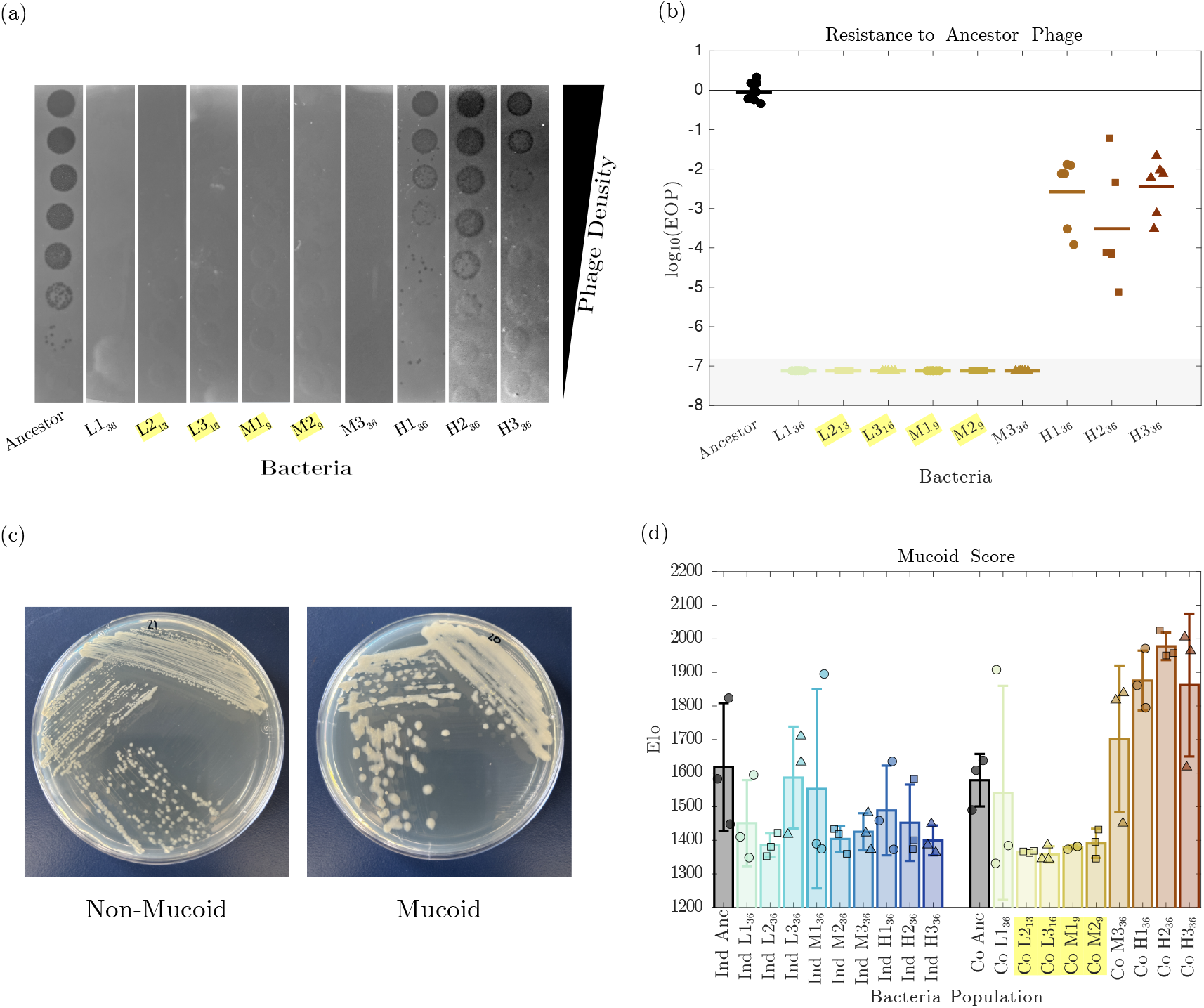
Environmental osmolality drives distinct resistance phenotypes during co-evolution. (a) Images of efficiency of plating (EOP) assays. The same sample of ancestral phage was serially diluted in tenfold increments and spot-plated onto the ancestral strain and each of the co-evolved bacteria populations. Subscripts in population labels indicate passage. Samples taken before the end of the experiment (i.e., passage<36) due to phage extinction are highlighted in yellow. (b) EOP of the ancestral phage on each of the co-evolved bacteria populations immediately following phage extinction, or at the end of the experiment if no phage extinction occurred. The gray shaded region indicates below the limit of detection – no plaques were observed on any low- or medium-osmolality bacterial populations. (c) Representative images of the mucoid phenotype exhibited by several bacterial populations. (d) Degree of mucoidy in each of the independently evolved (Ind) and co-evolved (Co) bacteria populations, quantified by an Elo score (Methods).

When streaking out the co-evolved bacteria populations, we also observed a distinct ‘mucoid’ colony morphology in some of the populations (Fig. 3c). To explore this further, we systematically streaked all the independently and co-evolved bacteria populations and quantified the level of mucoidy (Methods). Briefly, we compared pairs of bacterial plates, blind to their evolutionary history, to judge which looked more mucoid. These comparisons were then used to calculate Elo scores, which reflected the relative mucoidy level of each population. Most populations had similar or lower average Elo scores compared to the ancestral strain, indicating that they are not mucoid (Fig. 3d). However, co-evolved populations M3, H1, H2, and H3 all showed higher average Elo scores than the ancestral strain. This pattern matched our initial qualitative observations.

### Distinct genetic mechanisms lead to phage resistance in co-evolved bacteria

To investigate the genetic basis of the differences in phage resistance and colony morphology, we turned to our sequencing of the co-evolved bacteria populations. Looking across all mutations in the genome (Fig. 4a), and again estimating the per gene probability of mutation (Fig. 4b), a clear pattern emerged: low- and medium-osmolality populations consistently acquired mutations in the phage receptor gene *ompC* and *rrsG* (encoding one of the 16S ribosomal RNA genes [42]), while high-osmolality populations acquired mutations affecting the regulator of capsule synthesis (RCS) pathway.

**Figure 4:**
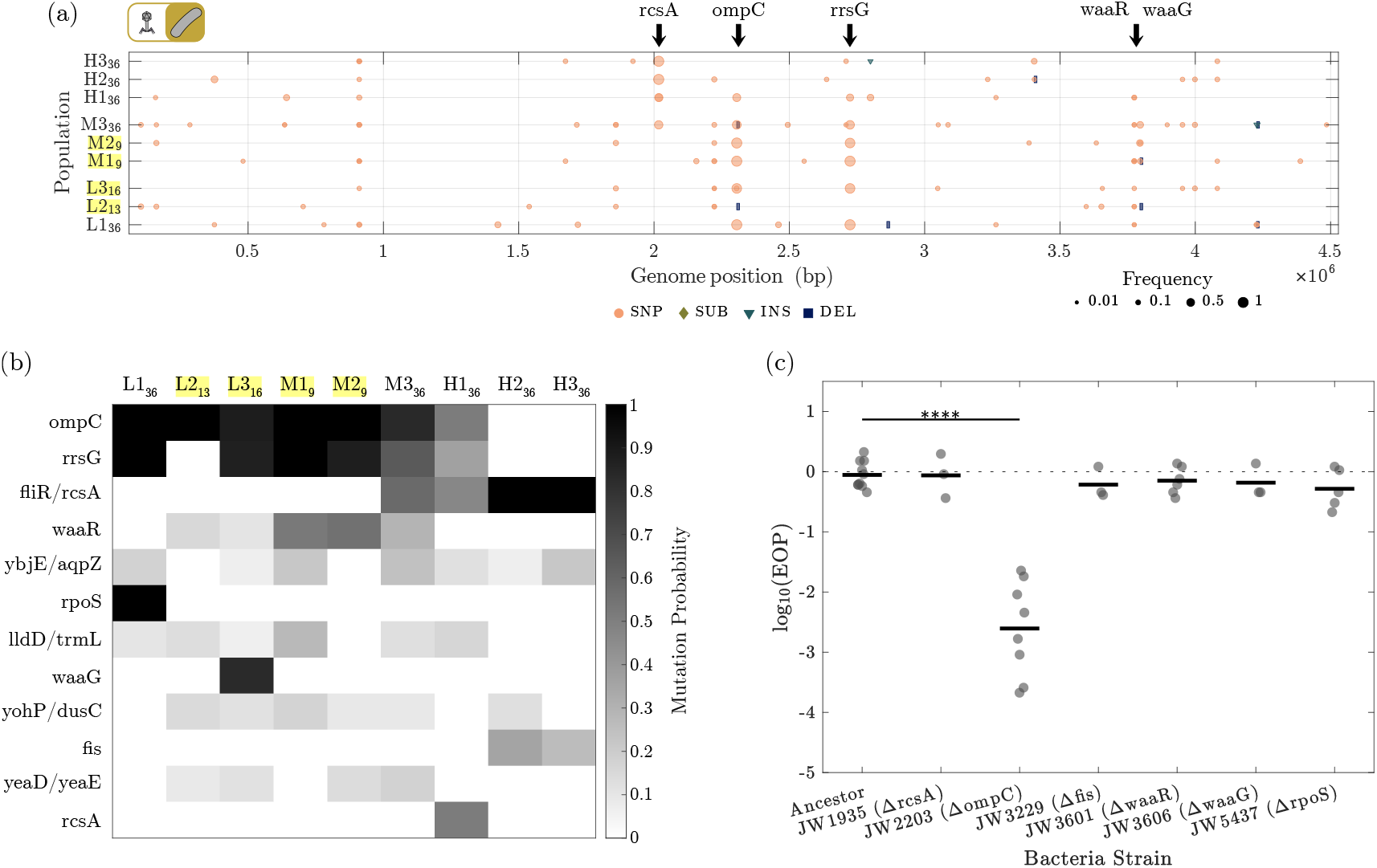
Population sequencing reveals distinct genetic bases of phage resistance. (a) Mutations plots of the co-evolved bacteria populations. Deletion bars are shown at a fixed minimum size for visibility. Labels show annotations for key genes discussed in the text. Population subscripts indicate the passage of the sample. Samples taken before the end of the experiment (i.e., passage<36) due to phage extinction are highlighted in yellow. (b) Per gene probability of having at least one mutation (Methods) in the co-evolved bacteria populations. Genes are ranked based on the sum of probabilities across all co-evolved populations, with the highest 12 being shown here. (c) EOP of the ancestral phage on various Keio knockout strains [43]. Statistical significance determined from one-way ANOVA followed by Dunnett’s test with the Ancestor as control. **** *p* < 0.0001.

Populations evolved under high-osmotic conditions consistently acquired mutations in *rcsA* or its promoter region (*fliR/rcsA*), while this gene was mutated in only one of the medium-osmolality populations, and none of the low-osmolality ones. The RCS pathway plays a critical role in sensing membrane stresses, including outer membrane damage, defects in lipopolysaccharide (LPS) synthesis, mechanical compression, and osmotic shock, while also regulating exopolysaccharide production [44]. Overproduction of exopolysaccharides via activation of this pathway results in the distinct ‘mucoid’ colony morphology we observed, characterized by an excess polysaccharide layer coating the cell surface [45]. Although the precise mutations we observe (Table S2) are distinct, these findings are consistent with previous studies demonstrating that this phenotype provides a generalized phage resistance mechanism by limiting access to receptor sites on the bacterial cell surface [46, 47].

In contrast, all low-and medium-osmolality populations had mutations in the *ompC* gene, a known T4 phage binding site, while only one high-osmolality population had mutations in this gene. A single mutation (R195P) dominated among *ompC* mutations and was highly correlated with a specific mutation in *rrsG* (Table S2, Fig. S6). To test the effect of *ompC* inactivation on phage infection, we examined phage resistance in an *ompC* knockout strain (JW2203) from the Keio collection [43]. As expected, the *ompC* knockout showed some resistance to the ancestral phage (Fig. 4c). Interestingly, however, while our co-evolved populations with *ompC* mutations were fully resistant to the ancestral phage (Fig. 3b), the *ompC* knockout strain was only partially resistant.

This may suggest that *ompC* mutations act in combination with additional genomic changes, such as the observed mutations in *rrsG*, to generate full resistance, even though inactivation of the other highly mutated genes does not independently confer resistance to the ancestral phage (Fig. 4c). To explore this possibility further, we isolated five clones from each co-evolved population, sequenced them, and quantified their resistance to the ancestral phage.

The isolates fell into three broad phenotypic classes: complete resistance, high/intermediate resistance, and low resistance (Fig. 5a). Populations L2, L3, M1, and M2 consisted entirely of fully resistant isolates, without detectable plaques. All L1 isolates showed intermediate resistance −log_10_(EOP) ∼−4. Two populations were heterogeneous: M3 contained a mix of highly resistant −log_10_(EOP) ≲ − 5 – and fully resistant clones, and H1 contained a mix of highly resistant and low-resistance clones – log_10_(EOP) ∼−1. In contrast, all isolates from H2 and H3 showed uniformly low resistance.

**Figure 5:**
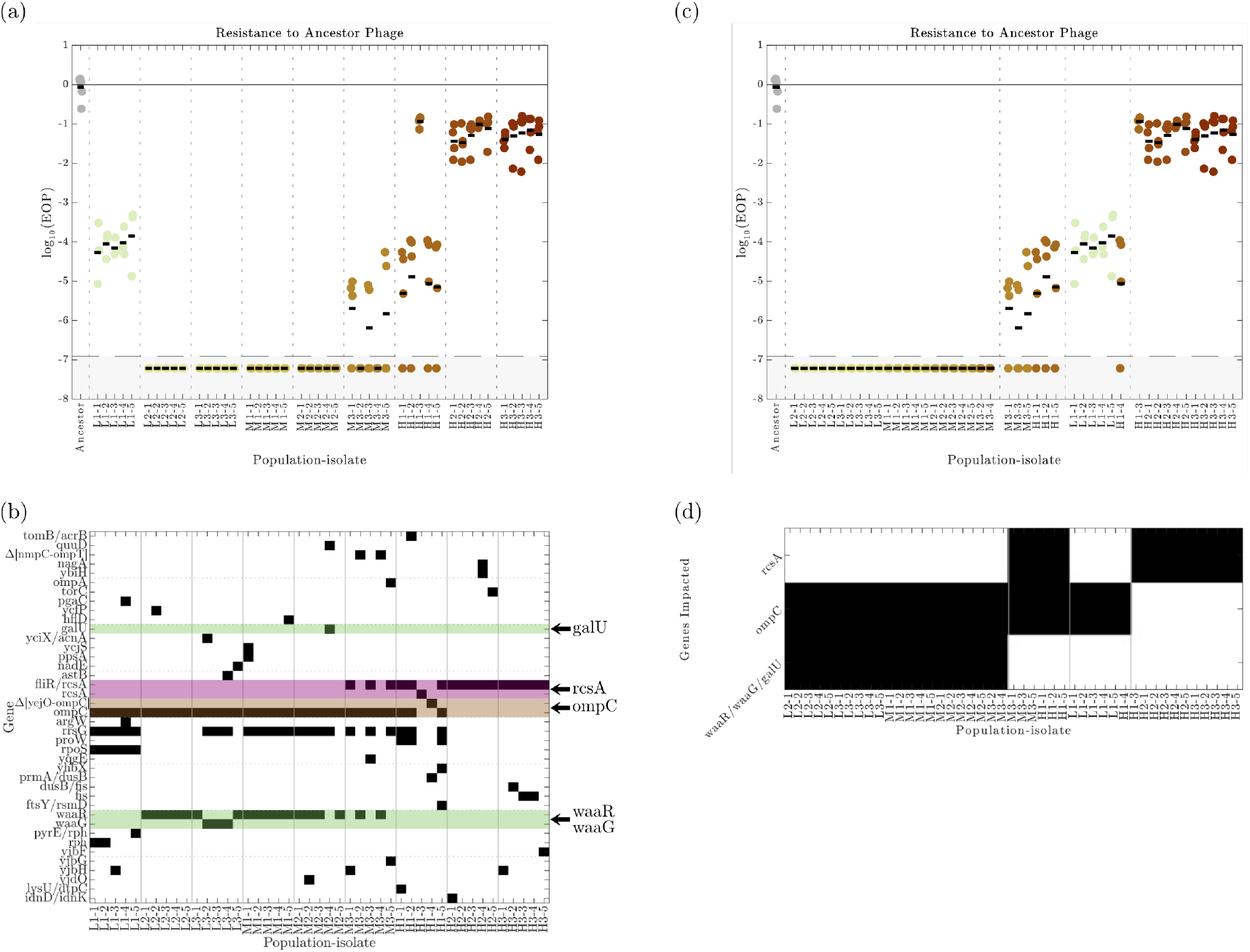
Isolate analyses reveal the genetic origins of resistance phenotypes. (a) EOP of the ancestral phage on each of the isolates from the co-evolved bacteria populations. The gray shaded region indicates below the limit of detection - points in this region indicate that no plaques were observed. (b) Binary matrix showing which genes are mutated (black) in each bacterial isolate. Large deletions affecting multiple genes are indicated as Δ[*geneA*-*geneB* ] with all genes between *geneA* and *geneB* (inclusive) being affected. (c, d) Same as (a, b) but with different sorting to highlight the link between the resistance phenotype (c) and underlying mutations (d).

These phenotypes correspond closely to the underlying mutations (Fig. 5b and Table S5). All isolates with complete resistance carried mutations in both *ompC* and genes affecting LPS core structure (*waaR, waaG*, or *galU*). Isolates with high, though not complete, levels of resistance carried *ompC* mutations and mutations affecting *rcsA*, but not affecting LPS. Isolates with intermediate levels of resistance, comparable to the *ompC* knockout strain (Fig. 4c), carried *ompC* mutations, but no mutations affecting LPS or *rcsA*. Finally, isolates with low levels of resistance carried mutations affecting *rcsA*, but not *ompC* or LPS. All isolates fell into one of these four groups, highlighting a clear link between resistance phenotypes (Fig. 5c) and underlying genetic changes (Fig. 5d).

### Diverse mutations target common receptor-binding functions in co-evolved phages

Finally, we wanted to explore whether co-evolved phages had developed adaptations to counter evolution in their respective bacterial hosts, and whether these adaptations were environment-specific or broadly conserved.

Focusing first on the sequencing results from these populations (Fig 6a), we found that even among high-frequency mutations (defined as *f* > 0.7), most were unique to a single co-evolved phage population (Fig 6b), suggesting that co-evolution generated diverse adaptive trajectories across replicate lines. Populations that went extinct averaged 1.25 high-frequency mutations, standard deviation *σ* = 0.96, while populations that persisted averaged 4.80, *σ* = 1.64, (Fig 6c). However, many of these mutations occurred in the same genes across different populations (Fig 6d), in particular in gene *37*, which encodes the large distal tail fiber subunit, and in genes *10* and *7*, which both encode baseplate wedge components [37].

**Figure 6:**
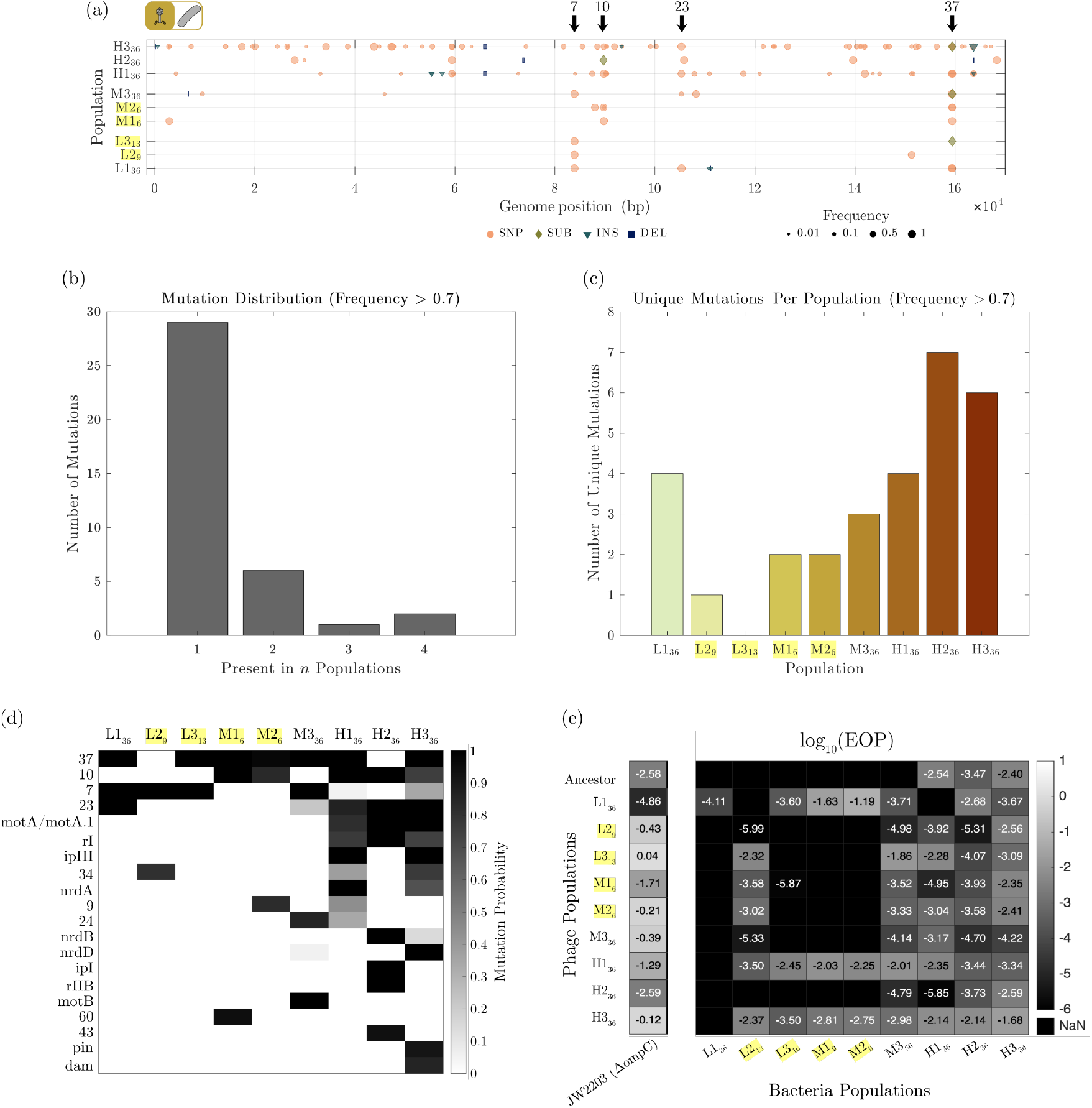
Diverse mutations target common receptor-binding functions in co-evolved phages. (a) Mutations plots of the co-evolved phage populations. Length of the deletion bars represent the length of the deletion, unless the deletions are less than 100 bp, in which case they are shown as 100 bp for visibility. Labels show annotations for key genes discussed in the text. Population subscripts indicate the passage of the sample. Samples taken before the end of the experiment (i.e., passage<36) due to phage extinction are highlighted in yellow. (b) The number of mutations with frequency *f* > 0.7 present in *n* co-evolved phage populations. (c) The number of unique mutations with frequency *f* > 0.7 present in each co-evolved phage population. (d) Per gene probability of having at least one mutation in the co-evolved phage populations. Genes are ranked based on the sum of probabilities across all co-evolved populations, with the highest 20 being shown here (e) EOP of all co-evolved phage populations on the *ompC* knockout strain and all co-evolved bacteria populations. Note that ‘missing’ values (black tiles) mean that no plaques were visible in any replicates for this combination of phage and bacteria. If plaques were visible in some replicates and not others, any missing titres were conservatively assumed to equal half the limit of detection.

To evaluate the specificity and breadth of resistance and counter-resistance adaptations, we tested the susceptibility of each co-evolved bacterial population to all co-evolved phage populations (Fig 6e). This allowed us to determine whether phage resistance and infectivity were restricted to their co-evolved counterparts, shared among populations evolved under similar osmotic conditions, or generalized across populations from all conditions.

We found that bacteria with mucoid mutations (M3, H1, H2, and H3) were partially susceptible to nearly all co-evolved phage populations, regardless of the conditions in which the phages evolved. In contrast, bacterial populations carrying mutations affecting *ompC* and LPS generally exhibited broad resistance, remaining fully resistant to most co-evolved phages. One exception to this was the L2 bacteria population, which showed partial susceptibility to several co-evolved phage populations. Interestingly, this population also carries a distinct *ompC* mutation: a 6 bp deletion, compared to the R195P SNP seen in other populations (Table S2), potentially explaining the observed variation in phage resistance.

Furthermore, despite the robust resistance observed in bacteria with *ompC* mutations, the *ompC* knockout strain displayed little or no resistance to many of the co-evolved phage populations. This is consistent with our analysis of individual isolates, which revealed a high prevalence of mutations also affecting LPS in these populations (Fig. 5). Phage populations L1, H1, and H3, were able to infect the majority of bacteria carrying *ompC* and LPS-related mutations, suggesting the presence of key adaptive mutations in these phages that enable binding to altered or other receptor sites.

## Discussion

In this study, we investigated how environmental osmolality shapes the independent evolution and co-evolution of *E. coli* and T4 phage. Independent evolution produced substantial and repeatable fitness gains in both bacteria and phage, particularly under high-osmolality conditions. In contrast, fitness gains during co-evolution were limited, phage resistance emerged, and several phage populations went extinct. Together, these findings indicate that environmental osmolality not only influences adaptation, but also constrains the evolutionary dynamics of bacteria-phage interactions.

A key finding from the co-evolution experiments was that environmental osmolality strongly shapes the evolution of phage-resistance. Specifically, mutations affecting *ompC* and LPS dominated in low and medium osmolality, whereas mutations affecting *rcsA* dominated in high osmolality, leading to a mucoid phenotype, despite this resulting in lower phage resistance. This pattern suggests that resistance mechanisms are not selected solely based on their effectiveness against phage but are constrained by abiotic conditions.

Two non-mutually exclusive explanations are consistent with this interpretation. First, loss or modification of *ompC* and/or LPS may be deleterious at high osmolality, particularly given that *ompC* is typically upregulated under osmotic stress [38]. Similar trade-offs have previously been documented when phages use efflux pumps as receptor sites and co-evolution occurs in the presence of antibiotics [48, 49]. Second, the mucoid phenotype itself may confer advantages under osmotic stress, as exopolysaccharides such as colonic acid can stabilize the cell envelope and improve osmotic stress tolerance [50, 51]. Consistent with these interpretations, populations exhibited environment-dependent fitness effects, although these measurements cannot be directly attributed to specific resistance mechanisms because they reflect the combined effects of multiple genetic and phenotypic changes.

Supporting this, analysis of multiple clones revealed that fully resistant isolates carry both *ompC* mutations and mutations in genes associated with LPS. The latter were often at low frequency (10-20%) and spread across multiple alleles, obscuring their contribution in population-level sequencing (Table S2). These findings are consistent with previous work showing that T4 plaque formation is reduced in *E. coli ompC* deletion mutants, but completely prevented in mutants lacking both *ompC* and the same LPS-associated genes identified here: *waaR, waaG*, and *galU* [52, 53].

*Furthermore, resistance mechanisms were not strictly mutually exclusive. Some isolates carried mutations affecting both ompC* and *rcsA*, and exhibited higher resistance than isolates carrying either mutation alone, suggesting that high-level resistance can emerge from combinations of mutations affecting multiple components of the cell envelope.

While measurements at the population and isolate-levels were qualitatively consistent, multiple isolates (19/45) exhibited less resistance than their source populations. Several factors may contribute to this. First, continued evolution during the isolation process, given that we observe mutations in some isolates that were not present in the population level measurements. Second, phenotypic state differences between isolates and populations: all discrepancies occurred in populations where phages persisted to the end of the experiment, where ongoing exposure to phages and lysed bacteria may stimulate stress responses, alter growth dynamics, and contribute to phage resistance [46, 54, 55]. Finally, some populations (e.g., M3, H1) contained multiple genotypes that contribute to a collective resistance in ways not captured by single isolates. These results again highlight the importance of considering population heterogeneity when interpreting resistance phenotypes.

In contrast, independent evolution of bacteria and phages followed more consistent evolutionary trajectories, where fitness gains were universal and driven primarily by the growth conditions rather than the evolutionary history. In bacteria, the dominant mutations observed (affecting *rpoB, rpoC*, and the *pyrE* /*rph* intergenic region) are commonly associated with improvements in growth rate in minimal media [39–41], suggesting that bacterial evolution was dominated by adaptation to the media [56]. Similarly, independently evolved phages showed strong genetic convergence with all populations acquiring mutations to a single codon in gene *23*. While mutations in this gene have previously been linked to altered capsid assembly [57–59], our analysis of phage life-history traits (including adsorption rate, burst size, lysis time, and stability) did not identify changes likely to explain the observed fitness gains (Supplementary Information S2).

Furthermore, the largest fitness gains occurred in the high osmolality conditions, where both the ancestral strain and ancestral phage exhibited the lowest growth rates and endpoint densities. This pattern is consistent with previous work showing that less well adapted genotypes exhibit larger fitness gains because the beneficial tail of the distribution of fitness effects is broader, whereas better adapted genotypes have fewer and weaker beneficial mutations available [60, 61].

In contrast to the convergent patterns observed during independent phage evolution, co-evolved phage populations exhibited substantially greater genetic variation. During co-evolution, bacterial resistance evolved over time, continuously shifting the selective pressures experienced by the phages. As a result, selection was less likely to favor a single optimal adaptive strategy across replicate populations. Nevertheless, most populations had mutations impacting gene *37*, which encodes the large distal tail fiber subunit (Table S4) [37]. Specifically, all observed gene *37* mutations occurred in or near the receptor-binding “tip” subdomain [62], where mutations have previously been shown to alter binding to *ompC* and/or LPS [52, 62], alter adsorption kinetics [63, 64], and expand host range [65]. Notably, mutations affecting residue Y953, which arose in multiple populations, have been shown to allow T4 to infect *E. coli* lacking both *ompC* and *waaR* [52].

As a whole, these results are relevant to therapeutic applications, as pre-evolving phages independently under high osmolality could generate variants with increased infectivity, whereas co-evolution with bacteria may select for phages better able to overcome evolving resistance mechanisms. Such ‘phage training’ has previously been shown to expand host range, improve infectivity, counter phage resistance, and target biofilms [66, 67]. Whether phages optimized for growth through independent evolution, or for overcoming resistance through co-evolution, or a combination of the two, would perform better in therapeutic settings remains an open question.

More broadly, our findings suggest that environmental conditions may shape not only the evolution of phage resistance, but also the broader physiological consequences associated with those resistance strategies. In our high-osmolality populations, resistance involved mutations affecting the RCS pathway and the emergence of a mucoid phenotype which has previously been linked to increased virulence, immune evasion, and biofilm formation in several pathogens [44,51,68–70]. These results raise the possibility that environmental conditions promoting phage resistance could also favor traits associated with bacterial persistence or virulence, particularly in fluctuating environments such as the gut, where osmotic perturbations are common [71]. Understanding how different environmental conditions influence the evolutionary trade-offs associated with phage resistance will therefore be important for the design of phage therapies.

Our findings offer insights into how osmotic conditions shape bacterial and phage evolution and highlight the importance of environmental context in bacteria-phage co-evolution. An important next step will be to determine how these dynamics manifest in more complex and physiologically relevant settings. Incorporating features characteristic of natural environments – such as spatial structure, immune interactions, and microbial diversity – could yield additional insights into evolutionary dynamics. In particular, in vivo gut models may help clarify how evolutionary patterns observed here manifest within hosts. Additionally, examining interactions between osmotic stress and other biologically relevant factors, such as fluctuating pH, nutrient availability, or host mucus interactions, will further enhance our understanding of how environmental pressures collectively shape microbial evolution.

In summary, our findings highlight the complex interplay between environmental conditions, bacterial adaptation, and phage evolution. More broadly, they suggest that environmental conditions can constrain the evolutionary trajectories available during bacteria-phage co-evolution, with important implications for phage therapy design. To optimize phage therapy, it will be essential to replicate relevant environmental conditions during phage selection to ensure broad-spectrum efficacy while minimizing the risk of unintentionally selecting for traits that enhance bacterial persistence or virulence. Together, these results emphasize that environmental context should be considered an active evolutionary variable in the development and application of phage therapies.

## Materials and Methods

### Data availability

Genomic sequence data are available at Sequence Read Archive (SRA) BioProject PRJNA1463809.

### Strains

This study used *Escherichia coli* K-12 BW25113 (CGSC#7636) and the obligate lytic bacteriophage T4. Single knockout strains JW1935 (Δ*rcsA*, CGSC#9610), JW2203 (Δ*ompC*, CGSC#9781), JW3229 (Δ*fis*, CGSC#10443), JW3601 (Δ*waaR*, CGSC#10651), JW3606 (Δ*waaG*, CGSC#10655), and JW5437 (Δ*rpoS*, CGSC#11387) from the Keio collection were also used [43].

### Ancestor populations

The bacteria ancestor clones used to initiate experimentally evolved lines were obtained by streaking 25% glycerol freezer stocks onto lysogeny broth (LB, Fisher BioReagents, Miller Formula) and 15 g/L agar (Fisher BioReagents) plates, and incubating overnight at 37°C. Single colonies were then picked and used to inoculate fresh liquid LB cultures which were incubated overnight. Unless otherwise stated, all incubations were done at 37°C, and with shaking at 200 RPM in the case of liquid cultures. These cultures represent the ancestor populations, and were stored as stocks of 25% glycerol at − 80°C.

The T4 ancestor clone used to initiate experimentally evolved lines was obtained by plating 20 *µ*L of serially diluted phage lysate, 200 *µ*L of overnight BW25113 culture, and 10 mL of 5 g/L LB-agar. After overnight incubation, a single plaque was isolated and inoculated into a liquid LB culture of exponentially growing BW25113. After incubation overnight, this culture was then centrifuged at 8,000 g for 5 mins to pellet any remaining bacteria and cellular debris, and the supernatant was stored at 4°C.

### Serial passage experiments

Experiments were conducted under three different osmotic conditions using M9 minimal media (Table 1) supplemented with sodium chloride (NaCl) to achieve osmolalities of 240, 450, and 920 mOsm/kg H_2_O. Osmolality was measured using an Advanced Instruments Osmo1 Single-Sample Micro-Osmometer. NaCl is commonly used to induce osmotic stress in microbial studies because it allows precise control of osmolality [24,56] and is relevant to the gut, where osmotic shifts can result from fluctuations in sodium and chloride ion absorption [72]. Each experiment (independentlyevolved bacteria, independently-evolved phage, and co-evolved bacteria and phage) consisted of three replicate populations per osmotic condition passaged daily for 36 days.

**Table 1:**
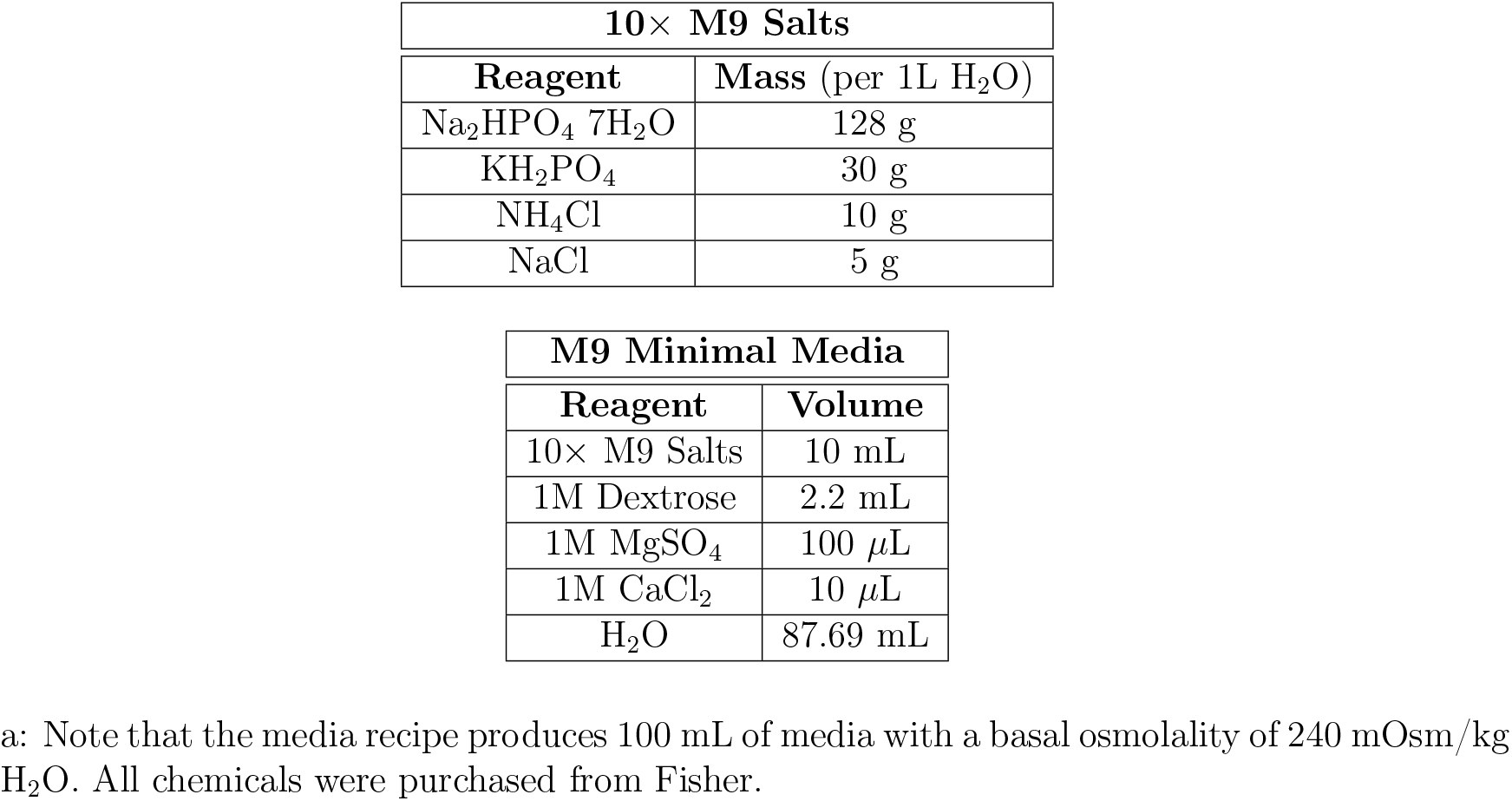
Recipe for M9 minimal media.^*a*^.

| 10× M9 Salts |  |
| --- | --- |
| Reagent | Mass (per 1L H<sub>2</sub>O) |
| Na<sub>2</sub>HPO<sub>4</sub> 7H<sub>2</sub>O | 128 g |
| KH<sub>2</sub>PO<sub>4</sub> | 30 g |
| NH<sub>4</sub>Cl | 10 g |
| NaCl | 5 g |

| M9 Minimal Media |  |
| --- | --- |
| Reagent | Volume |
| 10× M9 Salts | 10 mL |
| 1M Dextrose | 2.2 mL |
| 1M MgSO<sub>4</sub> | 100 μL |
| 1M CaCl<sub>2</sub> | 10 μL |
| H<sub>2</sub>O | 87.69 mL |

a: Note that the media recipe produces 100 mL of media with a basal osmolality of 240 mOsm/kg H<sub>2</sub>O. All chemicals were purchased from Fisher.

In the case of the independently-evolved bacteria, the experiment was initiated by inoculating 2 mL of fresh media with 20 *µ*L of the previously described ancestor culture. Each day, 1% of each population was passaged into 2 mL of fresh media, with samples collected every 3-4 days and stored as 25% glycerol stocks at −80°C.

In the case of the independently-evolved phages, the experiment was initiated by first growing the ancestral *E. coli* strain overnight under each osmotic condition, followed by a 1:50 subculture into 2 mL of fresh media and incubation for 4 hours. Then, 20 *µ*L of T4 ancestor population was added and incubated overnight. Each day, fresh sub-cultures of the ancestral *E. coli* strain were prepared in the same manner, and 1% of the previous day’s phage lysates were passaged into them. To limit bacteria carryover, lysates were centrifuged at 8,000 g for 5 min before transfer, and only the supernatant was used. Samples of the supernatant were taken every 3-4 days and stored at 4°C.

To assess the effectiveness of this procedure in limiting bacteria carryover, we spot-plated the final phage samples after the evolution experiment concluded, and observed no colonies. Because these samples had been stored at 4°C for a prolonged period, we considered that any bacteria carried over may have died during storage. We therefore passaged the final phage samples into the ancestral strain for one additional day using the same centrifugation protocol and spot-plated the resulting supernatant immediately. Again, no colonies were observed. While this suggests bacteria carryover was negligible in these samples, we note that centrifugation may not completely eliminate the possibility of minor carryover.

We note that during the initial independent phage evolution experiment, all three high-osmolality populations experienced a sudden population drop between passages 11 and 15 (Fig. S7), likely due either to the selection of a resistant bacterial colony or to a procedural error during passaging. To confirm this, the lines were restarted from passage 11 samples collected prior to the population drops, and no subsequent drops were observed the second time around, suggesting our hypothesis was correct. The re-run lines were therefore used for the main analyses, while the original lines are included in the Supplementary Information (Fig. S7), where they show qualitatively similar results. In the case of the co-evolved bacteria and phages, the experiment was initiated in the same manner as the independently evolved phage experiment: the *E. coli* ancestral strain was grown overnight, sub-cultured, and incubated for 4 hours before inoculation with the T4 ancestor. Thereafter, 1% of the co-culture was passaged into 2 mL of fresh media each day. For sample storage, every 3-4 days cultures were centrifuged to pellet bacteria and the supernatants were stored at 4°C as ‘phage-only’ samples. The bacteria pellets were then resuspended in fresh media before storage as 25% glycerol stocks at 80°C. While supernatant removal eliminated most phages, some remained in the bacterial stocks (Fig. S8).

To obtain clonal isolates from the co-evolved bacteria populations, the frozen 25% glycerol stocks were streaked onto 1.5% LB-agar plates and incubated overnight at 37°C. Five single colonies were then picked from each population and restreaked twice more for isolation. Finally, single colonies were picked and grown overnight in LB, then frozen at –80°C in 25% glycerol.

### Bacterial growth curves

Bacterial growth curves were performed to measure the growth rate and maximum density reached by a given bacteria population in a given growth condition. These measurements were performed by scraping bacteria a with a 1 *µ*l inoculation loop from the frozen glycerol stock, and depositing in 50 *µ*L of fresh media. Next, we added 2 *µ*L of this bacteria suspension to 200 *µ*L of media in a 96-well plate. Cultures were then grown at 37°C with continuous double-orbital shaking in a Synergy H1 plate reader (BioTek) or an Epoch 2 plate reader (BioTek) for 24-32 hours, with absorbance at 600 nm (optical density or OD) measured every 5 mins. The growth rate *r* and the maximum optical density OD_*max*_ for each curve were determined using a MATLAB script, adapted from that used in [24], which performs a least-squares fit of *y* = ln[OD(*t*)*/*OD(0)] to the Gompertz equation [73]

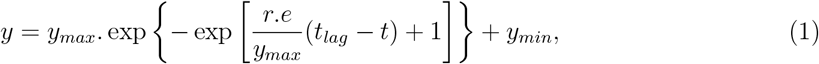

where *e* = exp(1) and OD_*max*_ = OD(0). exp(*y*_*min*_ + *y*_*max*_). A lag time *t*_*lag*_ was also determined from the data, although because the growth curves were started directly from frozen glycerol stocks, this should not be compared to more standard measures of lag time.

The reason for measuring bacterial growth directly from the glycerol stock (as opposed to first preparing an overnight culture and then diluting) was to minimize the risk of new mutations occurring in the populations, which could result in the genomic analyses being less relevant to observed phenotypes.

To minimize the concentration of phages in the co-evolved bacteria samples, we centrifuged the original co-cultures to pellet the bacteria and replaced the supernatant before storage. However, as this process does not eliminate phages within infected cells, some phages did remain in the coevolved bacteria samples. These phages could, in principle, affect the measured growth rate *r*_0_ of the co-evolved bacteria populations. Therefore, we measured the phage concentration at the start and end of the growth curves and confirmed that they were at least several orders of magnitude lower than the bacterial concentration in the culture (Fig. S8). Therefore, we expect that the residual phages present have negligible effect on the measured bacterial growth rate.

### Phage quantification

Phage titre was measured using plaque assays in which 10-fold serial dilutions of phage samples were prepared and 5 *µ*L was spotted onto plates containing 10 mL of 0.5% LB-agar and 200 *µ*L of overnight BW25113 culture (the ancestral *E. coli* strain was used, unless otherwise stated). These plates were incubated overnight and visible plaques were counted the following day to determine phage titre in terms of plaque forming units (PFU) per volume. Each plaque is assumed to have originated with a single phage, and represents one PFU. The uncertainty in PFU measurements was determined from the standard error of ≥ 3 replicates.

Using this procedure, it is generally not possible with our system to resolve more than ∼30 plaques in a 5 *µ*L spot, as beyond this number, plaques begin to overlap and merge. In the case where the approximate phage concentration was known beforehand, as was the case for the independently-evolved populations due to the limited variation, plates were prepared containing 10 mL of 0.5% LB-agar, 200 *µ*L of overnight BW25113 culture, and 20 *µ*L of the relevant phage dilution. Using this method, it is possible to resolve up to ∼400 plaques per plate, and the uncertainty was determined from the square root of the number of plaques counted [74].

### Phage endpoint density assay

Endpoint density assays were performed to measure the phage titres produced by a given phage population in a given media condition. Each assay was performed by first preparing an overnight culture of BW25113. The culture was then centrifuged at 8,000 g for 5 mins to pellet the bacteria and the supernatant was discarded. This pellet was then re-suspended in fresh media to achieve an optical density of OD=0.1. Following this, the phage population being tested was diluted in fresh media to achieve a concentration of 400 pfu/*µ*l. Then, 200 *µ*l of the bacteria culture was combined with 50 *µ*l of the diluted phage in a 96-well plate, then incubated overnight. Assuming that OD=0.1 equates to 10^8^ cells/mL, this means each assay should contain 2 ×10^7^ cells and 2 ×10^4^ phages, corresponding to a multiplicity of infection (MOI) of 0.001. The following day, final phage titre was measured as previously described. At least three replicate assays were performed in each case.

### DNA extraction and sequencing

Population-level bacterial DNA was extracted directly from frozen glycerol stocks (i.e., without a further growth period) to minimize any further changes that could occur during this period. Glycerol stocks were thawed, and 600 *µ*L were taken and centrifuged at 8,000 g for 2 mins. The supernatant was discarded and the bacteria pellet was re-suspended in 1 mL PBS (Fisher) before being centrifuged again at 8,000 g for 2 mins. The supernatant was again discarded and the bacteria pellet was re-suspended in 200 *µ*L PBS. DNA was then extracted using the DNeasy Blood and Tissue Kit (Qiagen) according to the manufacturer’s instructions.

To extract DNA from bacterial isolates, the glycerol stocks were streaked onto 1.5% LB-agar plates and incubated at 37°C overnight. Single colonies were picked, inoculated into 1 mL M9 media with osmolality matched to the population’s evolutionary history, and grown overnight. The culture was then centrifuged at 8,000 g for 2 mins, the supernatant was discarded, and the bacteria pellet was re-suspended in 200 *µ*L PBS. DNA was then extracted as before using the DNeasy Blood and Tissue Kit (Qiagen) according to the manufacturer’s instructions.

Where possible, phage DNA was directly extracted from refrigerated samples without additional growth to minimize the risk of further changes. For phage samples with concentrations exceeding 10_8_ pfu/mL, a portion was diluted in PBS to obtain 1 mL of a diluted sample at approximately 10_8_ pfu/mL, from which DNA was then extracted. To avoid contamination with bacterial DNA, 20 units of RNase-Free DNase I (Norgen Biotek) was added to each sample and incubated for 15 mins at room temperature, followed by DNase I inactivation at 75°C for 5 mins. Phage DNA was then extracted using the Phage DNA Isolation Kit (Norgen Biotek) according to the manufacturers instructions, including the addition of 4 *µ*L of 20 mg/mL Proteinase K (Qiagen) to increase DNA yields.

For samples with concentrations below 10_8_ pfu/mL, 1 mL of the original sample was used for DNA extraction. Additionally, 20 *µ*L of the original sample was utilized to produce a higher density lysate on the ancestral strain under the original media conditions, and DNA was extracted from these samples as well. Among the nine populations in this category, five yielded sufficient genetic material from the original samples. A comparison of these five populations indicates that minimal changes occurred during the additional growth step (Fig. S9).

Library preparation and sequencing for the bacteria and phage populations (excluding the three re-runs of the phages independently evolved in high osmolality) were performed by the Sequencing & Bioinformatics Consortium at The University of British Columbia. Extracted DNA was quantified using Qubit fluorometry. Genomic libraries were prepared using the Illumina DNA Prep kit (Illumina). Samples were sequenced with the Illumina NextSeq 2000 sequencing platform generating paired end 150 bp reads. Average coverage achieved was ∼450× for bacterial samples and ∼300× for phage samples. For the bacteria isolates and the three phage re-runs, library preparation and sequencing was performed by Plasmidsaurus. Genomic libraries were prepared using the Illumina DNA Prep kit and sequenced with the Illumina NovaSeq platform generating paired end 150 bp reads. Average coverage achieved was ∼1, 500× for the bacterial isolates and ∼35, 000× for the phage samples.

### Analysis of sequencing data

Raw reads were first processed using fastp [75] (v0.23.4) to filter low-quality reads and trim adapter sequences using default options. Mutations in the ancestor populations compared to the reference genomes were then identified using breseq [76] (v0.38.3) and Bowtie2 [77] (v2.5.3) with default options. The reference genomes were GenBank CP009273.1 [78] for *E. coli* BW25113 and NCBI NC_000866.4 [37] for T4 phage. The mutations identified in the ancestral populations were then used to create new reference sequences that were used as the point of comparison for all evolved populations. The mutations and allele frequencies in the evolved populations, relative to the ancestral populations, were then determined by running breseq in polymorphism mode with default filters.

The per gene probability of having at least one mutation *P*_*gene*_ was estimated from the observed mutation frequencies as

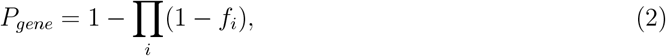

where *f*_*i*_ represents the frequency of mutation *i* affecting the gene. This calculation assumes independence of mutations across sites, and that mutation frequencies correspond to non-overlapping events within the genome. In practice, neither assumption is true. Mutations may be correlated, as suggested by the strong correlation between the R195P mutation in *ompC* and the noncoding (205/1542 nt) mutation in *rrsG* (Fig. S6). Additionally, multiple alleles can affect the same site, and single mutation events affecting neighboring sites may be incorrectly called as two distinct events, or vice-versa. These factors can bias *P*_*gene*_ away from the true fraction of the population carrying at least one mutation. However, given that we use this metric simply to broadly identify genes that are frequently mutated, the precise values are not important.

### Efficiency of plating (EOP)

To quantify the ability of a given phage population to infect and replicate in a given bacteria population, efficiency of plating (EOP) assays were performed. To do this, a 10-fold serial dilution of the phage population was prepared and spotted (as described in the ‘Phage quantification’ section) on a plate containing the reference bacterial population (ancestral strain) and a plate containing the test bacterial population. The overnight cultures of the bacterial populations were grown directly from glycerol stocks. The EOP was then determined as the ratio of the apparent titre on the test strain *ρ*_*test*_ to the apparent titre on the reference strain *ρ*_*anc*_

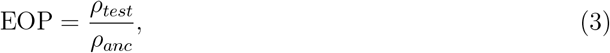

with an EOP<1 indicating that the phage less efficient at infecting the test bacteria compared to the reference. At least three biological replicate assays were performed in each case. For measurements involving the co-evolved bacteria populations where some phage were still present in the glycerol stocks (Fig. S8) overnight cultures were centrifuged and washed in PBS prior to plating to minimize the number of phages present in the lawn, and allow visualization of the plaques formed by phage sample dilutions (Fig. 3a).

### Mucoid quantification

To quantify the level of mucoidy in evolved bacterial populations, the populations were streaked from glycerol stocks onto 1.5% LB-agar plates and incubated at 37 °C overnight. The following day, images of all plates were captured for analysis. We employed a pairwise comparison approach, where the user identified which of two bacterial plates (if either) appeared more mucoid. This comparative method was used to avoid absolute, single-instance evaluations (e.g., assigning a single numerical score to an image), which are prone to bias and higher variability [79–81].

To generate a quantitative mucoid score from these comparisons, we implemented the Elo rating system, commonly used in competitive settings (such as chess) to rank players. Here, images of the bacterial plates are treated as ‘competitors,’ and user judgments about the relative mucoidy between two plates inform rating adjustments.

The Elo ratings for each image were initialized at 1500 and with each comparison, the ratings were updated according to the equation:

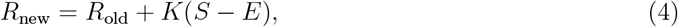

where *R*_new_ and *R*_old_ are the new and old ratings, *K* is a constant that controls the magnitude of changes in rating (set to *K* = 32 in our case), *S* is the score (1 for a win, 0 for a loss, 0.5 for a draw), and *E* is the expected score based on the current ratings. For two images, A and B, the expected scores *E*_*A*_ and *E*_*B*_ are calculated as:

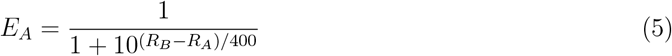

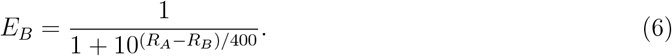

As more comparisons are made, the Elo scores of the images stabilize, reflecting the relative degree of mucoid appearance perceived by the user. The comparisons were continued until each plate had been evaluated at least 105 times, with the average Elo rating from the comparisons 95 to 105 taken as the final mucoid score for that ‘competition.’ The entire procedure (including streaking, imaging, and pairwise comparisons) was repeated three times, producing three independent mucoid score measurements for each population.

## Acknowledgments

The authors acknowledge support from the Michael Smith Health Research BC Trainee Award (RT-2023-3174, to MH), the Natural Sciences and Engineering Research Council of Canada (NSERC) Discovery Grants Program (RGPIN-2019-04591, to CT), the Canadian Institute for Advanced Research / Humans and the Microbiome (FL-001253 Appt 3362, to CT), the Michael Smith Foundation for Health Research Scholar Award (18239, to CT), the Canada Tier 2 Research Chair, Quantitative Microbiota Biology for Health Applications (CRC-2022-00036, to C.T.), and the Canada Foundation for Innovation / Infrastructure Operating Fund (38277, to CT). This research was supported in part through computational resources and services provided by Advanced Research Computing (ARC) at the University of British Columbia. Library preparation and sequencing were performed by the Sequencing & Bioinformatics Consortium at The University of British Columbia. The authors thank A. Srinivas, H. Ghezzi, and J. He for providing feedback on earlier drafts of the manuscript. The authors acknowledge that this work was completed on the traditional, ancestral, and unceded territory of the x^w^mǝθk^w^ǝỷǝm (Musqueam) people. The authors encourage the reader to learn about the history of the land they work on at Native Land Digital (www.native-land.ca).

