## Supplementary Information for "Osmotic conditions shape fitness gains and resistance mechanisms during *E. coli* and T4 phage co-evolution"

**S3 Correlation between *rrsG* and *ompC* mutations** **8**

**S4 Original results from independent phage evolution in high osmolality** **9**

**S5 Phage concentrations at start and end of co-evolved bacteria growth curves** **11**

**S6 The impact of an additional growth cycle on phage population sequencing** **12**

### S1 Bacterial growth curves

In the main text, we focused on exponential growth rate $r$ as our measure of bacterial fitness (Fig. 1). Here we present the growth curves underlying this analysis – Fig. S1.

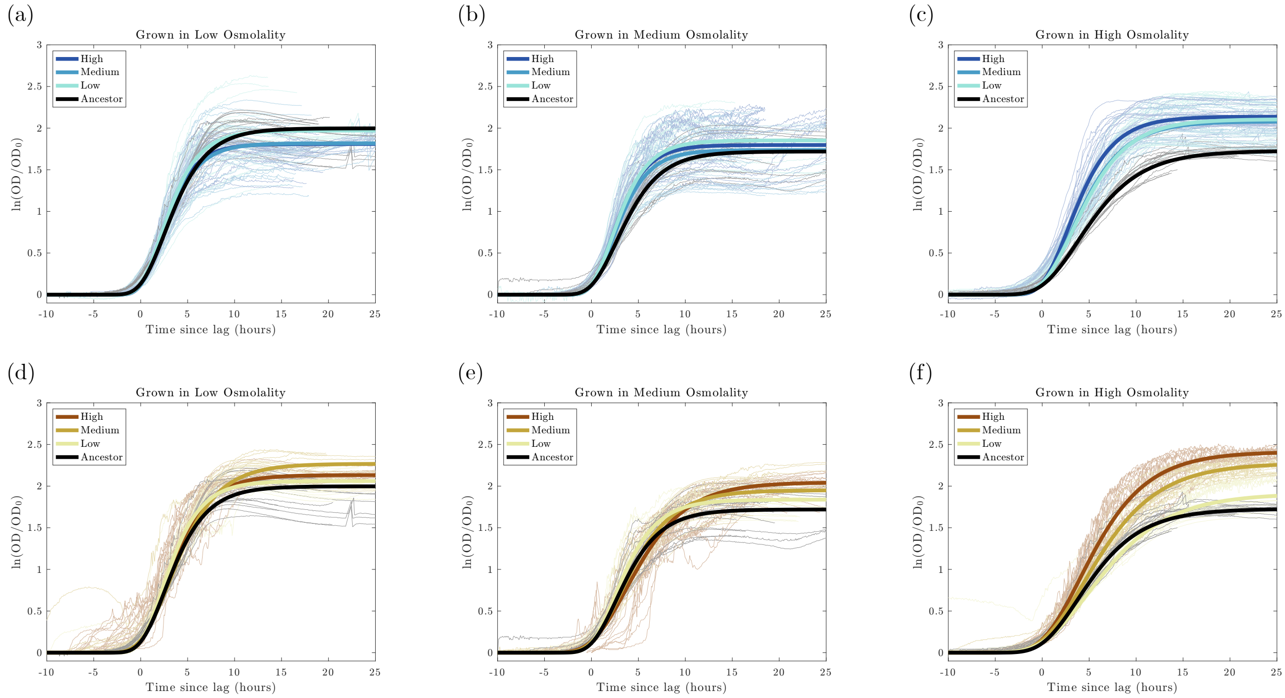

Figure S1: Growth curves of the ancestral strain and all independently (a-c) and co-evolved (d-f) bacteria populations after 36 passages in low, medium, and high osmolality (∼240, 450, and 920 mOsm/kg $H_2O$, respectively). Individual replicate curves are shown as faint lines colored according to the evolutionary history of the population. As described in the main text, the exponential growth rate $r$, maximum optical density, and lag time for each curve was determined by fitting the data to the Gompertz equation [1]. In these plots, each replicate is shown in relation to its extracted lag time. Thick lines show the Gompertz curve defined by the average fit parameters from all of the replicates with that evolutionary history.

From these growth curves, we also extract the average doubling times $t_d = \ln(2)/r$ of the ancestral strain in each condition, which we use in our mathematical model in the following section (Fig. S2).

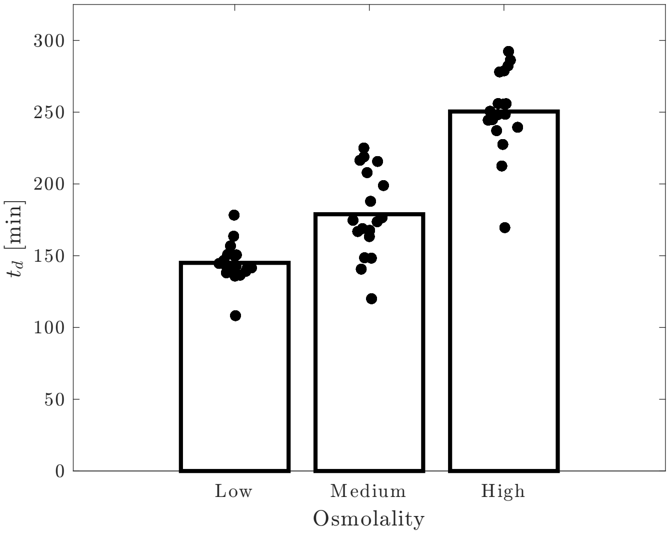

Figure S2: Doubling time $t_d$ for the ancestral strain in each osmotic condition.

### S2 The mechanistic origin of increased phage endpoint densities

In the main text, we showed that all independently evolved phages exhibited much larger endpoint densities than the ancestral phage in all conditions (Fig. 1). In particular, phages evolved and assayed under high osmotic conditions had a relative fitness of $\log_{10}(w) = 1.7$, corresponding to a 50-fold higher endpoint density than the ancestral phage. Here, we investigate potential mechanistic origins for this increase.

#### S2.1 Mathematical model of infection dynamics

To explore the mechanistic origin of the large increases to endpoint density $\rho$ that we observe, we used a simple mathematical model to explore whether changes to the phage life-history parameters could offer a plausible explanation. Briefly, we model the interactions between 3 populations: viruses (phages) $V$, uninfected host bacteria $B$, and infected host bacteria $I$. The infection process can be summarized as

$$V + B \xrightarrow[\alpha]{\text{rate}} I \xrightarrow[\tau]{\text{delay}} \beta V, \tag{1}$$

where $\alpha$ is the phage adsorption rate, $\beta$ is the phage burst size (the number of new phages released upon host lysis), and $\tau$ is the time between phage adsorption and host lysis (aka lysis time).

Neglecting any stochasticity, the dynamics of the populations can be described using a set of delayed differential equations (DDEs):

$$\frac{dV}{dt} = -\alpha V(B + I) + \beta\alpha V_{t-\tau}B_{t-\tau}, \tag{2a}$$

$$\frac{dB}{dt} = r_0 B\left(1 - \frac{B + I}{B_{max}}\right) - \alpha VB, \tag{2b}$$

$$\frac{dI}{dt} = \alpha VB - \alpha V_{t-\tau}B_{t-\tau}, \tag{2c}$$

where $V$, $B$, and $I$ indicate the size of the populations as a function of time. The subscript $t - \tau$ is used to indicate the time at which those terms are evaluated. We account for the logistic growth of bacteria with growth rate $r_0$ and carrying capacity $B_{max}$. We assume that infected cells are unable to replicate but still use resources to produce new phages and therefore contribute to the carrying capacity. The growth rate $r_0$ can also be expressed in terms of the bacterial doubling time $t_d = \ln(2)/r_0$.

It should be noted that we allow for adsorption of T4 to previously infected cells, though this has no impact on the state of the cell. Phage T4 has a well characterized superinfection-exclusion mechanism that causes rapid inhibition of DNA injected into previously infected cells [2]. It should also be noted that all key parameters ($\alpha$, $\beta$, $\tau$, and $r_0$) vary depending on the osmotic conditions (Sec. S2.4.1 and Table S1).

We then solved this model numerically to explore how the three phage life-history parameters ($\alpha$, $\beta$, and $\tau$) impact phage endpoint density $\rho$ – see Sec. S2.4.1. We find that increases in endpoint density similar in magnitude to those observed in our experiments – $\log_{10}(w = \rho/\rho_{anc}) \sim 1$ under high osmolality conditions – could be achieved by decreasing the adsorption rate $\alpha$ (Fig. S3a) or increasing the burst size $\beta$ (Fig. S3b). The modeling also showed that endpoint density could be increased by increasing the lysis time $\tau$ (Fig. S3c), although the predicted magnitude of the increase in endpoint density was substantially lower than we observed experimentally.

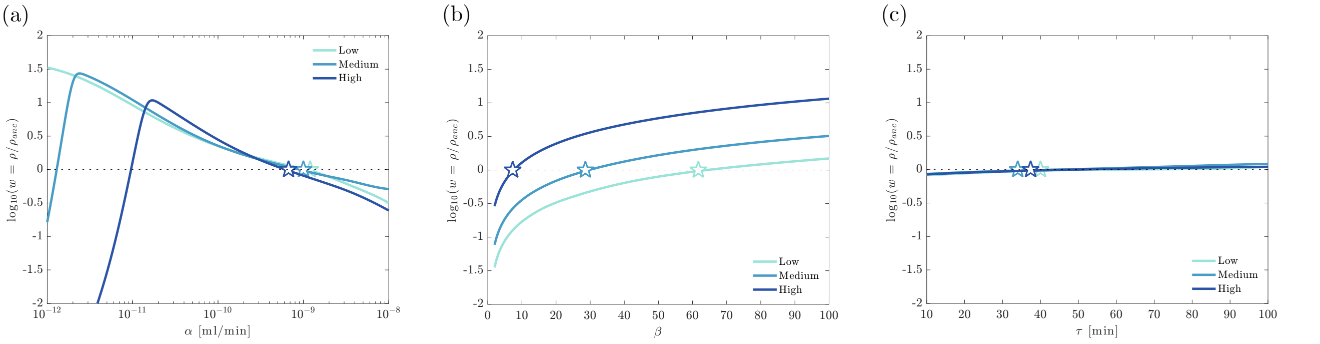

Figure S3: Relative fitness $w = \rho/\rho_{anc}$ as a function of adsorption rate $\alpha$ (e), burst size $\beta$ (f), and lysis time $\tau$ (g) predicted from the mathematical model of phage infection. Stars indicate the parameters for the ancestral phage in each condition.

#### S2.2 Experimental measurements of life-history parameters

Given that our modeling indicates that changes to life-history parameters could offer a plausible explanation for the large increases in endpoint phage density, we performed preliminary adsorption rate assays and one-step growth curves with the ancestral phage and each of the independently evolved phage populations (Sec. S2.4.2 and Sec. S2.4.3).

However, the experimental results do not show changes that are quantitatively consistent with our model predictions. While the average adsorption rate of the evolved populations decreased under all osmotic conditions (Fig. S4a), the magnitude of the reduction differed across conditions, with larger declines under low (−33%) and medium (−39%) osmolality, than under high osmolality (−14%). This despite the fact that high osmolality was the condition where we observed the largest increase in endpoint density experimentally. In contrast, mean burst size increased substantially under high osmotic conditions (55%), showed a modest increase under medium osmotic conditions (13%), and decreased under low osmotic conditions (−32%) – Fig. S4b.

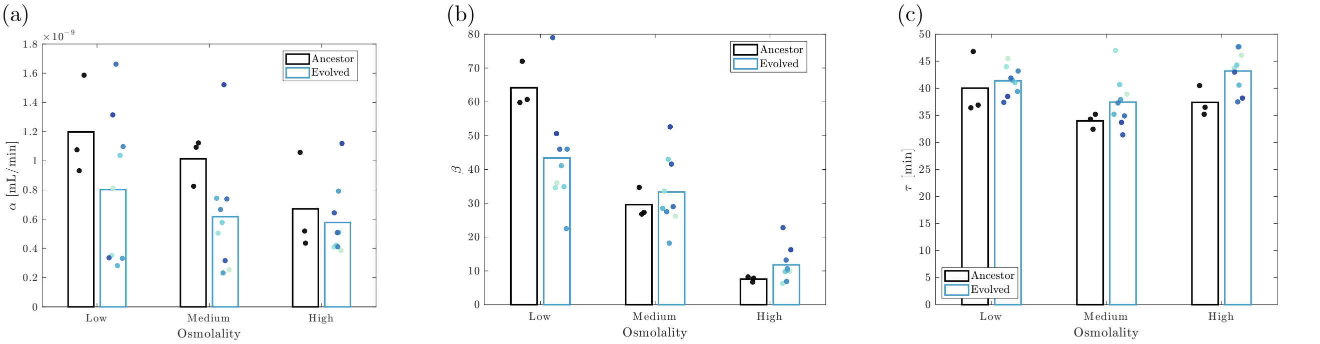

Figure S4: Experimental measurements of adsorption rate $\alpha$ (a), burst size $\beta$ (b), and lysis time $\tau$ (c) of the ancestral and independently evolved phages in each osmotic condition.

Using the mean adsorption rate, burst size, and lysis time of all experimentally evolved populations, our model predicts a relative fitness in each condition of $\log_{10}(w) \sim -0.1$ in low osmolality, $\log_{10}(w) \sim 0.1$ in medium osmolality, and $\log_{10}(w) \sim 0.2$ in high osmolality. These predictions are not consistent with our experimental data, which show $\log_{10}(w)$ ranges of 0.25–0.39 in low osmolality, 0.46–0.48 in medium osmolality, and 0.79–1.7 in high osmolality. This indicates that the observed increases in endpoint density cannot easily be attributed to the traditional life-history parameters.

We note that no replicate measurements were taken for the evolved populations; however, based on our model, if decreased adsorption rate or increased burst size were the cause of the productivity increases, we would expect to see changes substantially larger than those we observe.

#### S2.3 Phage stability measurements

To verify that phages were stable in our media conditions, we inoculated fresh media with a small volume of the ancestral phage and measured the population size over a period of two weeks – Fig. S5.

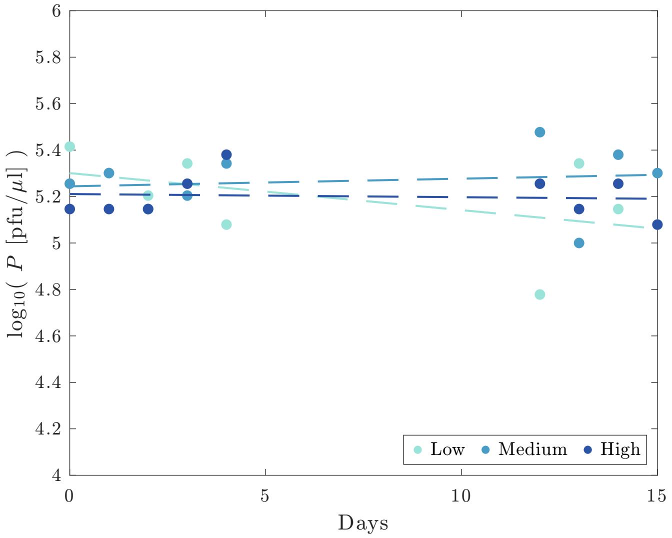

Figure S5: Ancestral phage titres during storage in the three media conditions for 2 weeks. No reduction in the viable population was observed.

We found no statistically significant reduction in the viable population was observed under any of the conditions ($p_{low} = 0.16$, $p_{medium} = 0.70$, $p_{high} = 0.81$). This is also consistent with the stability in viable titres in all of our stocks over the span of the study.

#### S2.4 Supplementary methods

##### S2.4.1 Numerical solutions to the model

The default parameters used in our model are shown in Table S1. At time $t = 0$ we set $B_0 = 2 \times 10^7$ and $V_0 = 2 \times 10^4$ to match the initial conditions of our phage endpoint density assays. We set as initial conditions no infected bacteria ($I_0 = 0$). Solutions were then determined using the Euler method with a step size of $\Delta t = 0.01$, as we did not observed any numerical instabilities at this level of precision. The parameter values were chosen to match values measured in experiments, with the same equivalent units (e.g., an experimental lysis time of 20 mins would be represented with a value of 20 in the model).

We used our mathematical model to characterize how the fitness or selective advantage of the phage population depended on the parameters $\alpha$, $\beta$, and $\tau$. The endpoint density was defined in the same manner as it was in our experimental assay, as the final titre ($\rho$) of the phage population reached after 24 hours of growth. As our model did not include any phage-resistance, the bacteria

Table S1: Default model parameters.

| Parameter | Osmolality [mOsm/kg $H_2O$] | | | Source |
|---|---|---|---|---|
| | 240 | 450 | 920 | |
| $B_0$ [ml$^{-1}$] | $2{\times}10^7$ | $2{\times}10^7$ | $2{\times}10^7$ | * |
| $B_{max}$ [ml$^{-1}$] | $10^9$ | $10^9$ | $10^9$ | [3] |
| $t_d$ [min] | 145 | 179 | 250 | Fig. S2 |
| $V_0$ [ml$^{-1}$] | $2{\times}10^4$ | $2{\times}10^4$ | $2{\times}10^4$ | * |
| $\alpha$ [ml/min] | $1.2{\times}10^{-9}$ | $1.0{\times}10^{-9}$ | $6.7{\times}10^{-10}$ | Fig. S4a |
| $\beta$ | 61.7 | 28.6 | 7.3 | Fig. S4b |
| $\tau$ [min] | 40.0 | 34.0 | 37.4 | Fig. S4c |
| $\Delta t$ [min] | 0.01 | 0.01 | 0.01 | - |

Where parameters different to the defaults are used, this is indicated in the text. Notes: ($*$) chosen to match the initial conditions of the endpoint density assays.

population usually went extinct ($B + I < 0.1$) before the 24 hour mark was reached, so for computational reasons this was the point at which phage titre was measured in these instances. The relative fitness $w$ of a given 'mutant' phage ($mut$) relative to the ancestral phage ($anc$) is therefore given by

$$w = \frac{\rho_{mut}}{\rho_{anc}}, \tag{3}$$

with 'ancestor' in this case referring to the default parameters given in Table S1.

##### S2.4.2 Adsorption rate measurements

Initially, 1 mL ancestor overnight culture was centrifuged at 8,000 g for 1 min, then the supernatant was removed, and the pellet re-suspended in 1 mL of fresh media. To determine bacterial concentration, 10-fold serial dilutions were prepared and three technical replicates of 5 $\mu$L were spotted onto 1.5% LB-agar. These plates were incubated at 37°C overnight and the average bacterial concentration was determined by counting colonies in the three replicates.

To perform the adsorption rate assay, 10 $\mu$L of phage ($\sim 10^7$ pfu/$\mu$L) were added to the bacteria culture, vortexed, and kept at 37°C with constant shaking. Every 45 seconds for 4.5 mins, a 100 $\mu$L sample of the culture was taken and added to a tube containing 75 $\mu$L PBS and 25 $\mu$L chloroform and vortexed. The chloroform kills both infected and uninfected bacteria but has no effect on the free phage population. These samples were then serially diluted and spot plated as previously described to determine the concentration of free phage.

Assuming that the concentration of available hosts is approximately constant over this short time period, the slope of log-transformed phage concentration $\ln(V)$ against time will be equal to the adsorption rate $\alpha$ multiplied by the bacterial density $B_0$ i.e.,

$$\ln(V) = -\alpha B_0 t + \ln(V_0), \tag{4}$$

where $V_0$ represents the initial concentration of free phage. Adsorption rate was then determined from the slope divided by the bacterial concentration.

##### S2.4.3 One-step growth curves

This procedure was adapted from Ellis and Delbrück [4]. Initially, 100 $\mu$L of ancestor overnight culture was diluted in 9.9 mL of fresh media and incubated at 37°C with constant shaking to generate an exponentially growing culture. This culture was then centrifuged at 8,000 g for 1 min, the supernatant was removed, and the bacteria were re-suspended in 100 $\mu$L of fresh media. This concentration step was used to ensure that initial MOI was low (MOI<0.1) and that phage adsorption occurred rapidly, thereby minimizing co-infections and synchronizing infections.

At the start of the experiment ($t = 0$), 1.5 $\mu$L of phage lysate was added to the concentrated exponential culture and incubated at 37°C with constant shaking for 1 min to allow infection. The culture was then serially diluted by factors of 10 in fresh media to obtain two 'test' cultures $D_n$ and $D_{n+1}$, corresponding to dilution factors of $10^n$ and $10^{n+1}$ respectively, which were kept incubated at 37°C with constant shaking. Every 3-5 mins for approximately 1 hour, 20 $\mu$L of each 'test' culture was plated with 200 $\mu$L of overnight culture (grown in LB) and 10 mL of molten 0.5% LB-agar. The dilution factors $10^n$ and $10^{n+1}$ depended on the initial concentration of phage used, and were chosen to produce a countable number of plaques when plating. As before, these plates were incubated overnight at 37°C and visible plaques were counted the following day.

To determine the mean lysis time $\tau$ and burst size $\beta$ from the data, we assumed that lysis events are normally distributed and fit the following function [5]

$$n_p = \frac{\beta}{2}\left[1 + \text{erf}\left(\frac{t-\tau}{\sigma_\tau\sqrt{2}}\right)\right] + 1, \tag{5}$$

where $n_p$ is the number of PFUs relative to the initial number of PFUs as a function of time, and $\sigma_\tau$ is the standard deviation of lysis times. The number of initial PFUs was estimated as the average of at least the first 3 measurements, although this number could be increased provided no lysis had yet occurred. The erf($x$) function is the error function given by:

$$\text{erf}(x) = \frac{2}{\sqrt{\pi}}\int_0^x e^{-y^2}\,dy. \tag{6}$$

### S3 Correlation between *rrsG* and *ompC* mutations

In the main text, we showed that all bacteria populations co-evolved under low and medium osmotic conditions, and one co-evolved under high osmotic conditions, had mutations in the *ompC* gene (Fig. 4). We also noted that among these, a single mutation (R195P) dominated, and co-occurred with a noncoding (205/154 nt) mutation in *rrsG* in 6 of the co-evolved populations. The frequency of these mutations were highly correlated, with a Pearson correlation coefficient of 0.952 (Fig. S6). Neither mutation appeared in any other co-evolved population.

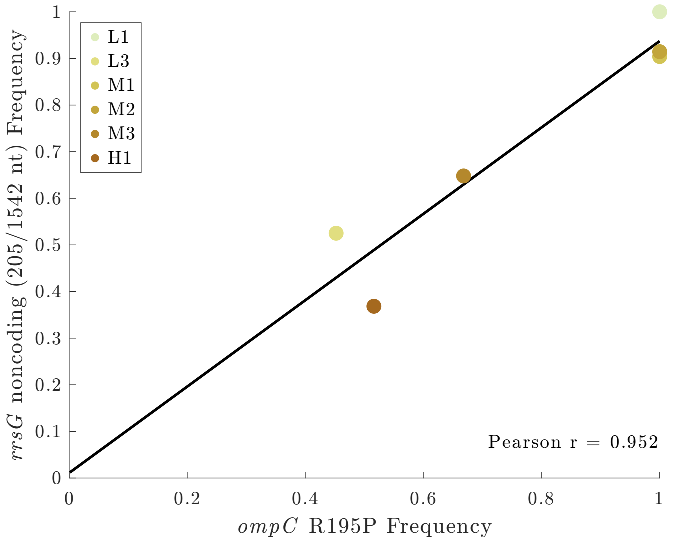

Figure S6: Correlation in frequency of two specific mutations affecting *ompC* and *rrsG*. The black line shows a linear fit to the data.

### S4 Original results from independent phage evolution in high osmolality

As noted in the main text, the first time we performed the independent phage evolution experiment, a sudden drop in the three high osmolality populations occurred between passages 11 and 15 (Fig. S7a). We suspect this was due either to the selection of a bacterial colony that happened to be resistant to phage infection, or to a procedural error during passaging (e.g., failure to inoculate bacterial sub-cultures on a given day). Either of these scenarios would have effectively diluted the phage by a factor of 100 on that day, which is consistent with the observed data.

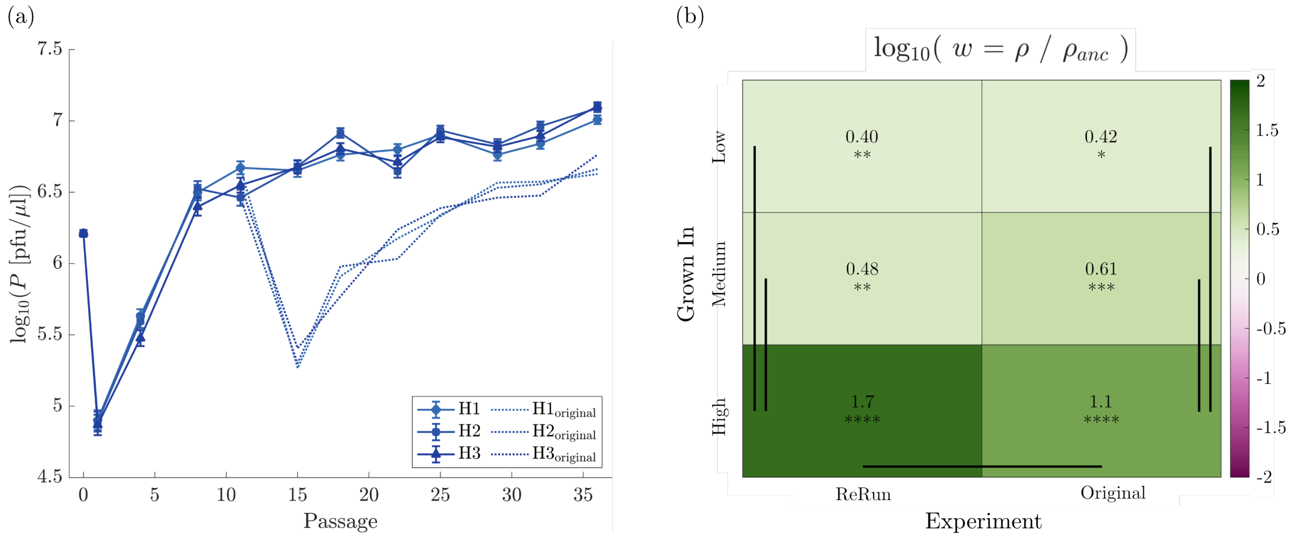

Figure S7: (a) Phage population size $P$ throughout the experiment measured by plaque assay on the ancestral bacteria strain. The solid lines show the rerun populations discussed in the main text, while the dotted lines show the original lines where a population crash occurs between passages 11 and 15. (b) The relative fitness $w$ of the final phage populations independently evolved in high-osmolality, from the rerun and original lines. Relative fitness is defined as the ratio of the population's endpoint density $\rho$ relative to the endpoint density of the ancestral phage $\rho_{anc}$ grown in the same conditions. This was determined by measuring the final phage concentration reached given a defined initial culture with the ancestral strain - see Methods in main text. Statistical comparisons between evolved and ancestor populations determined using two-sample t-tests followed by the Bonferroni correction applied as $p$-value adjustment. * $p < 0.05$, ** $p < 0.01$, *** $p < 0.001$, and **** $p < 0.0001$. Statistical comparisons between evolved groupings made using two-way ANOVA followed by Tukey's honestly significant difference test. Any comparison with $p < 0.05$ is shown using a solid bar. Note that only statistically significant comparisons *within* each "Grown In" or "Experiment" condition are shown.

To confirm this, we restarted these three lines from the passage 11 samples that were taken before the population drops, and no drops were observed the second time around, suggesting our hypothesis was correct. The results from these re-run lines are discussed in the main text; however, we include here results relating to the original lines for comparison. The relative fitness $w$ of the final populations, calculated as the ratio of phage endpoint densities ($w = \rho/\rho_{anc}$), are qualitatively consistent (Fig. S7b). All populations in both the original and rerun lines exhibited higher endpoint densities than the ancestor ($w > 1$) under all osmotic conditions, and the increase was greatest when the phages were assayed under the high-osmolality conditions. This effect was even larger in the rerun lines, which showed significantly higher relative fitness compared to the original lines under high-osmolality conditions.

### S5 Phage concentrations at start and end of co-evolved bacteria growth curves

In the main text, we described how, to minimize the number of phages present in the co-evolved bacteria samples, we centrifuged the co-cultures (8,000 g for 5 mins) to pellet the bacteria, then discarded the supernatant and resuspended the bacteria in fresh media prior to storage. While this eliminated most phages, some still remained inside infected cells. The presence of these phages could, in principle, reduce the apparent growth rate of the bacteria (Fig. 1).

Therefore, we measured the concentration of phages present in the populations both before and after 24 hours of growth (Fig. S8).

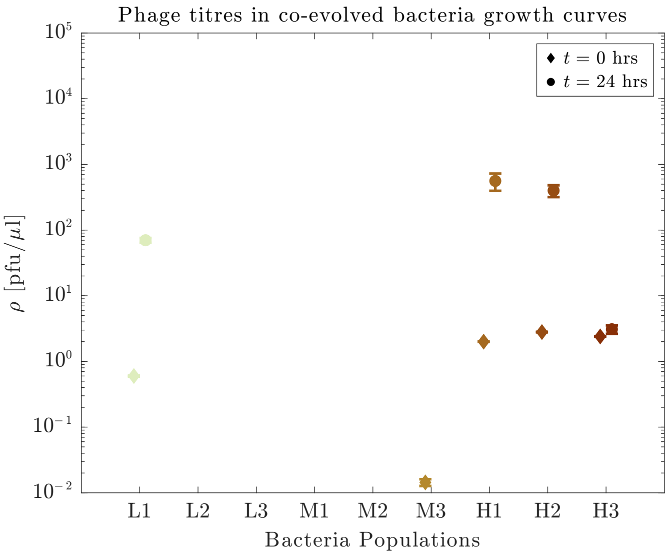

Figure S8: Phage concentrations presented at the start and end of co-evolved bacteria growth curves.

The phage concentrations at the start of the growth curves are all $< 10^1$ pfu/$\mu$l, and $< 10^3$ pfu/$\mu$l after 24 hours of growth. This is approximately three orders of magnitude lower than the concentration of bacteria in the cultures at those times; therefore, we expect the impact on apparent growth rate to be negligible.

#### S6 The impact of an additional growth cycle on phage population sequencing

As discussed in the main text, several of the phage samples contained phage concentrations below the level recommended by the extraction kit manufacturer. In these cases, we used 20 $\mu$L of the original sample to produce a higher density lysate on the ancestor bacteria under the original media conditions, and extracted DNA from those samples as well. Among the nine populations in this category (all of the co-evolved phage populations), five yielded sufficient genetic material from the original samples. A comparison of these five populations indicates that minimal changes occurred during the additional growth step (Fig. S9), and to ensure consistency, all of the sequencing results for the co-evolved phage populations refer to the results from the higher density samples.

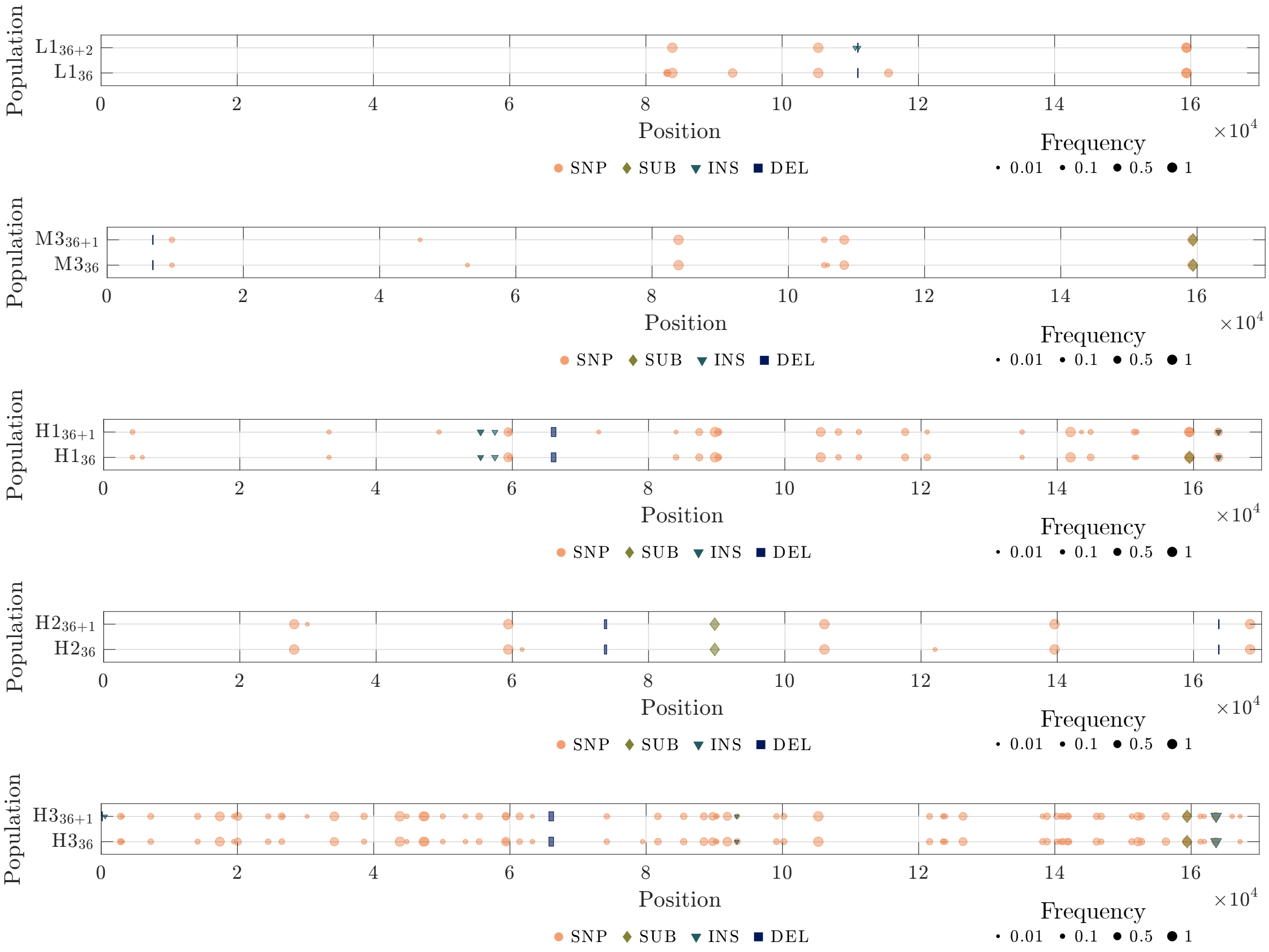

Figure S9: Mutations in the co-evolved phage populations before (bottom) and after (top) another cycle of growth, which was used to increase phage density for DNA extraction. The other four populations before the additional growth cycle did not yield enough DNA for sequencing. Length of the deletion bars represent the length of the deletion, unless the deletions are less than 100 bp, in which case they are shown as 100 bp for visibility. Labels show annotations for key genes discussed in the text. Population subscripts indicate the passage of the sample.
