## Supplementary material for "Osmotic conditions shape fitness gains and resistance mechanisms during *E. coli* and T4 phage co-evolution": Table S1: Table_S1_IndBac.html

Mutation Comparison


| Predicted mutations | | | | | | | | | | | | | |
| --- | --- | --- | --- | --- | --- | --- | --- | --- | --- | --- | --- | --- | --- |
| position | mutation | L1_36_ | L2_36_ | L3_36_ | M1_36_ | M2_36_ | M3_36_ | H1_36_ | H2_36_ | H3_36_ | annotation | gene | description |
| 104,042 | A→T |  |  |  |  |  | 13.0% |  |  |  | intergenic (+81/‑150) | *lpxC* → / → *secM* | UDP‑3‑O‑acyl N‑acetylglucosamine deacetylase/regulator of secA translation |
| 161,051 | A→T |  | 10.3% |  |  |  |  |  |  |  | intergenic (+30/‑166) | *hrpB* → / → *mrcB* | putative ATP‑dependent helicase/fused glycosyl transferase and transpeptidase |
| 284,974 | A→T |  | 7.3% |  | 7.5% |  |  |  |  | 8.3% | intergenic (‑101/+38) | *yagI* ← / ← *argF* | CP4‑6 prophage; putative DNA‑binding transcripti onal regulator/ornithine carbamoyltransferase 2, chain F; CP4‑6 prophage |
| 284,986 | A→T |  |  | 5.7% |  |  |  |  |  |  | intergenic (‑113/+26) | *yagI* ← / ← *argF* | CP4‑6 prophage; putative DNA‑binding transcripti onal regulator/ornithine carbamoyltransferase 2, chain F; CP4‑6 prophage |
| 288,008 | T→A |  | 6.0% |  |  |  |  | 10.9% |  |  | intergenic (+66/+25) | *yagJ* → / ← *yagK* | CP4‑6 prophage; uncharacterized protein;Phage or Prophage Related/CP4‑6 prophage; conserved protein |
| 295,710 | G→C |  |  |  |  |  | 7.9% |  |  |  | P312P (CCC→CCG) | *paoC* ← | PaoABC aldehyde oxidoreductase, Moco‑containing subunit |
| 375,461 | A→C |  |  |  | 7.2% | 6.1% |  | 8.2% |  |  | intergenic (‑124/+64) | *frmR* ← / ← *yaiO* | regulator protein that represses frmRAB operon/outer membrane protein |
| 662,155 | T→A |  |  |  |  | 6.1% |  |  |  |  | I507F (ATT→TTT) | *mrdA* ← | transpeptidase involved in peptidoglycan synthes is (penicillin‑binding protein 2) |
| 695,147 | (ACCTGCC)_1→2_ |  |  |  |  |  |  | 6.0% |  |  | coding (636/753 nt) | *umpH* ← | UMP phosphatase |
| 703,476 | C→A |  |  |  |  | 8.6% |  |  |  |  | intergenic (+263/‑314) | *glnS* → / → *chiP* | glutamyl‑tRNA synthetase/chitoporin, uptake of chitosugars |
| 780,449 | T→G |  |  |  | 6.1% |  |  |  |  |  | T109P (ACC→CCC) | *ybgS* ← | putative periplasmic protein |
| 780,465 | G→T |  |  |  |  |  |  | 6.4% |  |  | I103I (ATC→ATA) | *ybgS* ← | putative periplasmic protein |
| 843,405 | G→C |  |  |  |  | 19.0% |  |  |  |  | S19C (TCT→TGT) | *glnH* ← | glutamine transporter subunit |
| 852,559 | A→T |  |  |  | 6.7% |  |  |  |  |  | N381Y (AAC→TAC) | *ybiT* → | putative transporter subunit of ABC superfamily: ATP‑binding component |
| 910,654 | C→G |  |  |  |  | 6.7% | 6.9% | 7.9% |  | 5.1% | intergenic (‑341/+154) | *ybjE* ← / ← *aqpZ* | putative transporter/aquaporin Z |
| 910,667 | C→G | 6.9% | 5.1% |  |  | 5.6% | 7.6% | 6.9% |  |  | intergenic (‑354/+141) | *ybjE* ← / ← *aqpZ* | putative transporter/aquaporin Z |
| 910,669 | A→T | 11.3% | 7.7% |  |  | 8.7% | 9.1% |  |  | 10.9% | intergenic (‑356/+139) | *ybjE* ← / ← *aqpZ* | putative transporter/aquaporin Z |
| 910,683 | G→C |  |  |  |  |  | 6.2% |  |  |  | intergenic (‑370/+125) | *ybjE* ← / ← *aqpZ* | putative transporter/aquaporin Z |
| 1,139,001 | T→A |  |  | 7.9% |  |  |  |  |  |  | I275F (ATC→TTC) | *rne* ← | fused ribonucleaseE: endoribonuclease/RNA‑bindin g protein/RNA degradosome binding protein |
| 1,191,676 | C→T |  | 7.1% |  |  |  |  |  |  |  | H366H (CAC→CAT) | *icd* → | e14 prophage; isocitrate dehydrogenase, specific for NADP+ |
| 1,191,688 | C→T |  | 6.5% |  |  |  |  |  |  |  | T370T (ACC→ACT) | *icd* → | e14 prophage; isocitrate dehydrogenase, specific for NADP+ |
| 1,191,701 | T→C |  | 6.1% |  |  |  |  |  |  |  | L375L (TTA→CTA) ‡ | *icd* → | e14 prophage; isocitrate dehydrogenase, specific for NADP+ |
| 1,191,703 | A→G |  | 6.1% |  |  |  |  |  |  |  | L375L (TTA→TTG) ‡ | *icd* → | e14 prophage; isocitrate dehydrogenase, specific for NADP+ |
| 1,196,769 | T→A |  |  |  |  | 21.3% |  |  |  |  | intergenic (‑281/‑184) | *xisE* ← / → *ymfI* | e14 prophage; putative excisionase/e14 prophage; uncharacterized protein |
| 1,342,337 | A→T |  |  |  |  | 7.6% |  |  |  |  | V278D (GTC→GAC) | *rnb* ← | ribonuclease II |
| 1,364,337 | G→T |  |  |  |  |  |  |  | 10.7% |  | intergenic (+77/‑136) | *pspE* → / → *ycjM* | thiosulfate:cyanide sulfurtransferase (rhodanese )/alpha amylase catalytic domain family protein |
| 1,471,657 | A→T |  |  |  |  | 5.9% |  |  |  |  | T753S (ACG→TCG) | *ydbD* → | PF10971 family putative periplasmic methylglyoxa l resistance protein |
| 1,628,804 | G→A |  |  |  |  |  |  |  | 5.6% |  | A113A (GCC→GCT) | *tfaQ* ← | Qin prophage; putative tail fiber assembly prote in |
| 1,815,264 | C→G |  |  |  |  | 6.9% |  |  |  |  | A70P (GCC→CCC) | *chbC* ← | N,N'‑diacetylchitobiose‑specific enzyme IIC comp onent of PTS |
| 1,859,025 | T→A | 8.5% | 7.5% |  |  | 12.9% | 10.0% |  |  |  | intergenic (+34/+14) | *yeaD* → / ← *yeaE* | D‑hexose‑6‑phosphate epimerase‑like protein/aldo‑keto reductase, methylglyoxal to acetol, NA DPH‑dependent |
| 2,068,169 | C→A |  |  |  |  | 13.1% |  |  |  |  | intergenic (+30/‑91) | *flu* → / → *yeeR* | CP4‑44 prophage; antigen 43 (Ag43) phase‑variabl e biofilm formation autotransporter/CP4‑44 prophage; putative membrane protein |
| 2,135,510 | C→G |  |  |  |  | 7.2% |  |  |  |  | A63P (GCC→CCC) | *dcd* ← | 2'‑deoxycytidine 5'‑triphosphate deaminase |
| 2,156,362 | G→C |  | 10.9% |  |  |  |  |  |  |  | K2N (AAG→AAC) | *baeS* → | sensory histidine kinase in two‑component regula tory system with BaeR |
| 2,156,364 | T→A |  | 15.0% |  |  |  |  |  |  |  | F3Y (TTC→TAC) | *baeS* → | sensory histidine kinase in two‑component regula tory system with BaeR |
| 2,222,669 | T→A |  | 10.5% | 7.2% |  | 8.9% | 7.8% |  |  |  | intergenic (+125/+248) | *yohP* → / ← *dusC* | uncharacterized protein/tRNA‑dihydrouridine synthase C |
| 2,222,671 | G→C |  | 7.3% | 6.6% |  | 8.9% | 10.3% |  |  |  | intergenic (+127/+246) | *yohP* → / ← *dusC* | uncharacterized protein/tRNA‑dihydrouridine synthase C |
| 2,222,684 | G→C |  |  |  |  | 6.9% | 5.8% |  |  |  | intergenic (+140/+233) | *yohP* → / ← *dusC* | uncharacterized protein/tRNA‑dihydrouridine synthase C |
| 2,240,751 | G→C |  |  |  |  |  |  |  | 13.1% |  | P421A (CCG→GCG) | *lysP* ← | lysine transporter |
| 2,528,623 | G→C |  |  |  |  | 6.4% |  |  |  |  | R400P (CGT→CCT) | *ptsI* → | PEP‑protein phosphotransferase of PTS system (en zyme I) |
| 2,711,680 | G→C |  |  |  |  | 13.5% |  |  |  |  | R70G (CGC→GGC) | *yfiF* ← | putative methyltransferase |
| 2,718,215 | T→A |  |  |  |  | 5.7% |  |  |  |  | P297P (CCA→CCT) | *kgtP* ← | alpha‑ketoglutarate transporter |
| 2,724,354 | G→A |  |  |  |  |  |  |  |  | 14.0% | noncoding (163/1542 nt) | *rrsG* ← | 16S ribosomal RNA of rrnG operon |
| 2,931,054 | T→A |  |  |  |  |  |  | 5.7% |  |  | G86G (GGT→GGA) | *fucK* → | L‑fuculokinase |
| 2,959,548 | T→A |  | 6.5% |  |  |  |  |  |  |  | \*749L (TAG→TTG) | *ptsP* ← | fused PTS enzyme: PEP‑protein phosphotransferase (enzyme I)/GAF domain containing protein |
| 3,048,240 | T→G |  |  |  |  |  |  |  | 5.4% |  | K188N (AAA→AAC) | *ygfB* ← | UPF0149 family protein |
| 3,263,328 | A→C |  |  |  |  |  | 5.8% |  |  |  | intergenic (+367/+247) | *yhaC* → / ← *rnpB* | pentapetide repeats‑related protein/RNase P, M1 RNA component |
| 3,263,460 | C→A |  |  |  |  | 5.5% |  |  |  |  | intergenic (+499/+115) | *yhaC* → / ← *rnpB* | pentapetide repeats‑related protein/RNase P, M1 RNA component |
| 3,263,573 | T→A |  |  |  |  |  |  |  |  | 5.2% | intergenic (+612/+2) | *yhaC* → / ← *rnpB* | pentapetide repeats‑related protein/RNase P, M1 RNA component |
| 3,300,489 | T→A |  |  |  |  | 11.3% |  |  |  |  | D244V (GAT→GTT) | *deaD* ← | ATP‑dependent RNA helicase |
| 3,385,571 | C→G |  |  |  |  |  |  |  | 6.3% |  | intergenic (‑184/+246) | *tldD* ← / ← *yhdP* | putative peptidase/DUF3971‑AsmA2 domains protein |
| 3,433,818 | G→A |  |  |  |  |  |  | 38.1% |  |  | R191C (CGT→TGT) | *rpoA* ← | RNA polymerase, alpha subunit |
| 3,434,290 | +CCA |  |  |  | 6.6% |  |  |  |  |  | coding (99/990 nt) | *rpoA* ← | RNA polymerase, alpha subunit |
| 3,652,522 | A→T |  |  |  |  |  | 10.6% |  |  |  | intergenic (+269/‑70) | *gadE* → / → *mdtE* | gad regulon transcriptional activator/anaerobic multidrug efflux transporter, ArcA‑reg ulated |
| 3,652,527 | C→G |  | 6.3% |  |  |  |  |  |  |  | intergenic (+274/‑65) | *gadE* → / → *mdtE* | gad regulon transcriptional activator/anaerobic multidrug efflux transporter, ArcA‑reg ulated |
| 3,762,384 | A→T |  |  |  | 5.9% |  |  |  |  |  | pseudogene (50/282 nt) | *yibW* → | pseudogene, rhsA‑linked |
| 3,774,404 | T→A |  |  |  | 5.3% |  |  |  |  |  | intergenic (+27/‑171) | *lldD* → / → *trmL* | L‑lactate dehydrogenase, FMN‑linked/tRNA Leu mC34,mU34 2'‑O‑methyltransferase, SAM‑d ependent |
| 3,774,416 | A→T |  |  |  | 6.1% |  |  |  |  |  | intergenic (+39/‑159) | *lldD* → / → *trmL* | L‑lactate dehydrogenase, FMN‑linked/tRNA Leu mC34,mU34 2'‑O‑methyltransferase, SAM‑d ependent |
| 3,774,418 | A→T | 6.7% |  |  | 6.0% | 6.5% |  | 6.3% |  | 6.2% | intergenic (+41/‑157) | *lldD* → / → *trmL* | L‑lactate dehydrogenase, FMN‑linked/tRNA Leu mC34,mU34 2'‑O‑methyltransferase, SAM‑d ependent |
| 3,774,455 | G→T |  |  |  |  |  | 6.3% |  |  |  | intergenic (+78/‑120) | *lldD* → / → *trmL* | L‑lactate dehydrogenase, FMN‑linked/tRNA Leu mC34,mU34 2'‑O‑methyltransferase, SAM‑d ependent |
| 3,774,466 | G→C |  |  |  |  |  |  | 6.6% |  |  | intergenic (+89/‑109) | *lldD* → / → *trmL* | L‑lactate dehydrogenase, FMN‑linked/tRNA Leu mC34,mU34 2'‑O‑methyltransferase, SAM‑d ependent |
| 3,774,471 | C→G |  |  |  |  | 6.7% |  |  |  |  | intergenic (+94/‑104) | *lldD* → / → *trmL* | L‑lactate dehydrogenase, FMN‑linked/tRNA Leu mC34,mU34 2'‑O‑methyltransferase, SAM‑d ependent |
| 3,774,480 | T→A | 5.1% |  |  |  |  |  |  |  |  | intergenic (+103/‑95) | *lldD* → / → *trmL* | L‑lactate dehydrogenase, FMN‑linked/tRNA Leu mC34,mU34 2'‑O‑methyltransferase, SAM‑d ependent |
| 3,774,500 | A→T | 8.8% |  | 8.9% | 8.9% | 6.9% | 7.1% | 6.6% |  | 8.6% | intergenic (+123/‑75) | *lldD* → / → *trmL* | L‑lactate dehydrogenase, FMN‑linked/tRNA Leu mC34,mU34 2'‑O‑methyltransferase, SAM‑d ependent |
| 3,774,505 | C→G |  |  |  |  | 9.3% | 12.9% |  |  |  | intergenic (+128/‑70) | *lldD* → / → *trmL* | L‑lactate dehydrogenase, FMN‑linked/tRNA Leu mC34,mU34 2'‑O‑methyltransferase, SAM‑d ependent |
| 3,774,512 | A→T | 6.5% |  |  |  | 5.8% | 7.4% |  |  |  | intergenic (+135/‑63) | *lldD* → / → *trmL* | L‑lactate dehydrogenase, FMN‑linked/tRNA Leu mC34,mU34 2'‑O‑methyltransferase, SAM‑d ependent |
| 3,809,170 | Δ1 bp |  |  | 16.1% | 10.8% | 25.0% |  |  |  |  | intergenic (‑42/+24) | *pyrE* ← / ← *rph* | orotate phosphoribosyltransferase/ribonuclease PH (defective);enzyme; Degradation of RNA; RNase PH |
| 3,809,170 | G→T | 29.5% |  | Δ | Δ | Δ |  |  |  |  | intergenic (‑42/+24) | *pyrE* ← / ← *rph* | orotate phosphoribosyltransferase/ribonuclease PH (defective);enzyme; Degradation of RNA; RNase PH |
| 3,809,183 | C→T |  |  |  |  | 7.3% |  |  |  |  | intergenic (‑55/+11) | *pyrE* ← / ← *rph* | orotate phosphoribosyltransferase/ribonuclease PH (defective);enzyme; Degradation of RNA; RNase PH |
| 3,809,258:1 | +T | 6.6% |  |  |  |  |  |  |  |  | pseudogene (652/669 nt) | *rph* ← | ribonuclease PH (defective);enzyme; Degradation of RNA; RNase PH |
| 3,830,623 | A→T |  |  | 6.6% |  |  |  |  |  |  | Y104F (TAC→TTC) | *setC* → | putative arabinose efflux transporter |
| 3,952,834 | T→A |  | 5.4% |  |  | 6.9% | 5.3% |  |  |  | intergenic (+29/+58) | *ilvC* → / ← *ppiC* | ketol‑acid reductoisomerase, NAD(P)‑binding/peptidyl‑prolyl cis‑trans isomerase C (rotamase C) |
| 3,952,870 | T→A |  |  |  |  | 6.1% |  |  |  |  | intergenic (+65/+22) | *ilvC* → / ← *ppiC* | ketol‑acid reductoisomerase, NAD(P)‑binding/peptidyl‑prolyl cis‑trans isomerase C (rotamase C) |
| 3,998,693 | A→T |  | 6.7% |  |  |  |  |  |  |  | M158L (ATG→TTG) | *pldA* → | outer membrane phospholipase A |
| 3,998,699 | T→A |  |  |  |  | 5.9% |  |  |  |  | Y160N (TAT→AAT) | *pldA* → | outer membrane phospholipase A |
| 4,080,323 | A→T | 9.6% |  |  |  |  |  |  |  |  | intergenic (+114/‑39) | *fdhD* → / → *yiiG* | formate dehydrogenase formation protein/DUF3829 family lipoprotein |
| 4,081,456 | T→A | 6.5% | 5.3% |  |  | 5.7% |  |  |  | 8.5% | intergenic (+39/+11) | *yiiG* → / ← *frvR* | DUF3829 family lipoprotein/putative frv operon regulator; contains a PTS EI IA domain |
| 4,171,585 | (CAACGGTACCTTTGTTAT)_1→2_ |  |  |  | 9.1% |  | 8.7% |  |  |  | coding (413/4029 nt) | *rpoB* → | RNA polymerase, beta subunit |
| 4,171,618 | T→G |  | 75.5% |  |  |  |  |  |  |  | L149R (CTG→CGG) | *rpoB* → | RNA polymerase, beta subunit |
| 4,173,034 | C→T |  |  |  |  |  |  | 18.8% |  |  | S621F (TCC→TTC) | *rpoB* → | RNA polymerase, beta subunit |
| 4,173,082 | G→T |  |  |  |  |  |  |  | 86.0% |  | R637L (CGT→CTT) | *rpoB* → | RNA polymerase, beta subunit |
| 4,173,162 | G→C |  |  |  |  |  |  | 13.9% |  |  | G664R (GGT→CGT) | *rpoB* → | RNA polymerase, beta subunit |
| 4,176,035 | (TCCGCTGGT)_1→2_ |  |  |  | 10.8% |  | 47.6% |  | 9.2% | 94.5% | coding (758/4224 nt) | *rpoC* → | RNA polymerase, beta prime subunit |
| 4,178,889 | (GAACGTGTA)_1→2_ |  |  |  |  |  |  | 16.2% |  |  | coding (3612/4224 nt) | *rpoC* → | RNA polymerase, beta prime subunit |
| 4,378,486 | C→A | 14.7% |  |  |  |  |  |  |  |  | Q234H (CAG→CAT) | *mscM* ← | mechanosensitive channel protein, miniconductanc e |
| 4,388,187 | A→T |  |  |  |  |  |  | 7.9% |  |  | D320V (GAT→GTT) | *mutL* → | methyl‑directed mismatch repair protein |
