## Supplementary material for "Osmotic conditions shape fitness gains and resistance mechanisms during *E. coli* and T4 phage co-evolution": Table S2: Table_S2_CoBac.html

Mutation Comparison


| Predicted mutations | | | | | | | | | | | | | |
| --- | --- | --- | --- | --- | --- | --- | --- | --- | --- | --- | --- | --- | --- |
| position | mutation | L1_36_ | L2_13_ | L3_16_ | M1_9_ | M2_9_ | M3_36_ | H1_36_ | H2_36_ | H3_36_ | annotation | gene | description |
| 104,042 | A→T |  |  |  |  |  | 8.8% |  |  |  | intergenic (+81/‑150) | *lpxC* → / → *secM* | UDP‑3‑O‑acyl N‑acetylglucosamine deacetylase/regulator of secA translation |
| 104,054 | G→T |  | 14.5% |  |  |  |  |  |  |  | intergenic (+93/‑138) | *lpxC* → / → *secM* | UDP‑3‑O‑acyl N‑acetylglucosamine deacetylase/regulator of secA translation |
| 157,963 | G→C |  |  |  |  |  |  | 5.6% |  |  | S4C (TCT→TGT) | *sfsA* ← | sugar fermentation stimulation protein A |
| 161,051 | A→T |  | 10.4% |  |  | 13.3% | 5.8% |  |  |  | intergenic (+30/‑166) | *hrpB* → / → *mrcB* | putative ATP‑dependent helicase/fused glycosyl transferase and transpeptidase |
| 284,974 | A→T |  |  |  |  |  | 5.6% |  |  |  | intergenic (‑101/+38) | *yagI* ← / ← *argF* | CP4‑6 prophage; putative DNA‑binding transcripti onal regulator/ornithine carbamoyltransferase 2, chain F; CP4‑6 prophage |
| 375,461 | A→C | 6.5% |  |  |  |  |  |  |  |  | intergenic (‑124/+64) | *frmR* ← / ← *yaiO* | regulator protein that represses frmRAB operon/outer membrane protein |
| 375,802 | G→A |  |  |  |  |  |  |  | 31.3% |  | A166V (GCA→GTA) | *yaiO* ← | outer membrane protein |
| 481,346 | A→T |  |  |  | 5.1% |  |  |  |  |  | T44S (ACG→TCG) | *acrR* → | transcriptional repressor |
| 635,028 | G→T |  |  |  |  |  | 5.5% |  |  |  | intergenic (+64/‑181) | *ahpC* → / → *ahpF* | alkyl hydroperoxide reductase, C22 subunit/alkyl hydroperoxide reductase, F52a subunit, FAD /NAD(P)‑binding |
| 635,034 | G→T |  |  |  |  |  | 5.9% |  |  |  | intergenic (+70/‑175) | *ahpC* → / → *ahpF* | alkyl hydroperoxide reductase, C22 subunit/alkyl hydroperoxide reductase, F52a subunit, FAD /NAD(P)‑binding |
| 642,062 | C→A |  |  |  |  |  |  | 22.5% |  |  | intergenic (‑26/+25) | *citT* ← / ← *citG* | citrate:succinate antiporter/2‑(5''‑triphosphoribosyl)‑3'‑dephosphocoenzyme‑A synthase |
| 703,445 | C→A |  | 7.0% |  |  |  |  |  |  |  | intergenic (+232/‑345) | *glnS* → / → *chiP* | glutamyl‑tRNA synthetase/chitoporin, uptake of chitosugars |
| 780,477 | G→C | 5.9% |  |  |  |  |  |  |  |  | R99R (CGC→CGG) | *ybgS* ← | putative periplasmic protein |
| 910,654 | C→G | 6.2% |  |  | 7.5% |  | 5.7% |  |  | 6.5% | intergenic (‑341/+154) | *ybjE* ← / ← *aqpZ* | putative transporter/aquaporin Z |
| 910,667 | C→G |  |  |  | 7.3% |  |  |  |  | 6.1% | intergenic (‑354/+141) | *ybjE* ← / ← *aqpZ* | putative transporter/aquaporin Z |
| 910,669 | A→T | 12.4% |  | 6.7% | 9.3% |  | 10.1% | 12.3% | 7.0% | 11.4% | intergenic (‑356/+139) | *ybjE* ← / ← *aqpZ* | putative transporter/aquaporin Z |
| 910,672 | G→T |  |  |  |  |  | 10.7% |  |  |  | intergenic (‑359/+136) | *ybjE* ← / ← *aqpZ* | putative transporter/aquaporin Z |
| 1,423,022 | T→A | 21.8% |  |  |  |  |  |  |  |  | intergenic (‑39/‑30) | *insH1* ← / → *lomR* | IS5 transposase and trans‑activator;IS, phage, T n; Transposon‑related functions; extrachromosomal; transpo son related/pseudogene, Rac prophage lom homolog;Phage or Pr ophage Related; interrupted by IS5 and N‑ter deletion |
| 1,538,659 | G→C |  | 7.9% |  |  |  |  |  |  |  | pseudogene (363/381 nt) | *yddK* ← | pseudogene, leucine‑rich protein; putative glyco portein |
| 1,672,825 | G→T |  |  |  | 8.0% |  |  |  |  | 7.8% | A48S (GCT→TCT) | *ydgH* → | DUF1471 family periplasmic protein |
| 1,714,610 | C→G |  |  |  |  |  | 9.5% |  |  |  | intergenic (+10/+37) | *slyB* → / ← *slyA* | outer membrane lipoprotein/global transcriptional regulator |
| 1,718,605 | C→G | 15.4% |  |  |  |  |  |  |  |  | G103A (GGG→GCG) | *sodC* ← | superoxide dismutase, Cu, Zn, periplasmic |
| 1,859,012 | C→G |  |  |  |  |  | 7.9% |  |  |  | intergenic (+21/+27) | *yeaD* → / ← *yeaE* | D‑hexose‑6‑phosphate epimerase‑like protein/aldo‑keto reductase, methylglyoxal to acetol, NA DPH‑dependent |
| 1,859,025 | T→A |  | 8.7% | 11.8% |  | 13.8% | 10.7% |  |  |  | intergenic (+34/+14) | *yeaD* → / ← *yeaE* | D‑hexose‑6‑phosphate epimerase‑like protein/aldo‑keto reductase, methylglyoxal to acetol, NA DPH‑dependent |
| 1,922,058 | T→A |  |  |  |  |  |  |  |  | 7.2% | T347S (ACC→TCC) | *ptrB* ← | protease II |
| 2,017,304 | C→T |  |  |  |  |  | 58.4% | 46.9% |  | 100% | intergenic (+145/‑145) | *fliR* → / → *rcsA* | flagellar export pore protein/transcriptional regulator of colanic acid capsul ar biosynthesis |
| 2,017,308 | T→C |  |  |  |  |  |  |  | 100% |  | intergenic (+149/‑141) | *fliR* → / → *rcsA* | flagellar export pore protein/transcriptional regulator of colanic acid capsul ar biosynthesis |
| 2,017,986 | A→T |  |  |  |  |  |  | 51.5% |  |  | I180F (ATC→TTC) | *rcsA* → | transcriptional regulator of colanic acid capsul ar biosynthesis |
| 2,156,364 | T→A |  |  |  | 13.4% |  |  |  |  |  | F3Y (TTC→TAC) | *baeS* → | sensory histidine kinase in two‑component regula tory system with BaeR |
| 2,222,669 | T→A |  | 8.7% | 7.0% | 9.7% | 9.2% | 9.4% |  | 12.7% |  | intergenic (+125/+248) | *yohP* → / ← *dusC* | uncharacterized protein/tRNA‑dihydrouridine synthase C |
| 2,222,671 | G→C |  | 6.3% |  | 8.5% |  |  |  |  |  | intergenic (+127/+246) | *yohP* → / ← *dusC* | uncharacterized protein/tRNA‑dihydrouridine synthase C |
| 2,222,684 | G→C |  |  | 5.1% |  |  |  |  |  |  | intergenic (+140/+233) | *yohP* → / ← *dusC* | uncharacterized protein/tRNA‑dihydrouridine synthase C |
| 2,305,126 | T→A |  |  | 10.0% |  |  |  |  |  |  | \*368L (TAA→TTA) | *ompC* ← | outer membrane porin protein C |
| 2,305,627 | Δ6 bp |  | 100% |  |  |  |  |  |  |  | coding (597‑602/1104 nt) | *ompC* ← | outer membrane porin protein C |
| 2,305,645 | C→G | 100% |  | 87.0% | 100% | 100% | 66.8% | 51.5% |  |  | R195P (CGT→CCT) | *ompC* ← | outer membrane porin protein C |
| 2,305,782 | Δ1 bp |  |  |  |  |  | 30.6% |  |  |  | coding (447/1104 nt) | *ompC* ← | outer membrane porin protein C |
| 2,305,783 | Δ1 bp |  |  |  |  |  | 30.3% |  |  |  | coding (446/1104 nt) | *ompC* ← | outer membrane porin protein C |
| 2,459,789 | T→A | 18.5% |  |  |  |  |  |  |  |  | noncoding (2/75 nt) | *argW* → | tRNA‑Arg |
| 2,494,198 | C→A |  |  |  |  |  | 12.1% |  |  |  | Q113K (CAA→AAA) | *ypdB* → | response regulator activating yhjX; pyruvate‑res ponsive YpdAB two‑component system |
| 2,554,625 | C→G |  |  |  | 6.0% |  |  |  |  |  | intergenic (+368/‑102) | *yffL* → / → *yffM* | CPZ‑55 prophage; uncharacterized protein/CPZ‑55 prophage; uncharacterized protein |
| 2,637,659 | G→T |  |  |  |  |  |  |  | 7.3% |  | intergenic (‑17/+133) | *rlmN* ← / ← *ndk* | dual specificity 23S rRNA m(2)A2503, tRNA m(2)A3 7 methyltransferase, SAM‑dependent/multifunctional nucleoside diphosphate kinase an d apyrimidinic endonuclease and 3'‑phosphodiesterase |
| 2,709,569 | G→A |  |  |  |  |  | 5.9% |  |  | 7.6% | R80R (CGC→CGT) | *grcA* ← | autonomous glycyl radical cofactor |
| 2,724,312 | T→C | 100% |  | 87.2% | 100% | 88.1% | 64.8% | 36.8% |  |  | noncoding (205/1542 nt) | *rrsG* ← | 16S ribosomal RNA of rrnG operon |
| 2,799,636 | (TTCCAGGGCGTGCGCGTTCCGG)_1→2_ |  |  |  |  |  |  |  |  | 6.5% | coding (268/1065 nt) | *proW* → | glycine betaine transporter subunit |
| 2,799,794 | G→A |  |  |  |  |  |  | 28.5% |  |  | W142\* (TGG→TGA) | *proW* → | glycine betaine transporter subunit |
| 2,860,448 | Δ1 bp | 100% |  |  |  |  |  |  |  |  | coding (463/993 nt) | *rpoS* ← | RNA polymerase, sigma S (sigma 38) factor |
| 3,048,247 | A→T |  |  | 6.7% |  |  |  |  |  |  | V186E (GTA→GAA) | *ygfB* ← | UPF0149 family protein |
| 3,050,051 | A→T |  |  |  |  |  | 5.4% |  |  |  | Y151F (TAT→TTT) | *fau* → | 5‑formyltetrahydrofolate cyclo‑ligase family pro tein |
| 3,086,401 | G→A |  |  |  |  |  | 7.8% |  |  |  | G36R (GGG→AGG) | *yqgE* → | uncharacterized protein |
| 3,232,961 | T→A |  |  |  |  |  |  |  | 7.2% |  | intergenic (+57/‑342) | *alx* → / → *sstT* | putative membrane‑bound redox modulator/sodium:serine/threonine symporter |
| 3,263,408 | T→A | 6.6% |  |  |  |  |  |  |  |  | intergenic (+447/+167) | *yhaC* → / ← *rnpB* | pentapetide repeats‑related protein/RNase P, M1 RNA component |
| 3,263,502 | C→T |  |  |  |  |  |  | 6.0% |  |  | intergenic (+541/+73) | *yhaC* → / ← *rnpB* | pentapetide repeats‑related protein/RNase P, M1 RNA component |
| 3,385,662 | C→G |  |  |  |  | 6.7% |  |  |  |  | intergenic (‑275/+155) | *tldD* ← / ← *yhdP* | putative peptidase/DUF3971‑AsmA2 domains protein |
| 3,404,676 | IS*5* (+) +4 bp |  |  |  |  |  |  |  |  | 10.0% | coding (47‑50/297 nt) | *fis* → | global DNA‑binding transcriptional dual regulato r |
| 3,404,765 | Δ1 bp |  |  |  |  |  |  |  | 32.3% |  | coding (136/297 nt) | *fis* → | global DNA‑binding transcriptional dual regulato r |
| 3,404,861 | G→A |  |  |  |  |  |  |  |  | 18.2% | A78T (GCG→ACG) | *fis* → | global DNA‑binding transcriptional dual regulato r |
| 3,404,881 | C→A |  |  |  |  |  |  |  | 5.2% |  | N84K (AAC→AAA) | *fis* → | global DNA‑binding transcriptional dual regulato r |
| 3,596,944 | A→T |  | 5.8% |  |  |  |  |  |  |  | G220G (GGT→GGA) | *ftsY* ← | Signal Recognition Particle (SRP) receptor |
| 3,634,171 | T→A |  |  |  |  | 5.9% |  |  |  |  | intergenic (+266/‑51) | *uspA* → / → *dtpB* | universal stress global response regulator/dipeptide and tripeptide permease B |
| 3,653,036 | G→C |  | 11.8% |  |  |  |  |  |  |  | A149P (GCA→CCA) | *mdtE* → | anaerobic multidrug efflux transporter, ArcA‑reg ulated |
| 3,656,700 | C→G |  |  | 5.1% |  |  |  |  |  |  | T976S (ACC→AGC) | *mdtF* → | anaerobic multidrug efflux transporter, ArcA‑reg ulated |
| 3,774,418 | A→T |  | 7.1% |  | 5.7% |  | 6.0% | 5.7% |  |  | intergenic (+41/‑157) | *lldD* → / → *trmL* | L‑lactate dehydrogenase, FMN‑linked/tRNA Leu mC34,mU34 2'‑O‑methyltransferase, SAM‑d ependent |
| 3,774,438 | A→T | 5.7% |  |  |  |  |  |  |  |  | intergenic (+61/‑137) | *lldD* → / → *trmL* | L‑lactate dehydrogenase, FMN‑linked/tRNA Leu mC34,mU34 2'‑O‑methyltransferase, SAM‑d ependent |
| 3,774,455 | G→T |  |  |  | 5.2% |  |  |  |  |  | intergenic (+78/‑120) | *lldD* → / → *trmL* | L‑lactate dehydrogenase, FMN‑linked/tRNA Leu mC34,mU34 2'‑O‑methyltransferase, SAM‑d ependent |
| 3,774,466 | G→C | 5.0% |  |  |  |  |  |  |  |  | intergenic (+89/‑109) | *lldD* → / → *trmL* | L‑lactate dehydrogenase, FMN‑linked/tRNA Leu mC34,mU34 2'‑O‑methyltransferase, SAM‑d ependent |
| 3,774,500 | A→T |  | 6.9% | 6.5% | 7.4% |  | 6.9% |  |  |  | intergenic (+123/‑75) | *lldD* → / → *trmL* | L‑lactate dehydrogenase, FMN‑linked/tRNA Leu mC34,mU34 2'‑O‑methyltransferase, SAM‑d ependent |
| 3,774,505 | C→G |  |  |  | 12.5% |  |  | 11.3% |  |  | intergenic (+128/‑70) | *lldD* → / → *trmL* | L‑lactate dehydrogenase, FMN‑linked/tRNA Leu mC34,mU34 2'‑O‑methyltransferase, SAM‑d ependent |
| 3,795,449 | IS*5* (+) +4 bp |  |  |  | 10.9% |  |  |  |  |  | coding (967‑970/1020 nt) | *waaR* ← | UDP‑D‑galactose:(glucosyl)lipopolysaccharide‑ al pha‑1,3‑D‑galactosyltransferase |
| 3,795,481 | G→T |  |  |  |  | 29.2% |  |  |  |  | A313E (GCA→GAA) | *waaR* ← | UDP‑D‑galactose:(glucosyl)lipopolysaccharide‑ al pha‑1,3‑D‑galactosyltransferase |
| 3,795,502 | Δ1 bp |  | 15.4% |  |  |  |  |  |  |  | coding (917/1020 nt) | *waaR* ← | UDP‑D‑galactose:(glucosyl)lipopolysaccharide‑ al pha‑1,3‑D‑galactosyltransferase |
| 3,795,902 | IS*5* (–) +4 bp |  |  | 10.8% | 11.6% |  |  |  |  |  | coding (514‑517/1020 nt) | *waaR* ← | UDP‑D‑galactose:(glucosyl)lipopolysaccharide‑ al pha‑1,3‑D‑galactosyltransferase |
| 3,795,902 | IS*5* (+) +4 bp |  |  |  |  | 24.8% |  |  |  |  | coding (514‑517/1020 nt) | *waaR* ← | UDP‑D‑galactose:(glucosyl)lipopolysaccharide‑ al pha‑1,3‑D‑galactosyltransferase |
| 3,796,012 | C→T |  |  |  | 13.3% |  |  |  |  |  | C136Y (TGT→TAT) | *waaR* ← | UDP‑D‑galactose:(glucosyl)lipopolysaccharide‑ al pha‑1,3‑D‑galactosyltransferase |
| 3,796,024 | Δ12 bp |  |  |  | 21.5% |  |  |  |  |  | coding (384‑395/1020 nt) | *waaR* ← | UDP‑D‑galactose:(glucosyl)lipopolysaccharide‑ al pha‑1,3‑D‑galactosyltransferase |
| 3,796,093 | G→T |  |  |  |  | 6.0% |  |  |  |  | A109E (GCA→GAA) | *waaR* ← | UDP‑D‑galactose:(glucosyl)lipopolysaccharide‑ al pha‑1,3‑D‑galactosyltransferase |
| 3,796,114 | IS*5* (–) +4 bp |  |  |  | 11.3% |  |  |  |  |  | coding (302‑305/1020 nt) | *waaR* ← | UDP‑D‑galactose:(glucosyl)lipopolysaccharide‑ al pha‑1,3‑D‑galactosyltransferase |
| 3,796,114 | IS*5* (+) +4 bp |  |  |  |  | 11.8% |  |  |  |  | coding (302‑305/1020 nt) | *waaR* ← | UDP‑D‑galactose:(glucosyl)lipopolysaccharide‑ al pha‑1,3‑D‑galactosyltransferase |
| 3,796,397 | C→A |  |  |  |  |  | 29.5% |  |  |  | E8\* (GAA→TAA) | *waaR* ← | UDP‑D‑galactose:(glucosyl)lipopolysaccharide‑ al pha‑1,3‑D‑galactosyltransferase |
| 3,799,422 | IS*5* (–) +4 bp |  |  | 83.2% |  |  |  |  |  |  | coding (1003‑1006/1125 nt) | *waaG* ← | glucosyltransferase I |
| 3,896,169 | C→A |  |  |  |  |  | 5.1% |  |  |  | S298I (AGT→ATT) | *bglB* ← | cryptic phospho‑beta‑glucosidase B |
| 3,952,834 | T→A |  |  | 6.1% |  |  | 8.5% |  | 7.1% |  | intergenic (+29/+58) | *ilvC* → / ← *ppiC* | ketol‑acid reductoisomerase, NAD(P)‑binding/peptidyl‑prolyl cis‑trans isomerase C (rotamase C) |
| 3,998,685 | A→T |  |  |  |  |  | 12.0% |  |  |  | D155V (GAT→GTT) | *pldA* → | outer membrane phospholipase A |
| 3,998,693 | A→T |  |  | 9.8% |  |  |  |  | 9.4% |  | M158L (ATG→TTG) | *pldA* → | outer membrane phospholipase A |
| 4,081,456 | T→A |  |  | 6.2% | 6.5% |  |  |  | 8.8% | 8.4% | intergenic (+39/+11) | *yiiG* → / ← *frvR* | DUF3829 family lipoprotein/putative frv operon regulator; contains a PTS EI IA domain |
| 4,225,570 | T→C | 8.1% |  |  |  |  |  |  |  |  | intergenic (+235/‑264) | *pgi* → / → *yjbE* | glucosephosphate isomerase/extracellular polysaccharide production threonin e‑rich protein |
| 4,226,049 | Δ9 bp | 6.9% |  |  |  |  |  |  |  |  | coding (216‑224/243 nt) | *yjbE* → | extracellular polysaccharide production threonin e‑rich protein |
| 4,226,981 | Δ1 bp |  |  |  |  |  | 18.5% |  |  |  | coding (157/738 nt) | *yjbG* → | extracellular polysaccharide export OMA protein |
| 4,227,826 | C→T | 9.2% |  |  |  |  |  |  |  |  | Q89\* (CAG→TAG) | *yjbH* → | DUF940 family extracellular polysaccharide prote in |
| 4,227,938:1 | +G |  |  |  |  |  | 15.6% |  |  |  | coding (377/2097 nt) | *yjbH* → | DUF940 family extracellular polysaccharide prote in |
| 4,388,187 | A→T |  |  |  | 8.6% |  |  |  |  |  | D320V (GAT→GTT) | *mutL* → | methyl‑directed mismatch repair protein |
| 4,485,076 | A→T |  |  |  |  |  | 5.3% |  |  |  | I317I (ATT→ATA) | *ahr* ← | aldehyde reductase, NADPH‑dependent, Zn‑containi ng, broad specificity |
