## Supplementary material for "Osmotic conditions shape fitness gains and resistance mechanisms during *E. coli* and T4 phage co-evolution": Table S3: Table_S3_IndPhage.html

Mutation Comparison


| Predicted mutations | | | | | | | | | | | | | |
| --- | --- | --- | --- | --- | --- | --- | --- | --- | --- | --- | --- | --- | --- |
| position | mutation | L1_36_ | L2_36_ | L3_36_ | M1_36_ | M2_36_ | M3_36_ | H1_36_ | H2_36_ | H3_36_ | annotation | gene | description |
| 316 | T→A |  | 8.2% |  |  |  |  |  |  |  | pseudogene (1874/2178 nt) | *rIIA* ← | RIIA lysis inhibitor |
| 8,515 | Δ1 bp |  |  | 6.6% |  |  |  |  |  |  | coding (380/684 nt) | *dexA* ← | exonuclease |
| 8,516 | Δ1 bp |  |  | 6.6% |  |  |  |  |  |  | coding (379/684 nt) | *dexA* ← | exonuclease |
| 8,917 | G→A |  |  | 6.8% |  |  |  |  |  |  | R74C (CGT→TGT) | *dexA.1* ← | DexA.1 hypothetical protein |
| 10,948 | A→G |  | 14.6% |  |  |  |  |  | 12.9% |  | S26P (TCT→CCT) | *dda.1* ← | Dda.1 hypothetical protein |
| 17,718 | Δ1 bp | 11.2% |  | 70.5% | 10.5% | 11.6% |  | 51.2% |  |  | intergenic (‑107/+203) | *dam* ← / ← *61* | Dam family site‑specific DNA‑(adenine‑N6)‑methyl transferase/DNA primase |
| 17,719 | Δ1 bp | 9.9% |  | 63.7% | 8.8% | 9.3% |  | 48.3% |  |  | intergenic (‑108/+202) | *dam* ← / ← *61* | Dam family site‑specific DNA‑(adenine‑N6)‑methyl transferase/DNA primase |
| 34,030 | A→T |  | 5.5% |  |  |  |  |  | 9.5% |  | H314Q (CAT→CAA) | *46* ← | SbcC‑like subunit of palindrome specific endonuc lease |
| 44,347 | C→T | 5.2% |  |  |  |  |  |  |  |  | Q331Q (CAG→CAA) | *nrdD* ← | anaerobic ribonucleoside reductase large subunit |
| 46,390 | C→T |  | 9.7% |  |  |  |  |  |  |  | Q99Q (CAG→CAA)  Q151Q (CAG→CAA) | *49* ← *49* ← | endonuclease VII endonuclease VII |
| 57,896 | Δ1 bp | 5.4% |  |  |  |  |  |  |  |  | coding (24/105 nt) | *mobD.2* ← | MobD.2 conserved hypothetical protein |
| 57,897 | Δ1 bp | 5.3% |  |  |  |  |  |  |  |  | coding (23/105 nt) | *mobD.2* ← | MobD.2 conserved hypothetical protein |
| 58,278 | A→G |  |  |  |  |  |  | 5.6% |  |  | D20D (GAT→GAC) | *mobD.3* ← | MobD.3 hypothetical protein |
| 59,453 | T→C |  |  |  | 12.2% |  |  |  |  |  | M11V (ATG→GTG) | *rI* ← | lysis inhibition |
| 63,963 | G→A | 13.0% |  |  |  |  |  |  |  | 7.0% | P95S (CCA→TCA) | *vs.7* ← | hypothetical protein |
| 73,450 | G→T |  |  |  |  |  |  |  | 27.5% |  | S15Y (TCT→TAT) | *trna.4* ← | hypothetical protein |
| 78,491 | G→A |  |  |  | 7.0% | 19.8% | 7.7% |  |  |  | G185E (GGA→GAA) | *5* → | baseplate hub subunit and tail lysozyme |
| 83,923 | C→T |  | 12.4% |  |  |  |  |  | 20.5% |  | A295V (GCC→GTC) | *7* → | baseplate wedge subunit |
| 92,138 | A→G |  |  |  |  |  |  |  |  | 6.0% | P10P (CCA→CCG) | *wac* → | fibritin neck whisker |
| 92,141 | C→T |  |  |  |  |  |  | 6.7% | 7.3% | 8.3% | F11F (TTC→TTT) | *wac* → | fibritin neck whisker |
| 92,587 | C→T |  |  | 14.8% |  |  | 11.5% |  |  |  | T160I (ACT→ATT) | *wac* → | fibritin neck whisker |
| 92,668 | T→C | 47.8% |  | 16.1% |  |  | 16.4% |  |  |  | V187A (GTC→GCC) | *wac* → | fibritin neck whisker |
| 92,681 | G→T | 9.5% | 72.6% |  |  |  |  |  |  |  | Q191H (CAG→CAT) | *wac* → | fibritin neck whisker |
| 92,811 | C→T |  |  | 16.3% |  |  |  |  |  |  | R235C (CGT→TGT) | *wac* → | fibritin neck whisker |
| 92,854 | A→G |  |  |  |  |  |  |  |  | 5.5% | D249G (GAC→GGC) | *wac* → | fibritin neck whisker |
| 92,959 | Δ6 bp |  | 5.1% |  |  |  |  |  |  |  | coding (851‑856/1464 nt) | *wac* → | fibritin neck whisker |
| 93,129 | T→C | 10.7% | 8.1% |  |  |  |  |  |  |  | W341R (TGG→CGG) ‡ | *wac* → | fibritin neck whisker |
| 93,130 | G→T |  | 49.5% |  |  |  |  |  |  |  | W341L (TGG→TTG) ‡ | *wac* → | fibritin neck whisker |
| 97,948 | A→G |  |  |  | 5.6% |  |  |  |  |  | T433A (ACT→GCT)  T346A (ACT→GCT)  T328A (ACT→GCT)  T239A (ACT→GCT) | *17* → *17* → *17* → *17* → | terminase large subunit terminase large subunit terminase large subunit large terminase protein |
| 98,877 | T→C |  | 8.8% |  |  |  |  |  |  |  | pseudogene (362/1980 nt) | *18* → | tail sheath |
| 99,017 | A→G |  | 8.9% |  |  |  |  |  |  |  | pseudogene (502/1980 nt) | *18* → | tail sheath |
| 99,069 | G→A |  |  | 5.0% |  |  |  |  |  |  | pseudogene (554/1980 nt) | *18* → | tail sheath |
| 100,398 | A→G |  |  |  |  |  |  |  |  | 11.2% | pseudogene (1883/1980 nt) | *18* → | tail sheath |
| 100,894 | C→A |  |  |  |  |  |  |  | 6.1% |  | Q95K (CAA→AAA) | *19* → | tail protein |
| 103,256 | G→A |  |  | 5.2% |  |  |  |  |  |  | R85H (CGC→CAC) | *68* → | head scaffolding protein |
| 105,273 | A→G |  | 33.9% |  | 24.2% | 49.2% | 72.6% |  | 16.8% |  | M117V (ATG→GTG) ‡ | *23* → | major head protein |
| 105,275 | G→A | 74.7% | 56.8% | 86.4% | 63.8% | 36.9% | 18.5% | 76.6% | 62.1% | 75.7% | M117I (ATG→ATA) ‡ | *23* → | major head protein |
| 105,276 | A→G |  |  |  | 6.5% |  |  |  | 18.7% | 12.0% | N118D (AAC→GAC) | *23* → | major head protein |
| 117,392 | T→C |  |  |  |  |  |  | 17.2% | 17.6% | 21.3% | M51T (ATG→ACG) ‡ | *27* → | baseplate hub |
| 117,393 | G→A |  |  |  |  |  |  | 11.2% | 11.7% | 13.4% | M51I (ATG→ATA) ‡ | *27* → | baseplate hub |
| 134,981 | C→G |  |  |  | 26.4% | 9.2% | 52.0% |  |  |  | G58R (GGT→CGT) | *pseT.1* ← | PseT.1 conserved hypothetical protein |
| 136,145 | T→C |  |  |  |  |  |  |  | 5.6% |  | T50A (ACC→GCC) | *alc* ← | inhibitor of host transcription |
| 136,416 | A→G |  |  | 10.8% |  |  |  |  |  |  | C356R (TGT→CGT) | *rnlA* ← | RNA ligase and tail fiber protein attachment cat alyst |
| 136,509 | C→A |  |  |  | 67.3% |  |  |  |  |  | A325S (GCA→TCA) | *rnlA* ← | RNA ligase and tail fiber protein attachment cat alyst |
| 136,520 | T→C |  |  | 8.2% |  |  |  |  |  |  | Y321C (TAT→TGT) | *rnlA* ← | RNA ligase and tail fiber protein attachment cat alyst |
| 136,559 | T→C | 20.7% |  |  |  |  |  |  |  |  | D308G (GAC→GGC) | *rnlA* ← | RNA ligase and tail fiber protein attachment cat alyst |
| 136,560 | C→T |  |  | 45.5% |  |  |  |  |  |  | D308N (GAC→AAC) | *rnlA* ← | RNA ligase and tail fiber protein attachment cat alyst |
| 136,760 | A→T |  |  |  |  |  |  |  |  | 21.0% | I241N (ATT→AAT) | *rnlA* ← | RNA ligase and tail fiber protein attachment cat alyst |
| 138,429 | G→A |  |  |  | 5.5% |  |  |  |  |  | A237V (GCC→GTC) | *nrdB* ← | ribonucleotide reductase class Ia beta subunit |
| 139,330 | A→G |  |  | 10.1% |  |  |  | 5.7% |  |  | V136A (GTG→GCG) | *nrdB* ← | ribonucleotide reductase class Ia beta subunit |
| 139,366 | T→C | 17.1% |  |  |  |  |  |  |  |  | H124R (CAT→CGT) | *nrdB* ← | ribonucleotide reductase class Ia beta subunit |
| 139,472 | G→T |  |  |  |  |  |  | 33.4% | 13.3% |  | R89S (CGT→AGT) | *nrdB* ← | ribonucleotide reductase class Ia beta subunit |
| 139,660 | G→A | 6.7% | 5.6% | 6.8% |  |  |  |  | 11.2% |  | A26V (GCT→GTT) | *nrdB* ← | ribonucleotide reductase class Ia beta subunit |
| 140,747 | A→T |  |  |  |  |  |  |  | 23.4% |  | V650E (GTG→GAG) | *nrdA* ← | NrdA‑like aerobic NDP reductase large subunit |
| 141,315 | C→T |  |  |  |  |  |  |  | 7.5% |  | A461T (GCA→ACA) | *nrdA* ← | NrdA‑like aerobic NDP reductase large subunit |
| 141,395 | C→T | 7.9% |  |  |  |  |  |  |  |  | S434N (AGT→AAT) | *nrdA* ← | NrdA‑like aerobic NDP reductase large subunit |
| 141,830 | C→A |  |  |  |  |  |  |  |  | 9.6% | C289F (TGT→TTT) | *nrdA* ← | NrdA‑like aerobic NDP reductase large subunit |
| 141,951 | C→T |  |  |  |  |  |  | 9.4% |  | 5.2% | A249T (GCT→ACT) | *nrdA* ← | NrdA‑like aerobic NDP reductase large subunit |
| 141,953 | C→T |  |  |  |  |  |  | 25.0% |  | 27.0% | R248H (CGC→CAC) | *nrdA* ← | NrdA‑like aerobic NDP reductase large subunit |
| 142,083 | G→A | 8.5% |  |  |  |  |  |  |  | 24.8% | P205S (CCA→TCA) | *nrdA* ← | NrdA‑like aerobic NDP reductase large subunit |
| 142,103 | G→A |  |  |  |  | 82.7% |  |  |  |  | S198F (TCT→TTT) | *nrdA* ← | NrdA‑like aerobic NDP reductase large subunit |
| 142,160 | T→C |  | 17.9% |  |  |  |  |  | 25.2% |  | H179R (CAT→CGT) | *nrdA* ← | NrdA‑like aerobic NDP reductase large subunit |
| 142,808 | G→A |  | 8.9% |  |  |  |  |  |  |  | P69S (CCT→TCT) | *nrdA.1* ← | hypothetical protein |
| 144,601 | Δ9 bp |  |  |  | 9.7% |  |  |  |  |  | coding (518‑526/549 nt) | *td* ← | thymidylate synthase |
| 144,792 | T→A |  | 7.1% |  |  |  |  |  |  |  | Q112L (CAG→CTG) | *td* ← | thymidylate synthase |
| 144,864 | T→A |  |  |  |  |  |  |  | 6.9% |  | D88V (GAT→GTT) | *td* ← | thymidylate synthase |
| 147,929 | C→G |  |  |  |  |  | 5.4% |  |  |  | Q209H (CAG→CAC) | *SegG* ← | homing endonuclease |
| 151,474 | C→T | 16.8% | 26.2% | 59.5% | 47.7% | 27.7% | 41.7% | 37.1% | 20.9% | 22.6% | R218C (CGT→TGT) | *34* → | tail fiber protein proximal subunit |
| 151,657 | A→T |  |  | 33.2% |  |  |  |  |  |  | I279F (ATT→TTT) | *34* → | tail fiber protein proximal subunit |
| 151,730 | T→C |  |  |  |  | 6.7% |  |  |  |  | L303S (TTA→TCA) | *34* → | tail fiber protein proximal subunit |
| 151,744 | A→G |  |  | 11.1% |  |  |  | 9.5% |  |  | T308A (ACT→GCT) | *34* → | tail fiber protein proximal subunit |
| 151,777 | A→T | 77.7% |  |  |  |  |  |  |  |  | M319L (ATG→TTG) | *34* → | tail fiber protein proximal subunit |
| 151,797 | C→A |  |  |  |  | 5.7% | 9.1% |  |  |  | N325K (AAC→AAA) | *34* → | tail fiber protein proximal subunit |
| 151,810 | +AAA |  |  |  |  |  | 37.3% |  |  |  | coding (988/3870 nt) | *34* → | tail fiber protein proximal subunit |
| 151,853 | C→T |  |  |  | 36.5% | 15.4% | 21.5% |  |  |  | T344I (ACT→ATT) | *34* → | tail fiber protein proximal subunit |
| 151,888 | C→A |  |  |  | 13.5% |  |  |  |  |  | H356N (CAC→AAC) | *34* → | tail fiber protein proximal subunit |
| 151,928 | C→T |  | 13.1% |  |  |  |  |  |  |  | A369V (GCT→GTT) | *34* → | tail fiber protein proximal subunit |
| 151,973 | C→T |  | 17.6% |  |  |  |  |  |  |  | P384L (CCA→CTA) | *34* → | tail fiber protein proximal subunit |
| 151,975 | C→T |  |  |  |  | 6.1% |  |  |  |  | P385S (CCT→TCT) ‡ | *34* → | tail fiber protein proximal subunit |
| 151,976 | C→T |  |  |  |  | 52.6% |  |  |  |  | P385L (CCT→CTT) ‡ | *34* → | tail fiber protein proximal subunit |
| 152,758 | G→A |  |  |  | 12.8% |  |  |  |  |  | G646R (GGA→AGA) | *34* → | tail fiber protein proximal subunit |
| 156,701 | A→T | 17.9% |  |  |  |  |  |  |  |  | D49V (GAT→GTT) | *37* → | long tail fiber protein distal subunit |
| 156,713 | C→T | 18.8% |  |  |  |  |  |  |  |  | A53V (GCT→GTT) | *37* → | long tail fiber protein distal subunit |
| 156,807 | Δ12 bp |  |  | 16.3% |  |  |  |  |  |  | coding (252‑263/3081 nt) | *37* → | long tail fiber protein distal subunit |
| 157,349 | A→G |  |  |  | 12.3% |  |  |  |  |  | D265G (GAT→GGT) | *37* → | long tail fiber protein distal subunit |
| 160,149 | A→G |  |  |  |  |  |  |  |  | 5.3% | E162E (GAA→GAG) | *38* → | tail fiber assembly |
| 160,572 | G→A |  |  |  | 24.7% | 20.3% | 16.5% |  |  |  | D113N (GAT→AAT) ‡ | *t* → | holin |
| 160,573 | A→G |  |  |  | 12.3% | 7.1% | 5.4% |  |  |  | D113G (GAT→GGT) ‡ | *t* → | holin |
| 160,752 | T→C |  |  |  | 13.7% |  | 10.5% |  |  |  | Y173H (TAC→CAC) | *t* → | holin |
| 161,221 | G→A |  |  | 7.9% |  |  |  |  |  |  | T37I (ACA→ATA) | *asiA.1* ← | AsiA.1 hypothetical protein |
