## Supplementary material for "Osmotic conditions shape fitness gains and resistance mechanisms during *E. coli* and T4 phage co-evolution": Table S4: Table_S4_CoPhage.html

Mutation Comparison


| Predicted mutations | | | | | | | | | | | | | |
| --- | --- | --- | --- | --- | --- | --- | --- | --- | --- | --- | --- | --- | --- |
| position | mutation | L1_36_ | L2_9_ | L3_13_ | M1_6_ | M2_6_ | M3_36_ | H1_36_ | H2_36_ | H3_36_ | annotation | gene | description |
| 77 | Δ12 bp |  |  |  |  |  |  |  |  | 6.4% | pseudogene (2102‑2113/2178 nt) | *rIIA* ← | RIIA lysis inhibitor |
| 572:1 | +C |  |  |  |  |  |  |  |  | 5.7% | pseudogene (1618/2178 nt) | *rIIA* ← | RIIA lysis inhibitor |
| 2,841 | G→A |  |  |  |  |  |  |  |  | 31.8% | intergenic (‑39/+12) | *60* ← / ← *60* | DNA topoisomerase II/DNA topoisomerase II |
| 2,908 | G→A |  |  |  | 94.4% |  |  |  |  |  | S28F (TCC→TTC) | *60* ← | DNA topoisomerase II |
| 3,077 | A→G |  |  |  |  |  |  |  |  | 8.3% | I92T (ATA→ACA) | *60.1* ← | gp60.1 hypothetical protein |
| 4,235 | T→C |  |  |  |  |  |  | 13.5% |  |  | pseudogene (1083/1551 nt) | *39* ← | DNA topoisomerase II large subunit |
| 6,662 | Δ1 bp |  |  |  |  |  | 100% |  |  |  | coding (467/489 nt) | *motB* ← | MotB‑like transcriptional regulator |
| 7,269 | T→C |  |  |  |  |  |  |  |  | 26.5% | intergenic (‑141/+9) | *motB* ← / ← *motB.1* | MotB‑like transcriptional regulator/hypothetical protein |
| 9,522 | C→T |  |  |  |  |  | 19.6% |  |  |  | A398A (GCG→GCA) | *dda* ← | Dda‑like helicase |
| 14,150 | T→C |  |  |  |  |  |  |  |  | 27.0% | Y18C (TAT→TGT) | *srh* ← | decoy of host sigma32 |
| 17,409 | T→C |  |  |  |  |  |  |  |  | 85.6% | D68G (GAT→GGT) | *dam* ← | Dam family site‑specific DNA‑(adenine‑N6)‑methyl transferase |
| 19,470 | T→C |  |  |  |  |  |  |  |  | 12.6% | D92G (GAC→GGC) | *61.2* ← | gp61.2 hypothetical protein |
| 20,039 | A→G |  |  |  |  |  |  |  |  | 63.1% | intergenic (‑2/+59) | *sp* ← / ← *61.4* | spackle periplasmic/gp61.4 hypothetical protein |
| 24,513 | A→G |  |  |  |  |  |  |  |  | 17.1% | V310A (GTC→GCC) | *b‑gt* ← | beta‑glucosyltransferase |
| 26,499 | T→C |  |  |  |  |  |  |  |  | 42.2% | T38A (ACA→GCA) | *imm* ← | immunity to superinfection |
| 26,563 | G→A |  |  |  |  |  |  |  |  | 5.2% | G16G (GGC→GGT) | *imm* ← | immunity to superinfection |
| 27,983 | C→G |  |  |  |  |  |  |  | 93.5% |  | V633L (GTT→CTT) | *43* ← | DNA polymerase |
| 29,921 | T→C |  |  |  |  |  |  |  | 5.0% |  | intergenic (‑42/+37) | *43* ← / ← *regA* | DNA polymerase/translation repressor |
| 30,219 | A→G |  |  |  |  |  |  |  |  | 5.7% | C36C (TGT→TGC) | *regA* ← | translation repressor |
| 33,093 | T→C |  |  |  |  |  |  | 7.9% |  |  | I47M (ATA→ATG) | *45.2* ← | protein GP45.2 |
| 34,220 | T→C |  |  |  |  |  |  |  |  | 82.4% | D251G (GAC→GGC) | *46* ← | SbcC‑like subunit of palindrome specific endonuc lease |
| 38,594 | T→C |  |  |  |  |  |  |  |  | 33.5% | T25A (ACA→GCA) | *mobB* ← | homing endonuclease |
| 43,839 | T→C |  |  |  |  |  |  |  |  | 100% | T501A (ACA→GCA) | *nrdD* ← | anaerobic ribonucleoside reductase large subunit |
| 44,851 | A→G |  |  |  |  |  |  |  |  | 11.6% | I250T (ATT→ACT) | *I‑TevII* ← | I‑TevII homing endonuclease |
| 45,948 | C→T |  |  |  |  |  | 5.5% |  |  |  | R142H (CGC→CAC) | *nrdD* ← | anaerobic ribonucleoside reductase large subunit |
| 47,321 | G→A |  |  |  |  |  |  |  |  | 91.8% | L17L (CTG→TTG) | *pin* ← | inhibitor of host Lon protease |
| 47,480 | G→A |  |  |  |  |  |  |  |  | 84.0% | A10V (GCG→GTG) | *49.1* ← | gp49.1 conserved protein of unknown function |
| 49,238 | A→G |  |  |  |  |  |  | 8.9% |  |  | H203H (CAT→CAC) | *nrdC.3* ← | hypothetical protein |
| 50,139 | T→C |  |  |  |  |  |  |  |  | 15.5% | D255G (GAT→GGT) | *nrdC.4* ← | hypothetical protein |
| 53,465 | A→G |  |  |  |  |  |  |  |  | 10.8% | S137P (TCA→CCA) | *nrdC.8* ← | hypothetical protein |
| 55,347:1 | +C |  |  |  |  |  |  | 21.6% |  |  | intergenic (‑40/+75) | *nrdC.10* ← / ← *nrdC.11* | hypothetical protein/nucleotidyltransferase |
| 55,347:2 | +C |  |  |  |  |  |  | 21.6% |  |  | intergenic (‑40/+75) | *nrdC.10* ← / ← *nrdC.11* | hypothetical protein/nucleotidyltransferase |
| 55,486 | A→G |  |  |  |  |  |  |  |  | 35.1% | M316T (ATG→ACG) | *nrdC.11* ← | nucleotidyltransferase |
| 57,448:1 | +G |  |  |  |  |  |  | 16.0% |  |  | coding (368/546 nt) | *mobD.1* ← | MobD.1 conserved hypothetical protein |
| 59,403 | A→T |  |  |  |  |  |  |  |  | 43.1% | D27E (GAT→GAA) | *rI* ← | lysis inhibition |
| 59,413 | G→A |  |  |  |  |  |  |  | 100% |  | A24V (GCG→GTG) | *rI* ← | lysis inhibition |
| 59,413 | G→T |  |  |  |  |  |  | 77.3% |  |  | A24E (GCG→GAG) | *rI* ← | lysis inhibition |
| 59,414 | C→T |  |  |  |  |  |  |  |  | 55.0% | A24T (GCG→ACG) | *rI* ← | lysis inhibition |
| 59,599 | A→G |  |  |  |  |  |  | 10.5% |  |  | L37P (CTC→CCC) | *rI.1* ← | rI.1 conserved hypothetical protein |
| 61,417 | T→C |  |  |  |  |  |  |  |  | 41.7% | D102G (GAC→GGC) | *vs* ← | valyl tRNA synthetase modifier |
| 63,300 | A→G |  |  |  |  |  |  |  |  | 5.0% | H11H (CAT→CAC) | *vs.4* ← | Vs.4 conserved hypothetical protein |
| 65,714 | Δ697 bp |  |  |  |  |  |  | 100% |  | 100% |  | *ipIII* | ipIII |
| 72,680 | G→A |  |  |  |  |  |  | 5.2% |  |  | P72L (CCT→CTT) | *trna.2* ← | Trna.2 conserved hypothetical protein |
| 73,500 | Δ365 bp |  |  |  |  |  |  |  | 100% |  |  | *ipI* | ipI |
| 74,220 | A→G |  |  |  |  |  |  |  |  | 20.1% | V57A (GTT→GCT) | *57B* ← | RNA ligase |
| 81,721 | T→A |  |  |  |  |  |  |  |  | 35.5% | S221T (TCA→ACA) | *6* → | baseplate wedge subunit |
| 83,886 | G→T | 100% | 100% | 100% |  |  | 100% |  |  |  | D283Y (GAC→TAC) | *7* → | baseplate wedge subunit |
| 84,039 | G→A |  |  |  |  |  |  | 5.2% |  |  | A334T (GCA→ACA) | *7* → | baseplate wedge subunit |
| 85,508 | T→C |  |  |  |  |  |  |  |  | 34.6% | G823G (GGT→GGC) | *7* → | baseplate wedge subunit |
| 87,440 | T→C |  |  |  |  |  |  | 44.4% |  |  | V81A (GTT→GCT) | *9* → | baseplate wedge tail fiber protein connector |
| 87,995 | G→A |  |  |  |  | 82.6% |  |  |  |  | S266N (AGT→AAT) | *9* → | baseplate wedge tail fiber protein connector |
| 88,478 | T→C |  |  |  |  |  |  |  |  | 43.4% | C138C (TGT→TGC) | *10* → | baseplate wedge subunit |
| 89,712 | 2 bp→AC |  |  |  |  |  |  |  | 100% |  | coding (1648‑1649/1809 nt) | *10* → | baseplate wedge subunit |
| 89,713 | T→C |  |  |  |  | 80.6% |  |  |  |  | V550A (GTC→GCC) | *10* → | baseplate wedge subunit |
| 89,770 | A→G |  |  |  | 100% | 16.8% |  | 100% |  | 57.0% | Y569C (TAC→TGC) | *10* → | baseplate wedge subunit |
| 90,233 | A→G |  |  |  |  |  |  | 44.8% |  |  | T121A (ACG→GCG) | *11* → | baseplate wedge subunit |
| 90,236 | G→A |  |  |  |  |  |  |  |  | 9.9% | A122T (GCA→ACA) | *11* → | baseplate wedge subunit |
| 90,252 | C→T |  |  |  |  |  |  | 20.4% |  |  | A127V (GCA→GTA) | *11* → | baseplate wedge subunit |
| 90,330 | G→A |  |  |  |  |  |  |  |  | 17.4% | G153D (GGT→GAT) | *11* → | baseplate wedge subunit |
| 91,917 | C→T |  |  |  |  |  |  |  |  | 64.0% | pseudogene (1389/1584 nt) | *12* → | tail collar fiber protein |
| 93,304 | +AGC |  |  |  |  |  |  |  |  | 6.6% | coding (1196/1464 nt) | *wac* → | fibritin neck whisker |
| 93,359 | C→T |  |  |  |  |  |  |  |  | 14.5% | R417R (CGC→CGT) | *wac* → | fibritin neck whisker |
| 99,134 | G→A |  |  |  |  |  |  |  |  | 20.4% | pseudogene (619/1980 nt) | *18* → | tail sheath |
| 100,210 | A→G |  |  |  |  |  |  |  |  | 27.4% | pseudogene (1695/1980 nt) | *18* → | tail sheath |
| 105,273 | A→G |  |  |  |  |  |  | 86.5% |  |  | M117V (ATG→GTG) | *23* → | major head protein |
| 105,275 | G→A |  |  |  |  |  | 22.6% |  |  | 100% | M117I (ATG→ATA) | *23* → | major head protein |
| 105,301 | C→T | 100% |  |  |  |  |  |  |  |  | A126V (GCA→GTA) | *23* → | major head protein |
| 105,821 | G→A |  |  |  |  |  |  |  | 100% |  | M299I (ATG→ATA) | *23* → | major head protein |
| 107,882 | C→A |  |  |  |  |  |  | 34.3% |  |  | A194D (GCT→GAT) | *24* → | capsid vertex protein |
| 108,206 | C→T |  |  |  |  |  | 85.2% |  |  |  | A302V (GCC→GTC) | *24* → | capsid vertex protein |
| 110,757 | +ATTC | 8.9% |  |  |  |  |  |  |  |  | coding (573/1131 nt) | *hoc* ← | Hoc‑like head decoration |
| 110,874 | A→G |  |  |  |  |  |  | 17.6% |  |  | T152T (ACT→ACC) | *hoc* ← | Hoc‑like head decoration |
| 111,067 | Δ10 bp | 14.3% |  |  |  |  |  |  |  |  | coding (254‑263/1131 nt) | *hoc* ← | Hoc‑like head decoration |
| 111,209 | +CTAC | 6.8% |  |  |  |  |  |  |  |  | coding (121/1131 nt) | *hoc* ← | Hoc‑like head decoration |
| 117,666 | T→A |  |  |  |  |  |  | 45.8% |  |  | D142E (GAT→GAA) | *27* → | baseplate hub |
| 120,901 | T→C |  |  |  |  |  |  | 9.5% |  |  | L77P (CTT→CCT) | *48* → | baseplate tail tube cap |
| 121,607 | T→C |  |  |  |  |  |  |  |  | 25.2% | G312G (GGT→GGC) | *48* → | baseplate tail tube cap |
| 123,617 | A→G |  |  |  |  |  |  |  |  | 18.1% | V635A (GTT→GCT) | *alt* ← | Alt‑like RNA polymerase ADP‑ribosyltransferase |
| 123,633 | C→T |  |  |  |  |  |  |  |  | 10.7% | E630K (GAA→AAA) | *alt* ← | Alt‑like RNA polymerase ADP‑ribosyltransferase |
| 123,904 | T→C |  |  |  |  |  |  |  |  | 14.8% | G539G (GGA→GGG) | *alt* ← | Alt‑like RNA polymerase ADP‑ribosyltransferase |
| 126,520 | T→C |  |  |  |  |  |  |  |  | 59.5% | D236G (GAT→GGT) | *30* ← | DNA ligase |
| 134,842 | C→T |  |  |  |  |  |  | 10.5% |  |  | G28E (GGG→GAG) | *pseT* ← | polynucleotide kinase |
| 138,196 | T→C |  |  |  |  |  |  |  |  | 14.5% | S315G (AGC→GGC) | *nrdB* ← | ribonucleotide reductase class Ia beta subunit |
| 138,837 | A→T |  |  |  |  |  |  |  |  | 37.9% | L23I (TTA→ATA) | *I‑TevIII* ← | endonuclease |
| 139,607 | T→A |  |  |  |  |  |  |  | 100% |  | I44F (ATC→TTC) | *nrdB* ← | ribonucleotide reductase class Ia beta subunit |
| 140,298 | T→C |  |  |  |  |  |  |  |  | 32.1% | pseudogene (134/425 nt) | *mobE* ← | homing endonuclease |
| 140,951 | T→C |  |  |  |  |  |  |  |  | 22.7% | D582G (GAC→GGC) | *nrdA* ← | NrdA‑like aerobic NDP reductase large subunit |
| 141,236 | T→C |  |  |  |  |  |  |  |  | 20.1% | D487G (GAT→GGT) | *nrdA* ← | NrdA‑like aerobic NDP reductase large subunit |
| 141,879 | T→C |  |  |  |  |  |  |  |  | 40.7% | T273A (ACT→GCT) | *nrdA* ← | NrdA‑like aerobic NDP reductase large subunit |
| 141,953 | C→T |  |  |  |  |  |  | 100% |  |  | R248H (CGC→CAC) | *nrdA* ← | NrdA‑like aerobic NDP reductase large subunit |
| 142,045 | T→C |  |  |  |  |  |  |  |  | 13.1% | R217R (CGA→CGG) | *nrdA* ← | NrdA‑like aerobic NDP reductase large subunit |
| 143,557 | G→T |  |  |  |  |  |  | 5.3% |  |  | P185Q (CCG→CAG) | *td* ← | thymidylate synthase |
| 144,897 | T→C |  |  |  |  |  |  | 15.5% |  |  | H77R (CAC→CGC) | *td* ← | thymidylate synthase |
| 146,134 | T→C |  |  |  |  |  |  |  |  | 35.5% | intergenic (‑116/+23) | *frd.1* ← / ← *frd.2* | DUF5417 domain‑containing protein/hypothetical protein |
| 146,797 | T→A |  |  |  |  |  |  |  |  | 34.0% | D7V (GAC→GTC) | *frd.3* ← | hypothetical protein |
| 151,288 | G→A |  | 80.9% |  |  |  |  |  |  |  | V156I (GTA→ATA) | *34* → | tail fiber protein proximal subunit |
| 151,289 | T→C |  |  |  |  |  |  |  |  | 14.3% | V156A (GTA→GCA) | *34* → | tail fiber protein proximal subunit |
| 151,320 | G→T |  |  |  |  |  |  | 24.5% |  |  | Q166H (CAG→CAT) | *34* → | tail fiber protein proximal subunit |
| 151,617 | A→G |  |  |  |  |  |  | 18.2% |  |  | V265V (GTA→GTG) | *34* → | tail fiber protein proximal subunit |
| 152,189 | T→A |  |  |  |  |  |  |  |  | 60.4% | V456D (GTC→GAC) | *34* → | tail fiber protein proximal subunit |
| 152,688 | A→G |  |  |  |  |  |  |  |  | 32.3% | G622G (GGA→GGG) | *34* → | tail fiber protein proximal subunit |
| 156,298 | T→C |  |  |  |  |  |  |  |  | 44.5% | G139G (GGT→GGC) | *36* → | hinge connector of long tail fiber protein dista l connector |
| 159,337 | G→A |  |  |  |  |  | 100% |  |  |  | E928K (GAG→AAG) | *37* → | long tail fiber protein distal subunit |
| 159,345 | C→A | 93.9% |  |  |  |  |  |  |  |  | S930R (AGC→AGA) | *37* → | long tail fiber protein distal subunit |
| 159,373 | G→C | 100% |  |  |  |  |  |  |  |  | G940R (GGT→CGT) | *37* → | long tail fiber protein distal subunit |
| 159,377 | T→C | 100% |  |  |  |  |  |  |  |  | V941A (GTA→GCA) | *37* → | long tail fiber protein distal subunit |
| 159,385 | A→T |  |  |  |  |  |  |  |  | 67.7% | N944Y (AAT→TAT) ‡ | *37* → | long tail fiber protein distal subunit |
| 159,386 | A→T |  |  |  |  |  |  | 100% |  | 68.3% | N944I (AAT→ATT) ‡ | *37* → | long tail fiber protein distal subunit |
| 159,403 | G→A |  |  |  |  |  |  | 8.8% |  |  | A950T (GCC→ACC) | *37* → | long tail fiber protein distal subunit |
| 159,412 | T→C |  |  |  |  | 92.2% |  | 93.6% |  |  | Y953H (TAC→CAC) ‡ | *37* → | long tail fiber protein distal subunit |
| 159,412 | 2 bp→CG |  |  | 100% |  |  | 100% |  |  | 100% | coding (2857‑2858/3081 nt) | *37* → | long tail fiber protein distal subunit |
| 159,413 | A→G |  |  |  |  | 62.4% |  | 93.6% |  |  | Y953C (TAC→TGC) ‡ | *37* → | long tail fiber protein distal subunit |
| 159,418 | G→A |  |  |  | 100% |  |  |  |  |  | A955T (GCG→ACG) | *37* → | long tail fiber protein distal subunit |
| 159,418 | G→C |  |  |  |  |  |  | 85.5% |  |  | A955P (GCG→CCG) | *37* → | long tail fiber protein distal subunit |
| 161,361 | A→G |  |  |  |  |  |  |  |  | 14.4% | V82A (GTC→GCC) | *arn* ← | inhibitor of MrcBC restriction |
| 161,927 | A→G |  |  |  |  |  |  |  |  | 8.7% | R87R (CGT→CGC) | *arn.2* ← | hypothetical protein |
| 163,598 | A→G |  |  |  |  |  |  |  |  | 61.7% | I7T (ATC→ACC) | *motA* ← | MotA‑like activator of middle period transcripti on |
| 163,664 | A→G |  |  |  |  |  |  | 60.0% |  |  | intergenic (‑47/+81) | *motA* ← / ← *motA.1* | MotA‑like activator of middle period transcripti on/MotA.1 hypothetical predicted periplasmic protei n |
| 163,666:1 | +C |  |  |  |  |  |  |  |  | 100% | intergenic (‑49/+79) | *motA* ← / ← *motA.1* | MotA‑like activator of middle period transcripti on/MotA.1 hypothetical predicted periplasmic protei n |
| 163,667 | Δ1 bp |  |  |  |  |  |  |  | 100% |  | intergenic (‑50/+78) | *motA* ← / ← *motA.1* | MotA‑like activator of middle period transcripti on/MotA.1 hypothetical predicted periplasmic protei n |
| 163,667 | A→C |  |  |  |  |  |  | 11.8% | Δ |  | intergenic (‑50/+78) | *motA* ← / ← *motA.1* | MotA‑like activator of middle period transcripti on/MotA.1 hypothetical predicted periplasmic protei n |
| 163,670 | G→T |  |  |  |  |  |  | 13.0% |  |  | intergenic (‑53/+75) | *motA* ← / ← *motA.1* | MotA‑like activator of middle period transcripti on/MotA.1 hypothetical predicted periplasmic protei n |
| 163,671 | T→A |  |  |  |  |  |  | 13.0% |  |  | intergenic (‑54/+74) | *motA* ← / ← *motA.1* | MotA‑like activator of middle period transcripti on/MotA.1 hypothetical predicted periplasmic protei n |
| 163,673 | T→A |  |  |  |  |  |  | 11.8% |  |  | intergenic (‑56/+72) | *motA* ← / ← *motA.1* | MotA‑like activator of middle period transcripti on/MotA.1 hypothetical predicted periplasmic protei n |
| 163,677:2 | +A |  |  |  |  |  |  | 12.3% |  |  | intergenic (‑60/+68) | *motA* ← / ← *motA.1* | MotA‑like activator of middle period transcripti on/MotA.1 hypothetical predicted periplasmic protei n |
| 163,680 | G→C |  |  |  |  |  |  | 12.4% |  |  | intergenic (‑63/+65) | *motA* ← / ← *motA.1* | MotA‑like activator of middle period transcripti on/MotA.1 hypothetical predicted periplasmic protei n |
| 166,033 | T→C |  |  |  |  |  |  |  |  | 9.4% | D8G (GAT→GGT) | *ndd* ← | Ndd‑like nucleoid disruption protein |
| 167,183 | T→C |  |  |  |  |  |  |  |  | 5.7% | D165G (GAC→GGC) | *denB* ← | DenB‑like DNA endonuclease IV |
| 168,314 | A→T |  |  |  |  |  |  |  | 100% |  | I203N (ATT→AAT) | *rIIB* ← | RIIB lysis inhibitor |
