## Supplementary material for "Osmotic conditions shape fitness gains and resistance mechanisms during *E. coli* and T4 phage co-evolution": Table S5: Table_S5_CoBacIsolates.html

Mutation Comparison


| Predicted mutations | | | | | | | | | | | | | | | | | | | | | | | | | | | | | | | | | | | | | | | | | | | | | | | | | |
| --- | --- | --- | --- | --- | --- | --- | --- | --- | --- | --- | --- | --- | --- | --- | --- | --- | --- | --- | --- | --- | --- | --- | --- | --- | --- | --- | --- | --- | --- | --- | --- | --- | --- | --- | --- | --- | --- | --- | --- | --- | --- | --- | --- | --- | --- | --- | --- | --- | --- |
| position | mutation | L1-1 | L1-2 | L1-3 | L1-4 | L1-5 | L2-1 | L2-2 | L2-3 | L2-4 | L2-5 | L3-1 | L3-2 | L3-3 | L3-4 | L3-5 | M1-1 | M1-2 | M1-3 | M1-4 | M1-5 | M2-1 | M2-2 | M2-3 | M2-4 | M2-5 | M3-1 | M3-2 | M3-3 | M3-4 | M3-5 | H1-1 | H1-2 | H1-3 | H1-4 | H1-5 | H2-1 | H2-2 | H2-3 | H2-4 | H2-5 | H3-1 | H3-2 | H3-3 | H3-4 | H3-5 | annotation | gene | description |
| 476,255 | IS*5* (–) +4 bp |  |  |  |  |  |  |  |  |  |  |  |  |  |  |  |  |  |  |  |  |  |  |  |  |  |  |  |  |  |  |  | 100% |  |  |  |  |  |  |  |  |  |  |  |  |  | intergenic (‑91/+452) | *tomB* ← / ← *acrB* | Hha toxicity attenuator; conjugation‑related pro tein/multidrug efflux system protein |
| 569,677 | IS*1* (+) +9 bp |  |  |  |  |  |  |  |  |  |  |  |  |  |  |  |  |  |  |  |  |  |  |  | 100% |  |  |  |  |  |  |  |  |  |  |  |  |  |  |  |  |  |  |  |  |  | coding (266‑274/384 nt) | *quuD* → | DLP12 prophage; putative antitermination protein |
| 571,242 | Δ9,220 bp |  |  |  |  |  |  |  |  |  |  |  |  |  |  |  |  |  |  |  |  |  |  |  |  |  |  | 100% |  | 100% |  |  |  |  |  |  |  |  |  |  |  |  |  |  |  |  | IS*5*‑mediated | *[nmpC]*–*[ompT]* | **17 genes** *[nmpC]*, *essD*, *rrrD*, *rzpD*, *rzoD*, *borD*, *ybcV*, *ybcW*, *ylcI*, *nohD*, *aaaD*, *tfaD*, *ybcY*, *ybcY*, *tfaX*, *appY*, *[ompT]* *[nmpC]*, *essD*, *rrrD*, *rzpD*, *rzoD*, *borD*, *ybcV*, *ybcW*, *ylcI*, *nohD*, *aaaD*, *tfaD*, *ybcY*, *ybcY*, *tfaX*, *appY*, *[ompT]* |
| 697,794 | (T)_7→6_ |  |  |  |  |  |  |  |  |  |  |  |  |  |  |  |  |  |  |  |  |  |  |  |  |  |  |  |  |  |  |  |  |  |  |  |  |  |  | 100% |  |  |  |  |  |  | coding (414/1149 nt) | *nagA* ← | N‑acetylglucosamine‑6‑phosphate deacetylase |
| 825,976 | C→T |  |  |  |  |  |  |  |  |  |  |  |  |  |  |  |  |  |  |  |  |  |  |  |  |  |  |  |  |  |  |  |  |  |  |  |  |  |  | 100% |  |  |  |  |  |  | G42R (GGG→AGG) | *ybiH* ← | DUF1956 domain‑containing tetR family putative t ranscriptional regulator |
| 1,014,626 | A→C |  |  |  |  |  |  |  |  |  |  |  |  |  |  |  |  |  |  |  |  |  |  |  |  |  |  |  |  |  | 100% |  |  |  |  |  |  |  |  |  |  |  |  |  |  |  | I295S (ATC→AGC) | *ompA* ← | outer membrane protein A (3a;II\*;G;d) |
| 1,054,515 | A→C |  |  |  |  |  |  |  |  |  |  |  |  |  |  |  |  |  |  |  |  |  |  |  |  |  |  |  |  |  |  |  |  |  |  |  |  |  |  |  | 100% |  |  |  |  |  | N326H (AAC→CAC) | *torC* → | trimethylamine N‑oxide (TMAO) reductase I, cytoc hrome c‑type subunit |
| 1,083,035 | IS*1* (+) +8 bp |  |  |  | 100% |  |  |  |  |  |  |  |  |  |  |  |  |  |  |  |  |  |  |  |  |  |  |  |  |  |  |  |  |  |  |  |  |  |  |  |  |  |  |  |  |  | coding (261‑268/1326 nt) | *pgaC* ← | biofilm PGA synthase PgaCD, catalytic subunit; p oly‑beta‑1,6‑N‑acetyl‑D‑glucosamine synthase; c‑di‑GMP‑sti mulated activity and dimerization |
| 1,160,853 | G→A |  |  |  |  |  |  | 100% |  |  |  |  |  |  |  |  |  |  |  |  |  |  |  |  |  |  |  |  |  |  |  |  |  |  |  |  |  |  |  |  |  |  |  |  |  |  | Q85Q (CAG→CAA) | *ycfP* → | putative UPF0227 family esterase |
| 1,187,999 | T→A |  |  |  |  |  |  |  |  |  |  |  |  |  |  |  |  |  |  |  | 100% |  |  |  |  |  |  |  |  |  |  |  |  |  |  |  |  |  |  |  |  |  |  |  |  |  | H30L (CAT→CTT) | *hflD* ← | putative lysogenization regulator |
| 1,287,416 | IS*1* (+) +9 bp |  |  |  |  |  |  |  |  |  |  |  |  |  |  |  |  |  |  |  |  |  |  |  | 100% |  |  |  |  |  |  |  |  |  |  |  |  |  |  |  |  |  |  |  |  |  | coding (504‑512/909 nt) | *galU* → | glucose‑1‑phosphate uridylyltransferase |
| 1,329,943 | Δ1 bp |  |  |  |  |  |  |  |  |  |  |  | 100% |  |  |  |  |  |  |  |  |  |  |  |  |  |  |  |  |  |  |  |  |  |  |  |  |  |  |  |  |  |  |  |  |  | intergenic (+228/‑145) | *yciX* → / → *acnA* | uncharacterized protein/aconitate hydratase 1 |
| 1,371,590 | G→T |  |  |  |  |  |  |  |  |  |  |  |  |  |  |  | 100% |  |  |  |  |  |  |  |  |  |  |  |  |  |  |  |  |  |  |  |  |  |  |  |  |  |  |  |  |  | G168C (GGC→TGC) | *ycjS* → | putative NADH‑binding oxidoreductase |
| 1,781,190 | IS*1* (+) +8 bp |  |  |  |  |  |  |  |  |  |  |  |  |  |  |  | 100% |  |  |  |  |  |  |  |  |  |  |  |  |  |  |  |  |  |  |  |  |  |  |  |  |  |  |  |  |  | coding (173‑180/2379 nt) | *ppsA* ← | phosphoenolpyruvate synthase |
| 1,817,101 | G→A |  |  |  |  |  |  |  |  |  |  |  |  |  |  | 100% |  |  |  |  |  |  |  |  |  |  |  |  |  |  |  |  |  |  |  |  |  |  |  |  |  |  |  |  |  |  | L129L (CTG→CTA) | *nadE* → | NAD synthetase, NH3/glutamine‑dependent |
| 1,822,277 | A→C |  |  |  |  |  |  |  |  |  |  |  |  |  | 100% |  |  |  |  |  |  |  |  |  |  |  |  |  |  |  |  |  |  |  |  |  |  |  |  |  |  |  |  |  |  |  | G80G (GGT→GGG) | *astB* ← | succinylarginine dihydrolase |
| 2,017,304 | C→T |  |  |  |  |  |  |  |  |  |  |  |  |  |  |  |  |  |  |  |  |  |  |  |  |  | 100% |  | 100% |  | 100% | 100% | 100% |  |  | 100% |  |  |  |  |  | 100% | 100% | 100% | 100% | 100% | intergenic (+145/‑145) | *fliR* → / → *rcsA* | flagellar export pore protein/transcriptional regulator of colanic acid capsul ar biosynthesis |
| 2,017,308 | T→C |  |  |  |  |  |  |  |  |  |  |  |  |  |  |  |  |  |  |  |  |  |  |  |  |  |  |  |  |  |  |  |  |  |  |  | 100% | 100% | 100% | 100% | 100% |  |  |  |  |  | intergenic (+149/‑141) | *fliR* → / → *rcsA* | flagellar export pore protein/transcriptional regulator of colanic acid capsul ar biosynthesis |
| 2,017,986 | A→T |  |  |  |  |  |  |  |  |  |  |  |  |  |  |  |  |  |  |  |  |  |  |  |  |  |  |  |  |  |  |  |  | 100% |  |  |  |  |  |  |  |  |  |  |  |  | I180F (ATC→TTC) | *rcsA* → | transcriptional regulator of colanic acid capsul ar biosynthesis |
| 2,283,593 | Δ22,956 bp |  |  |  |  |  |  |  |  |  |  |  |  |  |  |  |  |  |  |  |  |  |  |  |  |  |  |  |  |  |  |  |  |  | 100% |  |  |  |  |  |  |  |  |  |  |  | IS*5*‑mediated | *[yejO]*–*ompC* | **24 genes** *[yejO]*, *narP*, *ccmH*, *dsbE*, *ccmF*, *ccmE*, *ccmD*, *ccmC*, *ccmB*, *ccmA*, *napC*, *napB*, *napH*, *napG*, *napA*, *napD*, *napF*, *eco*, *mqo*, *yojI*, *alkB*, *ada*, *apbE*, *ompC* *[yejO]*, *narP*, *ccmH*, *dsbE*, *ccmF*, *ccmE*, *ccmD*, *ccmC*, *ccmB*, *ccmA*, *napC*, *napB*, *napH*, *napG*, *napA*, *napD*, *napF*, *eco*, *mqo*, *yojI*, *alkB*, *ada*, *apbE*, *ompC* |
| 2,305,126 | T→A |  |  |  |  |  |  |  |  |  |  | 100% |  |  |  | 100% |  |  |  |  |  |  |  |  |  |  |  |  |  |  |  |  |  |  | Δ |  |  |  |  |  |  |  |  |  |  |  | \*368L (TAA→TTA) | *ompC* ← | outer membrane porin protein C |
| 2,305,627 | Δ6 bp |  |  |  |  |  | 100% | 100% | 100% | 100% | 100% |  |  |  |  |  |  |  |  |  |  |  |  |  |  |  |  |  |  |  |  |  |  |  | Δ |  |  |  |  |  |  |  |  |  |  |  | coding (597‑602/1104 nt) | *ompC* ← | outer membrane porin protein C |
| 2,305,643 | Δ69 bp |  |  |  |  |  |  |  |  |  |  |  |  |  |  |  |  |  |  |  |  |  |  |  |  | 100% |  |  |  |  |  |  |  |  | Δ |  |  |  |  |  |  |  |  |  |  |  | coding (518‑586/1104 nt) | *ompC* ← | outer membrane porin protein C |
| 2,305,645 | C→G | 100% | 100% | 100% | 100% | 100% |  |  |  |  |  |  | 100% | 100% | 100% |  | 100% | 100% | 100% | 100% | 100% | 100% | 100% | 100% | 100% | Δ | 100% |  | 100% |  | 100% | 100% | 100% |  | Δ | 100% |  |  |  |  |  |  |  |  |  |  | R195P (CGT→CCT) | *ompC* ← | outer membrane porin protein C |
| 2,305,782 | Δ2 bp |  |  |  |  |  |  |  |  |  |  |  |  |  |  |  |  |  |  |  |  |  |  |  |  |  |  | 100% |  | 100% |  |  |  |  | Δ |  |  |  |  |  |  |  |  |  |  |  | coding (446‑447/1104 nt) | *ompC* ← | outer membrane porin protein C |
| 2,459,789 | T→A |  |  |  | 100% |  |  |  |  |  |  |  |  |  |  |  |  |  |  |  |  |  |  |  |  |  |  |  |  |  |  |  |  |  |  |  |  |  |  |  |  |  |  |  |  |  | noncoding (2/75 nt) | *argW* → | tRNA‑Arg |
| 2,724,312 | T→C | 100% | 100% | 100% | 100% | 100% |  |  |  |  |  |  | 100% | 100% | 100% |  | 100% | 100% | 100% | 100% | 100% | 100% | 100% | 100% | 100% |  | 100% |  | 100% |  | 100% | 100% | 100% |  |  | 100% |  |  |  |  |  |  |  |  |  |  | noncoding (205/1542 nt) | *rrsG* ← | 16S ribosomal RNA of rrnG operon |
| 2,799,794 | G→A |  |  |  |  |  |  |  |  |  |  |  |  |  |  |  |  |  |  |  |  |  |  |  |  |  |  |  |  |  |  | 100% | 100% |  |  | 100% |  |  |  |  |  |  |  |  |  |  | W142\* (TGG→TGA) | *proW* → | glycine betaine transporter subunit |
| 2,860,448 | Δ1 bp | 100% | 100% | 100% | 100% | 100% |  |  |  |  |  |  |  |  |  |  |  |  |  |  |  |  |  |  |  |  |  |  |  |  |  |  |  |  |  |  |  |  |  |  |  |  |  |  |  |  | coding (463/993 nt) | *rpoS* ← | RNA polymerase, sigma S (sigma 38) factor |
| 3,086,401 | G→A |  |  |  |  |  |  |  |  |  |  |  |  |  |  |  |  |  |  |  |  |  |  |  |  |  |  |  | 100% |  |  |  |  |  |  |  |  |  |  |  |  |  |  |  |  |  | G36R (GGG→AGG) | *yqgE* → | uncharacterized protein |
| 3,314,568 | A→C |  |  |  |  |  |  |  |  |  |  |  |  |  |  |  |  |  |  |  |  |  |  |  |  |  |  |  |  |  |  |  |  |  |  | 100% |  |  |  |  |  |  |  |  |  |  | V135V (GTT→GTG) | *yhbX* ← | putative EptAB family phosphoethanolamine transf erase, inner membrane protein |
| 3,403,589 | (A)_6→7_ |  |  |  |  |  |  |  |  |  |  |  |  |  |  |  |  |  |  |  |  |  |  |  |  |  |  |  |  |  |  |  |  |  | 100% |  |  |  |  |  |  |  |  |  |  |  | intergenic (+279/‑50) | *prmA* → / → *dusB* | methyltransferase for 50S ribosomal subunit prot ein L11/tRNA‑dihydrouridine synthase B |
| 3,404,609 | A→C |  |  |  |  |  |  |  |  |  |  |  |  |  |  |  |  |  |  |  |  |  |  |  |  |  |  |  |  |  |  |  |  |  |  |  |  |  |  |  |  |  | 100% |  |  |  | intergenic (+5/‑21) | *dusB* → / → *fis* | tRNA‑dihydrouridine synthase B/global DNA‑binding transcriptional dual regulato r |
| 3,404,861 | G→A |  |  |  |  |  |  |  |  |  |  |  |  |  |  |  |  |  |  |  |  |  |  |  |  |  |  |  |  |  |  |  |  |  |  |  |  |  |  |  |  |  |  | 100% | 100% |  | A78T (GCG→ACG) | *fis* → | global DNA‑binding transcriptional dual regulato r |
| 3,597,636 | C→A |  |  |  |  |  |  |  |  |  |  |  |  |  |  |  |  |  |  |  |  |  |  |  |  |  |  |  |  |  |  |  |  |  |  | 100% |  |  |  |  |  |  |  |  |  |  | intergenic (‑33/‑117) | *ftsY* ← / → *rsmD* | Signal Recognition Particle (SRP) receptor/16S rRNA m(2)G966 methyltransferase, SAM‑depende nt |
| 3,795,449 | IS*5* (+) +4 bp |  |  |  |  |  |  |  |  |  |  |  |  |  |  |  |  |  |  | 100% | 100% |  |  |  |  |  |  |  |  |  |  |  |  |  |  |  |  |  |  |  |  |  |  |  |  |  | coding (967‑970/1020 nt) | *waaR* ← | UDP‑D‑galactose:(glucosyl)lipopolysaccharide‑ al pha‑1,3‑D‑galactosyltransferase |
| 3,795,864 | (A)_6→7_ |  |  |  |  |  |  |  |  |  |  |  |  |  |  |  |  |  |  |  |  |  |  |  |  | 100% |  |  |  |  |  |  |  |  |  |  |  |  |  |  |  |  |  |  |  |  | coding (555/1020 nt) | *waaR* ← | UDP‑D‑galactose:(glucosyl)lipopolysaccharide‑ al pha‑1,3‑D‑galactosyltransferase |
| 3,795,902 | IS*5* (–) +4 bp |  |  |  |  |  |  |  |  |  |  | 100% |  |  |  | 100% |  |  |  |  |  |  |  |  |  |  |  |  |  |  |  |  |  |  |  |  |  |  |  |  |  |  |  |  |  |  | coding (514‑517/1020 nt) | *waaR* ← | UDP‑D‑galactose:(glucosyl)lipopolysaccharide‑ al pha‑1,3‑D‑galactosyltransferase |
| 3,795,902 | IS*5* (+) +4 bp |  |  |  |  |  |  |  |  |  |  |  |  |  |  |  |  |  |  |  |  | 100% | 100% |  |  |  |  |  |  |  |  |  |  |  |  |  |  |  |  |  |  |  |  |  |  |  | coding (514‑517/1020 nt) | *waaR* ← | UDP‑D‑galactose:(glucosyl)lipopolysaccharide‑ al pha‑1,3‑D‑galactosyltransferase |
| 3,796,008 | +C |  |  |  |  |  | 100% | 100% | 100% | 100% | 100% |  |  |  |  |  |  |  |  |  |  |  |  |  |  |  |  |  |  |  |  |  |  |  |  |  |  |  |  |  |  |  |  |  |  |  | coding (411/1020 nt) | *waaR* ← | UDP‑D‑galactose:(glucosyl)lipopolysaccharide‑ al pha‑1,3‑D‑galactosyltransferase |
| 3,796,012 | C→T |  |  |  |  |  |  |  |  |  |  |  |  |  |  |  | 100% |  |  |  |  |  |  |  |  |  |  |  |  |  |  |  |  |  |  |  |  |  |  |  |  |  |  |  |  |  | C136Y (TGT→TAT) | *waaR* ← | UDP‑D‑galactose:(glucosyl)lipopolysaccharide‑ al pha‑1,3‑D‑galactosyltransferase |
| 3,796,093 | G→T |  |  |  |  |  |  |  |  |  |  |  |  |  |  |  |  |  |  |  |  |  |  | 100% |  |  |  |  |  |  |  |  |  |  |  |  |  |  |  |  |  |  |  |  |  |  | A109E (GCA→GAA) | *waaR* ← | UDP‑D‑galactose:(glucosyl)lipopolysaccharide‑ al pha‑1,3‑D‑galactosyltransferase |
| 3,796,114 | IS*5* (–) +4 bp |  |  |  |  |  |  |  |  |  |  |  |  |  |  |  |  | 100% | 100% |  |  |  |  |  |  |  |  |  |  |  |  |  |  |  |  |  |  |  |  |  |  |  |  |  |  |  | coding (302‑305/1020 nt) | *waaR* ← | UDP‑D‑galactose:(glucosyl)lipopolysaccharide‑ al pha‑1,3‑D‑galactosyltransferase |
| 3,796,397 | C→A |  |  |  |  |  |  |  |  |  |  |  |  |  |  |  |  |  |  |  |  |  |  |  |  |  |  | 100% |  | 100% |  |  |  |  |  |  |  |  |  |  |  |  |  |  |  |  | E8\* (GAA→TAA) | *waaR* ← | UDP‑D‑galactose:(glucosyl)lipopolysaccharide‑ al pha‑1,3‑D‑galactosyltransferase |
| 3,799,422 | IS*5* (–) +4 bp |  |  |  |  |  |  |  |  |  |  |  | 100% | 100% | 100% |  |  |  |  |  |  |  |  |  |  |  |  |  |  |  |  |  |  |  |  |  |  |  |  |  |  |  |  |  |  |  | coding (1003‑1006/1125 nt) | *waaG* ← | glucosyltransferase I |
| 3,809,181 | G→A |  |  |  |  | 100% |  |  |  |  |  |  |  |  |  |  |  |  |  |  |  |  |  |  |  |  |  |  |  |  |  |  |  |  |  |  |  |  |  |  |  |  |  |  |  |  | intergenic (‑53/+13) | *pyrE* ← / ← *rph* | orotate phosphoribosyltransferase/ribonuclease PH (defective);enzyme; Degradation of RNA; RNase PH |
| 3,809,219 | Δ82 bp | 100% | 100% |  |  |  |  |  |  |  |  |  |  |  |  |  |  |  |  |  |  |  |  |  |  |  |  |  |  |  |  |  |  |  |  |  |  |  |  |  |  |  |  |  |  |  |  | *[rph]*–*[rph]* | *[rph]*, *[rph]* |
| 4,226,706 | C→T |  |  |  |  |  |  |  |  |  |  |  |  |  |  |  |  |  |  |  |  |  |  |  |  |  |  |  |  |  |  |  |  |  |  |  |  |  |  |  |  |  |  |  |  | 100% | Q173\* (CAG→TAG) | *yjbF* → | extracellular polysaccharide production lipoprot ein |
| 4,227,545 | C→T |  |  |  |  |  |  |  |  |  |  |  |  |  |  |  |  |  |  |  |  |  |  |  |  |  |  |  |  |  | 100% |  |  |  |  |  |  |  |  |  |  |  |  |  |  |  | Q241\* (CAG→TAG) | *yjbG* → | extracellular polysaccharide export OMA protein |
| 4,227,826 | C→T |  |  | 100% |  |  |  |  |  |  |  |  |  |  |  |  |  |  |  |  |  |  |  |  |  |  |  |  |  |  |  |  |  |  |  |  |  |  |  |  |  |  |  |  |  |  | Q89\* (CAG→TAG) | *yjbH* → | DUF940 family extracellular polysaccharide prote in |
| 4,227,938 | +G |  |  |  |  |  |  |  |  |  |  |  |  |  |  |  |  |  |  |  |  |  |  |  |  |  | 100% |  |  |  |  |  |  |  |  |  |  |  |  |  |  |  |  |  |  |  | coding (377/2097 nt) | *yjbH* → | DUF940 family extracellular polysaccharide prote in |
| 4,228,970 | T→C |  |  |  |  |  |  |  |  |  |  |  |  |  |  |  |  |  |  |  |  |  |  |  |  |  |  |  |  |  |  |  |  |  |  |  |  |  |  |  |  | 100% |  |  |  |  | L470P (CTT→CCT) | *yjbH* → | DUF940 family extracellular polysaccharide prote in |
| 4,342,819 | IS*1* (–) +9 bp |  |  |  |  |  |  |  |  |  |  |  |  |  |  |  |  |  |  |  |  |  | 100% |  |  |  |  |  |  |  |  |  |  |  |  |  |  |  |  |  |  |  |  |  |  |  | coding (95‑103/174 nt) | *yjdO* → | uncharacterized protein |
| 4,344,710 | C→A |  |  |  |  |  |  |  |  |  |  |  |  |  |  |  |  |  |  |  |  |  |  |  |  |  |  |  |  |  |  | 100% |  |  |  |  |  |  |  |  |  |  |  |  |  |  | intergenic (‑176/+61) | *lysU* ← / ← *dtpC* | lysine tRNA synthetase, inducible/dipeptide and tripeptide permease |
| 4,484,376 | (A)_7→6_ |  |  |  |  |  |  |  |  |  |  |  |  |  |  |  |  |  |  |  |  |  |  |  |  |  |  |  |  |  |  |  |  |  |  |  | 100% |  |  |  |  |  |  |  |  |  | intergenic (‑153/‑64) | *idnD* ← / → *idnK* | L‑idonate 5‑dehydrogenase, NAD‑binding/D‑gluconate kinase, thermosensitive |
